# StackHPpred: A Stacking-based ensemble learning framework for the identification of peptide hormones using multi-view feature representations

**DOI:** 10.64898/2026.09.16.751134

**Authors:** Jayasree Kirthipati, Mohan Krishna Chemarthi Ravi, Mounica Kirthipati

## Abstract

Peptide hormones are important signaling molecules that regulate diverse physiological processes and have substantial therapeutic relevance. However, their experimental identification can be challenging because of low abundance, limited stability, and complex post-translational processing, highlighting the need for reliable computational approaches. Here, we present StackHPpred, a stacking-based ensemble framework for accurate identification of hormone peptides (HPs) from amino acid sequences. We systematically evaluated 56 feature representations, comprising 35 conventional sequence descriptors and 21 PLM/NLP-based embeddings, across 10 machine-learning classifiers, generating 560 baseline models. Prediction-level correlation analysis was subsequently employed to identify 14 complementary, non-redundant feature representations, which were integrated through feature- and classifier-level stacking strategies using out-of-fold predicted probabilities. The final StackHPpred model, based on the top three feature representations CTDD, KSCTriad, and PTAB, achieved an AUC of 0.9965, MCC of 0.9506, ACC of 0.9752, and F1-score of 0.9749 on the independent dataset. StackHPpred was further evaluated under increasingly imbalanced independent datasets with positive-to-negative ratios ranging from 1:10 to 1:50, demonstrating robust predictive performance under challenging screening conditions. t-SNE visualization and SHAP analysis further showed improved class separation and highlighted the contributions of multiple classifier–feature combinations to the final prediction. Overall, StackHPpred provides a robust and interpretable framework for distinguishing HPs from non-HPs and is freely available as a web server at https://jayasreekirthipati.org/StackHPpred/, with a standalone implementation available at https://github.com/JayasreeKirthipati04/StackHPpred.

**Graphical abstract:** 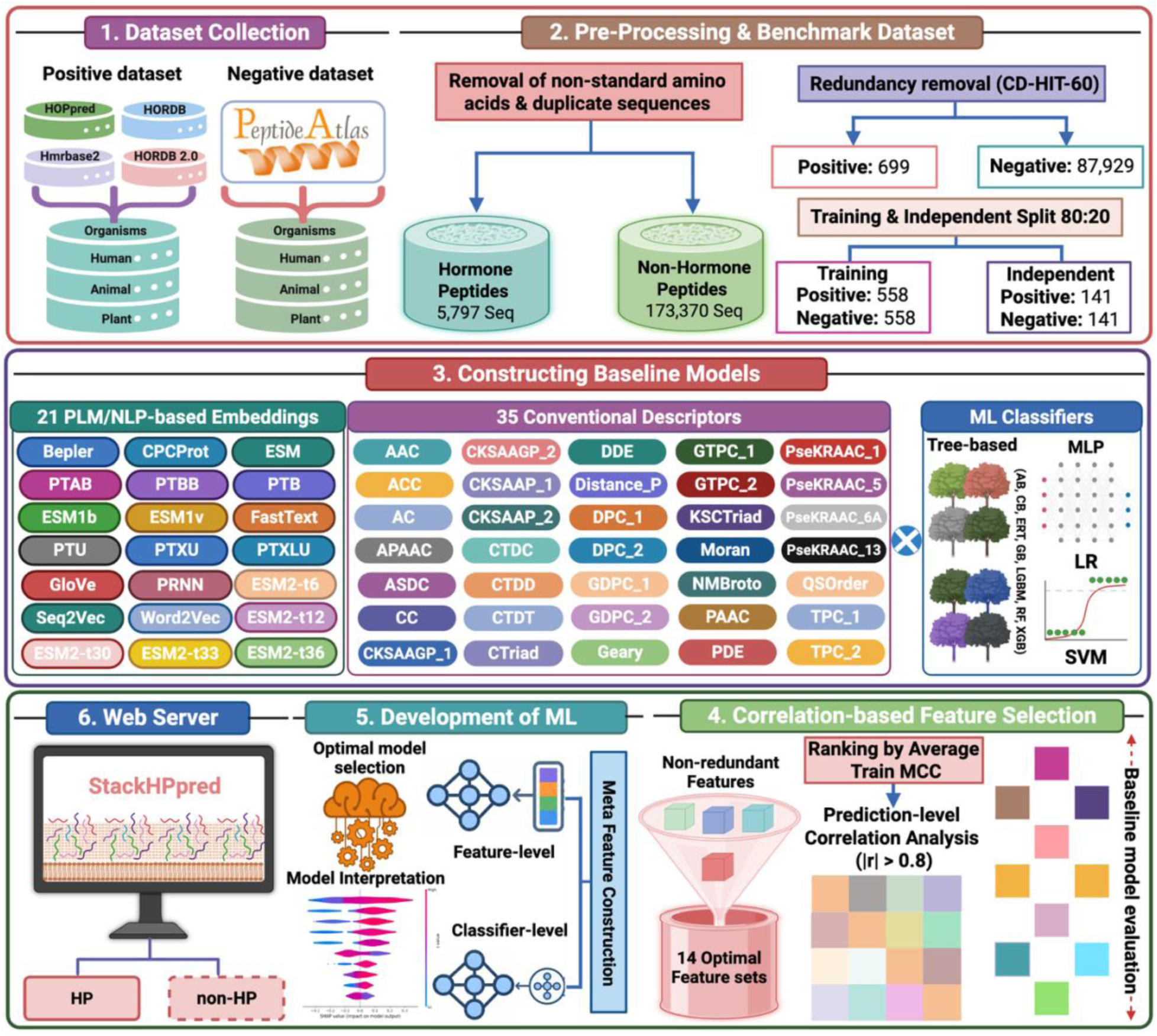

## Introduction

Peptide hormones are important signaling molecules that regulate communication between cells, tissues, and organs and play essential roles in maintaining physiological homeostasis. They are generally synthesized as larger precursor proteins that undergo proteolytic processing and, in many cases, post-translational modifications to generate mature bioactive peptides [1, 2]. By interacting with specific receptors on target cells, peptide hormones regulate diverse physiological processes, including metabolism, growth, appetite, reproduction, stress responses, and cardiovascular and gastrointestinal functions [1-3]. Alterations in their production or signaling can therefore contribute to various endocrine and metabolic disorders [1, 3].

Beyond their essential physiological roles, peptide hormones have provided an important foundation for the development of peptide-based therapeutics. The discovery of insulin established one of the earliest and most successful examples of peptide-based therapy and laid the foundation for modern therapeutic peptide development [4-6]. Since then, several naturally occurring hormones and their analogues, including insulin, glucagon-like peptide-1, oxytocin, and gonadotropin-releasing hormone, have been explored or developed for therapeutic applications [4, 5]. Their high biological activity, target specificity, and generally favorable pharmacological properties make peptide hormones attractive candidates for drug development [4-6].

Despite their biological and therapeutic relevance, experimental identification and characterization of peptide hormones can be challenging because of their low abundance, limited stability, complex processing, and post-translational modifications [1, 2, 7]. Computational approaches can therefore complement experimental methods by enabling rapid identification of potential hormone peptides directly from amino acid sequences. However, computational prediction of peptide hormones remains relatively limited compared with other classes of bioactive peptides. HOPPred was introduced as an ensemble machine-learning approach for peptide hormone prediction [8], while mHPpred subsequently improved prediction performance by integrating multiple sequence representations through a meta-learning framework [9].

Although these methods have improved the computational identification of peptide hormones, there remains scope for enhancing predictive robustness and generalization [8, 9]. Peptide sequences contain diverse information, including amino acid composition, physicochemical properties, local sequence patterns, and contextual representations learned by pretrained protein language models [10, 11]. Effectively identifying complementary information from these representations and integrating their predictive strengths may therefore further improve hormone peptide (HP) prediction. In this study, we developed StackHPpred, a stacking-based ensemble framework for distinguishing HPs from non-HPs. StackHPpred systematically evaluates conventional sequence descriptors and PLM/NLP-based embeddings using multiple machine-learning algorithms and integrates complementary representations through feature- and classifier-level stacking. By leveraging diverse information from multiple feature representations and classifiers within an advanced meta-learning framework, StackHPpred captures informative sequence patterns associated with HPs. The model was evaluated using balanced and increasingly imbalanced independent datasets and implemented as a freely accessible web server and standalone program. Overall, these evaluations support the robustness and practical utility of StackHPpred for peptide hormone prediction.

## Materials and Methods

### Dataset construction and pre-processing

To construct a benchmark dataset for HP prediction, positive samples were compiled from four publicly available databases, including HOPPred [8], HORDB [12], Hmrbase2 [13], and HORDB 2.0 [14], whereas negative samples were collected from PeptideAtlas using corresponding human, animal, and plant sources. Sequences containing non-standard amino acids (B, O, U, X, J, and Z) and duplicate sequences were excluded, resulting in 5,797 HPs and 173,370 non-HPs. It should be noted that high sequence similarity in benchmark datasets can lead to an overestimation of model performance. In previous studies, redundancy reduction was performed separately within the positive and negative classes, an approach that reduces within-class similarity but may retain homologous sequences between the two classes. To address this issue, we combined the positive and negative samples prior to clustering with CD-HIT [15] and applied a sequence identity threshold of 0.60, resulting in 699 HPs and 87,929 non-HPs. The non-redundant positive samples were then randomly partitioned into training and independent test datasets at an 80:20 ratio, after which negative samples were selected from the corresponding non-redundant pool to establish a balanced positive-to-negative ratio of 1:1. Consequently, the training dataset consisted of 558 HPs and 558 non-HPs, whereas the independent test dataset comprised 141 HPs and 141 non-HPs. To further evaluate model robustness under increasingly imbalanced conditions, additional independent test datasets with positive-to-negative ratios of 1:10, 1:20, 1:30, 1:40, and 1:50 were constructed by retaining the same HP samples while progressively increasing the number of non-HP samples.

### Feature extraction

To represent peptide sequences as numerical feature vectors, we employed 21 PLM/NLP-based embeddings and 35 conventional variants derived from 26 descriptor types. The PLM/NLP-based representations included pretrained protein language models and sequence embeddings trained on large-scale protein sequence databases, enabling the extraction of informative representations from peptide and protein sequences. Specifically, we employed Bepler [16], CPCProt [17], ESM, ESM1b [18], ESM1v [19], five ESM2 variants (ESM2-t6, ESM2-t12, ESM2-t30, ESM2-t33, and ESM2-t36) [20], FastText, GloVe, PRNN, ProtTransAlbertBFD (PTAB), ProtTransBertBFD (PTBB), ProtTransT5BFD (PTB), ProtTransT5UniRef50 (PTU), ProtTransT5XLU50 (PTXU), ProtTransXLNetUniRef100 (PTXLU) [21], Seq2Vec [22], and Word2Vec. Several of these protein-sequence embedding models are available through the bio_embeddings framework [23], which provides a standardized interface for generating sequence representations.

The conventional feature descriptors covered diverse composition-, correlation-, sequence-order-, grouped-residue-, and physicochemical-property-based information [24-26]. These included amino acid composition (AAC), auto-cross covariance (ACC), auto covariance (AC), amphiphilic pseudo-amino acid composition (APAAC), adaptive skip dipeptide composition (ASDC), cross covariance (CC), composition of k-spaced amino acid group pairs (CKSAAGP), composition of k-spaced amino acid pairs (CKSAAP), composition (CTDC), distribution (CTDD), and transition (CTDT) descriptors, conjoint triad (CTriad), dipeptide deviation from expected mean (DDE), distance pair (DistancePair), dipeptide composition (DPC), grouped dipeptide composition (GDPC), Geary autocorrelation (Geary), grouped tripeptide composition (GTPC), k-spaced conjoint triad (KSCTriad), Moran autocorrelation (Moran), normalized Moreau–Broto autocorrelation (NMBroto), pseudo-amino acid composition (PAAC), peptide descriptor encoding (PDE) [27], pseudo K-tuple reduced amino acid composition (PseKRAAC), quasi-sequence-order (QSOrder), and tripeptide composition (TPC). For descriptors supporting multiple encoding schemes, Type-1 and Type-2 variants were employed for CKSAAGP, CKSAAP, DPC, GDPC, GTPC, and TPC, whereas Type-4, Type-5, Type-6A, and Type-13 were used for PseKRAAC [28].

### Implementation of ML algorithms

In this study, we employed ten ML classifiers for model development and validation: random forest (RF) [29], extremely randomized trees (ERT) [30], gradient boosting (GB) [31], adaptive boosting (AdaBoost or AB) [32], extreme gradient boosting (XGB) [33], LightGBM (LGB) [34], CatBoost (CB) [35], support vector machine (SVM) [36], logistic regression (LR) [37], and multilayer perceptron (MLP) [38]. RF and ERT are tree-based ensemble methods that construct multiple decision trees, with ERT introducing additional randomness during tree construction to reduce variance and improve generalization. GB, AB, XGB, LGB, and CB are boosting algorithms that sequentially combine weak learners, with successive learners focusing on errors made by preceding models. XGB and LGB provide computationally efficient implementations of gradient boosting, while CB is particularly designed to effectively handle complex feature relationships. SVM identifies a decision boundary that maximizes the margin between classes and, through kernel functions, can capture complex nonlinear relationships in the feature space. LR estimates the probability of binary outcomes using a logistic function, whereas MLP is a neural network architecture capable of learning complex nonlinear relationships. These classifiers have demonstrated strong performance across diverse bioinformatics prediction tasks.

### Construction of baseline models

To construct robust baseline models, we employed 56 feature representations, comprising 21 PLM/NLP-based embeddings and 35 conventional descriptors, each of which was independently combined with 10 ML classifiers. Model performance was evaluated using 10-randomized 10-fold cross-validation (CV) on the training dataset [39]. In each CV iteration, the training data were divided into 10 folds, of which nine were used for model training and the remaining fold was used for validation; this procedure was repeated until each fold had served once as the validation set. Hyperparameters for each classifier were optimized using grid search within the CV procedure [40]. The out-of-fold predictions obtained across the 10 folds were combined to estimate the overall CV performance of each feature–classifier combination. Using the selected hyperparameters, each baseline model was subsequently retrained on the complete training dataset and evaluated on the independent dataset. In total, 560 baseline models (56 feature representations × 10 classifiers) were developed and systematically evaluated.

### Construction of meta-models

To construct the meta-learning models, the 14 non-redundant feature representations retained after correlation-based filtering were ranked according to their average CV MCC. The out-of-fold predicted probabilities generated by the baseline models during CV were used as meta-features to avoid information leakage. Two complementary stacking strategies were explored [41]. In the feature-level strategy, the selected features were progressively integrated from Top-1 to Top-14 according to their average CV MCC. For each feature, the predicted probabilities generated by the 10 classifiers were concatenated, producing meta-feature vectors ranging from 10 dimensions for Top-1 to 140 dimensions for Top-14. These meta-features were subsequently used to train secondary meta-learners. In the classifier-level strategy, the 10 classifiers were ranked according to their average CV MCC, and their predicted probabilities across all 14 selected features were progressively combined from Top-1 to Top-10, resulting in meta-feature vectors ranging from 14 to 140 dimensions. The optimal meta-model configuration was selected solely based on CV MCC, while the independent dataset was reserved for final performance evaluation.

### Performance evaluation metrics

The predictive performance of all developed models was assessed using seven widely adopted classification metrics: Matthews correlation coefficient (MCC) [42], sensitivity (Sn), specificity (Sp), precision (PRE), accuracy (ACC), area under the receiver operating characteristic curve (AUC) [43], and F1-score [44]. These metrics provide complementary assessments of the models’ ability to distinguish HPs from non-HPs. Their mathematical definitions are provided below:

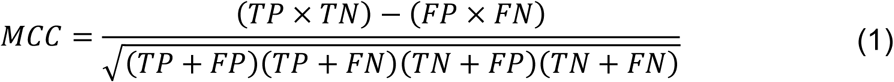

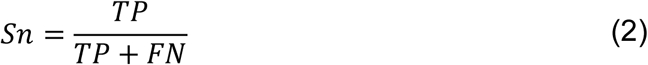

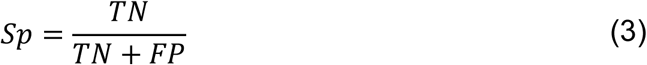

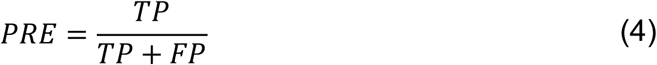

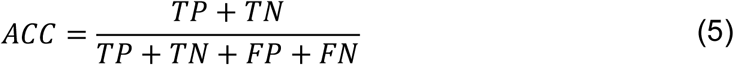

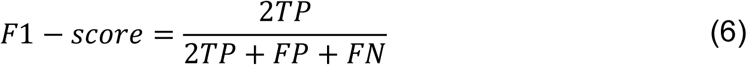

where TP, TN, FP, and FN denote true-positive, true-negative, false-positive, and false-negative predictions, respectively.

## Results and Discussion

### Compositional and Positional Sequence Analysis

Sequence-length distributions of HPs and non-HPs were analyzed using the combined training and independent datasets. The lengths of these peptides ranged from 11 to 41 amino acids, with substantial overlap between the two classes (Fig. S1), suggesting that sequence length alone provides limited discriminatory information. In contrast, amino acid composition analysis revealed clear differences between HPs and non-HPs (Fig. S2; Welch’s *t*-test). Arg showed the strongest enrichment in HPs, accounting for approximately 7.1% of residues compared with 2.8% in non-HPs (*p* = 7.1 × 10⁻⁶⁰). Cys, Phe, Lys, Gly, Met, Asn, Trp, and Tyr were also significantly enriched in HPs, whereas Ala, Glu, Ile, Pro, Gln, Thr, and Val were depleted. Among the depleted residues, Val showed a pronounced difference between HPs and non-HPs (4.3% vs. 6.8%, *p* = 1.6 × 10⁻¹⁷).

Position-specific residue patterns were further investigated using Two Sample Logo (TSL) analysis of the terminal 15 residues (Fig. S3). At the N-terminus, Ser showed strong enrichment at the first position, while Arg and Lys were recurrently enriched across several downstream positions, whereas Val was strongly depleted at the first position. The C-terminal region also showed distinct positional patterns, with Arg and Lys prominently enriched at multiple positions. These observations indicate that HPs differ from non-HPs not only in overall amino acid composition but also in the position-specific distribution of residues at their termini. Together, the compositional and positional differences provide complementary sequence information for distinguishing HPs from non-HPs.

### Performance analysis of conventional descriptors and PLM/NLP-based embeddings in HP prediction using 10 conventional ML algorithms

We systematically evaluated 56 feature representations comprising 35 conventional descriptors and 21 PLM/NLP-based embeddings (Table S1). To assess their ability to distinguish HPs from non-HPs, each representation was paired with 10 different ML algorithms, resulting in a total of 560 single-feature baseline models (Table S2). Their overall predictive performance, measured using MCC, is illustrated in Fig. 2A and 2C for the training dataset and Fig. 2B and 2D for the independent dataset. Among the 560 baseline models, 68 models achieved MCC values exceeding 0.8200. However, substantial performance variation was observed across different classifiers and feature sets. For instance, CKSAAP-type-1-based models achieved MCC values ranging from 0.4696 to 0.8462 across the different classifiers, highlighting substantial intra-feature variability depending on the learning algorithm. The CB model utilizing CTDD demonstrated the best performance among the conventional feature-based models on the training dataset, with AUC, MCC, ACC, Sn, Sp, Precision, and F1-score values of 0.9835, 0.9070, 0.9525, 0.9230, 0.9821, 0.9810, and 0.9509, respectively. The corresponding metrics for this model on the independent dataset were 0.9932, 0.9231, 0.9610, 0.9362, 0.9858, 0.9851, and 0.9600, respectively. Among the PLM/NLP-based embedding models, the LRT model utilizing Seq2Vec demonstrated the best performance on the training dataset, with AUC, MCC, ACC, Sn, Sp, Precision, and F1-score values of 0.9838, 0.8812, 0.9400, 0.9247, 0.9552, 0.9543, and 0.9388, respectively. The corresponding metrics for this model on the independent dataset were 0.9869, 0.9433, 0.9716, 0.9716, 0.9716, 0.9716, and 0.9716, respectively. To evaluate the overall discriminative ability of each feature representation, we averaged training MCC obtained across the 10 ML classifiers. The analysis showed that CTDD, Seq2Vec, DPC-type-2, DPC-type-1, and DDE were the five highest-performing representations, with average MCC values ranging from 0.8026 to 0.8844. Among these, CTDD achieved the highest average MCC of 0.8844, followed by Seq2Vec (0.8304), DPC-type-2 (0.8285), DPC-type-1 (0.8249), and DDE (0.8026). Upon further examination, Seq2Vec was the only NLP-based embedding among the five top-performing representations and captures contextual and biophysical information from protein sequences. The remaining four were conventional descriptors. Specifically, CTDD captures the positional distribution of physicochemical properties within peptides, while DPC-type-1 and DPC-type-2 characterize local dipeptide composition patterns. DDE further measures the deviation of observed dipeptide occurrences from their expected frequencies. These findings indicate that distinguishing HPs from non-HPs is supported by diverse information captured from physicochemical properties, local sequence patterns, and contextual sequence representations. However, high-performing representations may still capture redundant information. Therefore, we next performed Pearson correlation analysis on their predicted probability scores to identify complementary feature representations for StackHPpred development.

**Figure 1.**
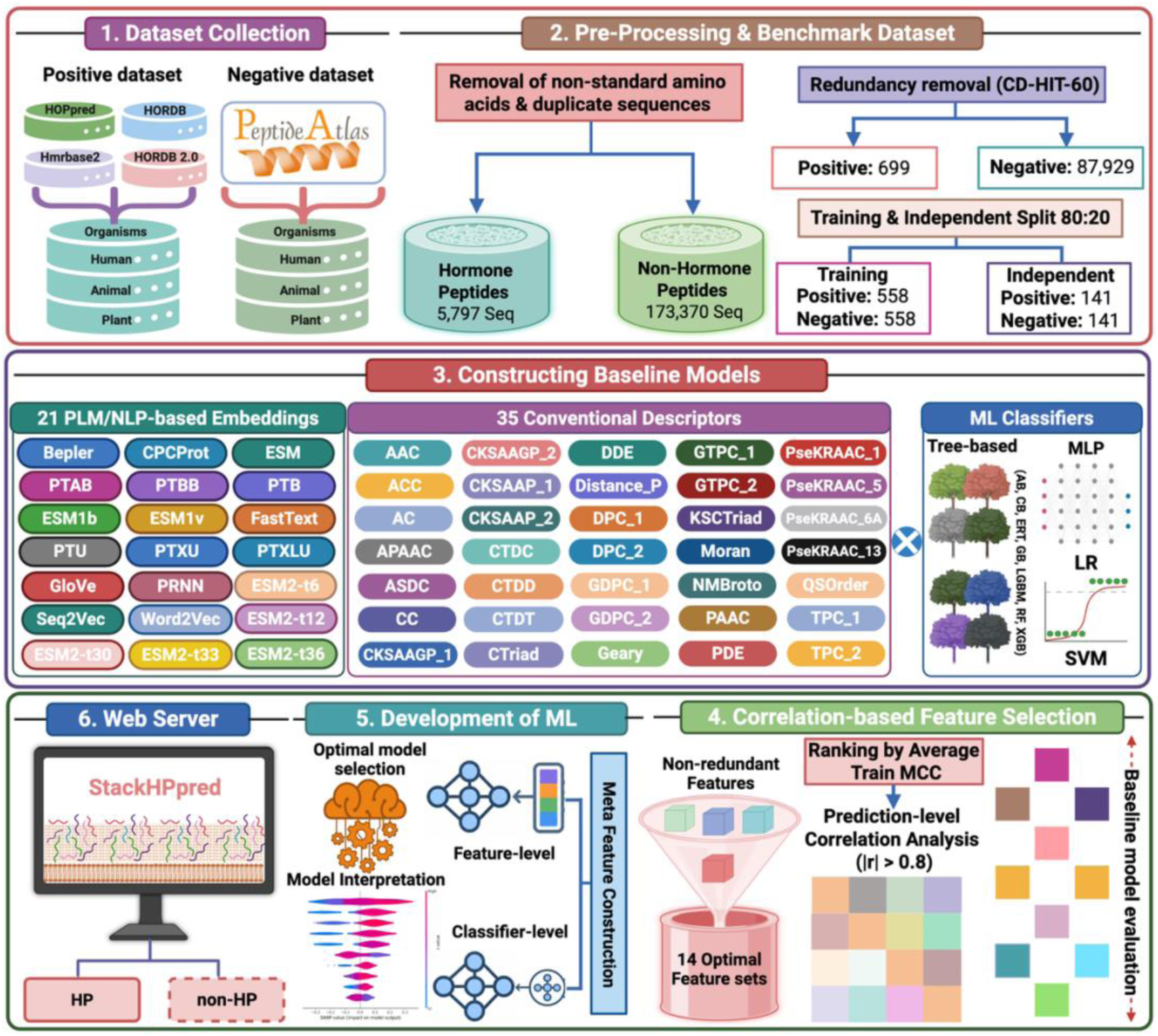
Schematic overview of the StackHPpred development workflow, comprising (i) dataset collection, (ii) sequence pre-processing, redundancy reduction, and benchmark dataset construction, (iii) development of baseline models using 21 PLM/NLP-based embeddings, 35 conventional descriptors, and 10 machine-learning classifiers, (iv) correlation-based selection of 14 complementary feature representations, (v) development of StackHPpred through feature- and classifier-level stacking, optimal model selection, and model interpretation, and (vi) web server development for classification of HPs and non-HPs.

**Figure 2.**
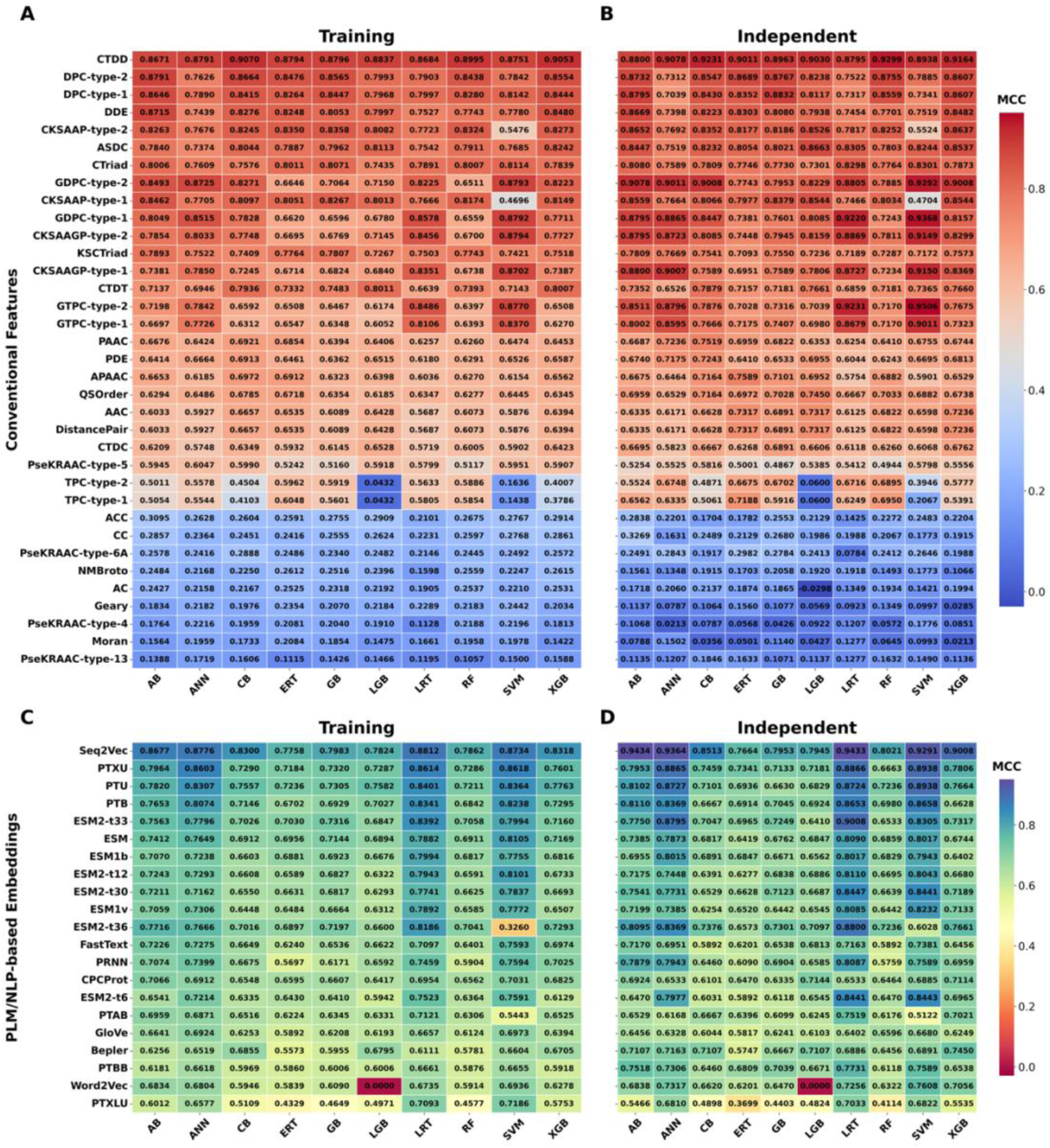
Performance comparison of baseline models using 35 conventional feature descriptors, 21 PLM/NLP-based embeddings, and 10 machine-learning classifiers. Heatmaps show the Matthews correlation coefficient (MCC) of conventional descriptor-based models on the training (A) and independent (B) datasets and PLM/NLP embedding-based models on the training (C) and independent (D) datasets.

### Correlation-based selection of complementary feature representations

Although several feature representations achieved strong individual performance, their prediction patterns may still contain substantial redundancy. We therefore examined pairwise Pearson correlations among the out-of-fold prediction probabilities generated through 10-fold cross-validation. For each feature, the probabilities from the 10 classifiers were averaged and used for the correlation analysis (Fig. S4). The features were ranked from highest to lowest according to their average cross-validation MCC. When two features showed |*r*| > 0.80, their prediction patterns were considered highly redundant, and the feature with the higher average MCC was retained. CTDD achieved the highest average cross-validation MCC of 0.8844, followed by Seq2Vec with 0.8304. However, the prediction probabilities of Seq2Vec showed a strong correlation with those of CTDD (|*r*| = 0.8913), indicating substantial similarity in their predictive behavior. Similar patterns were observed across several other features, resulting in the selection of 14 features with distinct prediction patterns. Of the 21 PLM/NLP-based embeddings, only two were retained, whereas 12 conventional descriptors remained after correlation-based filtering. This observation suggests that the conventional descriptors exhibited greater diversity in their predictive behavior for HP prediction in this study. The final 14 features were CTDD, KSCTriad, PTAB, PseKRAAC-type-5, PTXLU, TPC-type-2, ACC, PseKRAAC-type-6A, NMBroto, AC, Geary, PseKRAAC-type-4, Moran, and PseKRAAC-type-13 (Fig. 3). These selected features were subsequently used for feature- and classifier-level integration in StackHPpred.

**Figure 3.**
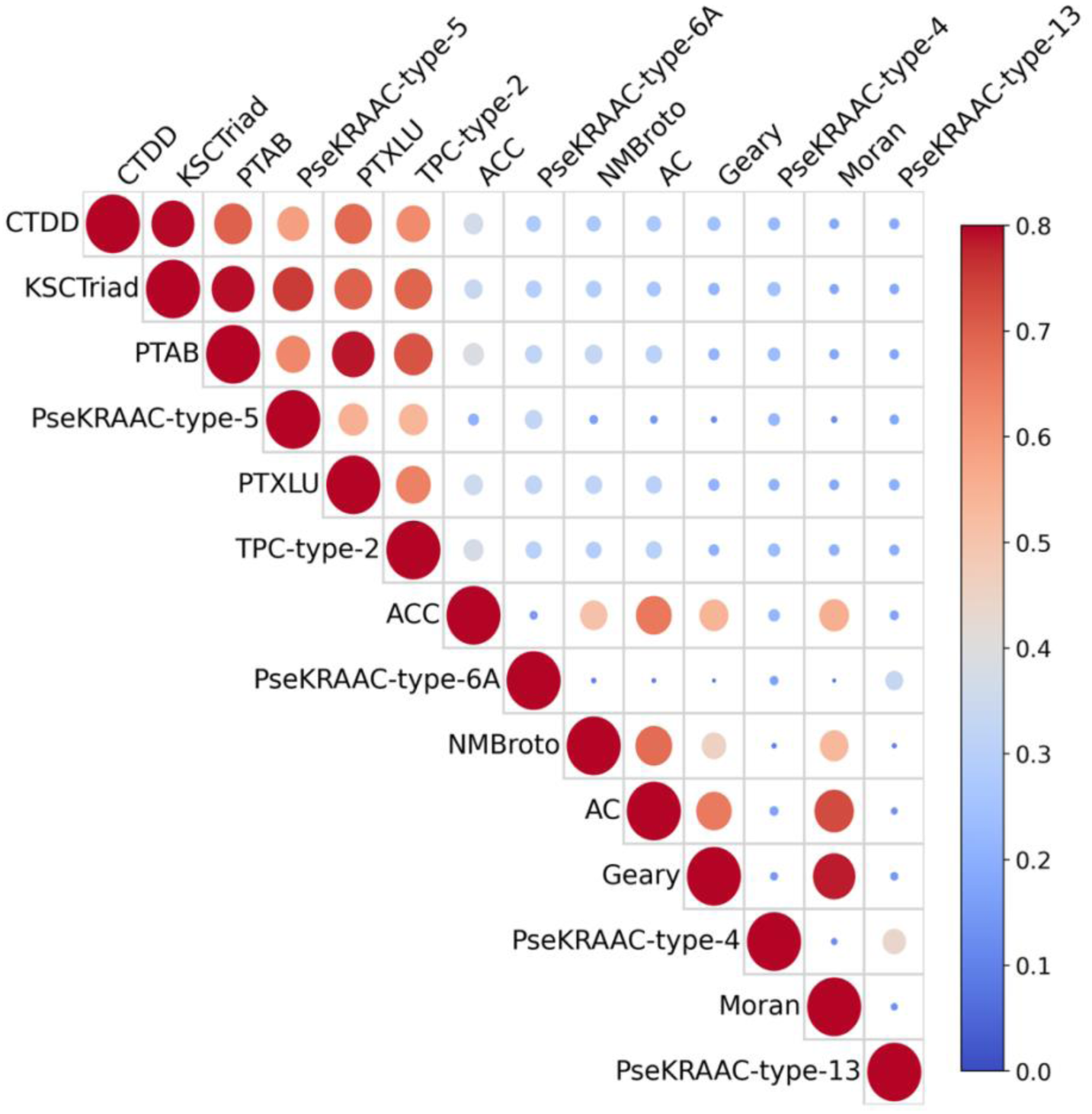
Prediction-level Pearson correlation among the 14 non-redundant feature representations retained after correlation-based filtering. Circle size and color intensity represent the magnitude of the correlation coefficients.

### Construction of StackHPpred through feature- and classifier-level integration

To determine whether integrating the selected feature representations and classifiers could further improve HP prediction, we explored feature- and classifier-level integration strategies using the out-of-fold predicted probabilities generated by the baseline models during cross-validation. For each strategy, the five top-performing models were selected based on CV MCC, and their training and independent performance is presented in Fig. 4A–B and Fig. 5A–B, respectively. In the feature-level strategy, the selected features were progressively integrated based on their average CV MCC ranking. For each feature, the predicted probabilities generated by the 10 classifiers were concatenated to construct meta-feature representations, resulting in configurations ranging from Top-1 to Top-14. These meta-features were subsequently used to train secondary meta-learners. Among these models, SVM model utilizing Top-3 demonstrated best performance on the training dataset with the AUC, MCC, ACC, Sn, Sp, Precision, and F1-score values of 0.9914, 0.9626, 0.9812, 0.9767, 0.9856, 0.9856, and 0.9810, respectively. The corresponding metrics for this model on independent dataset were 0.9965, 0.9506, 0.9752, 0.9645, 0.9858, 0.9855, and 0.9749. Notably, the top-performing feature-level configurations showed only marginal differences in CV MCC, with values ranging from 0.9607 to 0.9626. Increasing the number of integrated features did not lead to a consistent improvement in CV performance. In contrast, greater variation was observed on the independent dataset, where SVM-MF-Top7 achieved the highest MCC of 0.9717. These findings indicate that model performance did not consistently improve with the inclusion of additional features, highlighting the effectiveness of a compact subset of informative representations.

**Figure 4.**
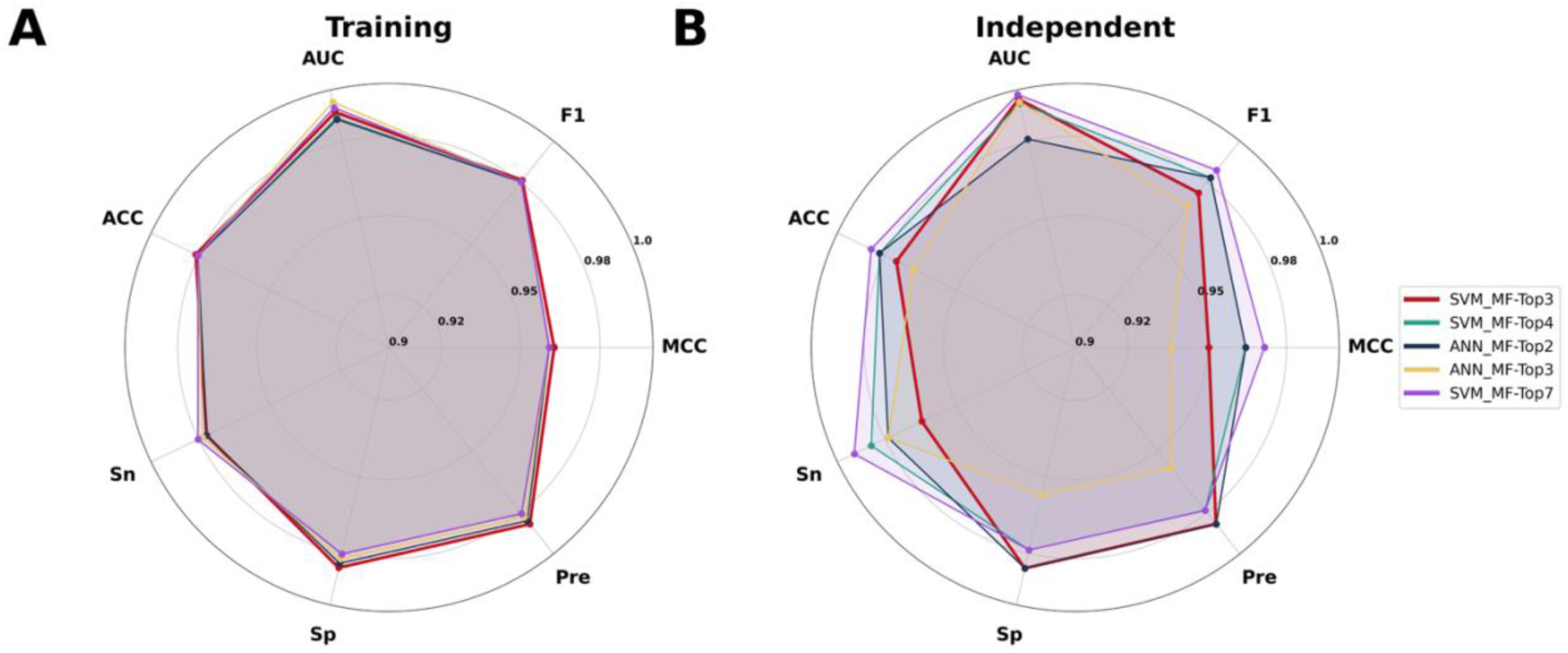
Performance comparison of the top five feature-level stacking models on the training (A) and independent (B) datasets across seven evaluation metrics: AUC, MCC, ACC, Sn, Sp, Precision, and F1-score.

**Figure 5.**
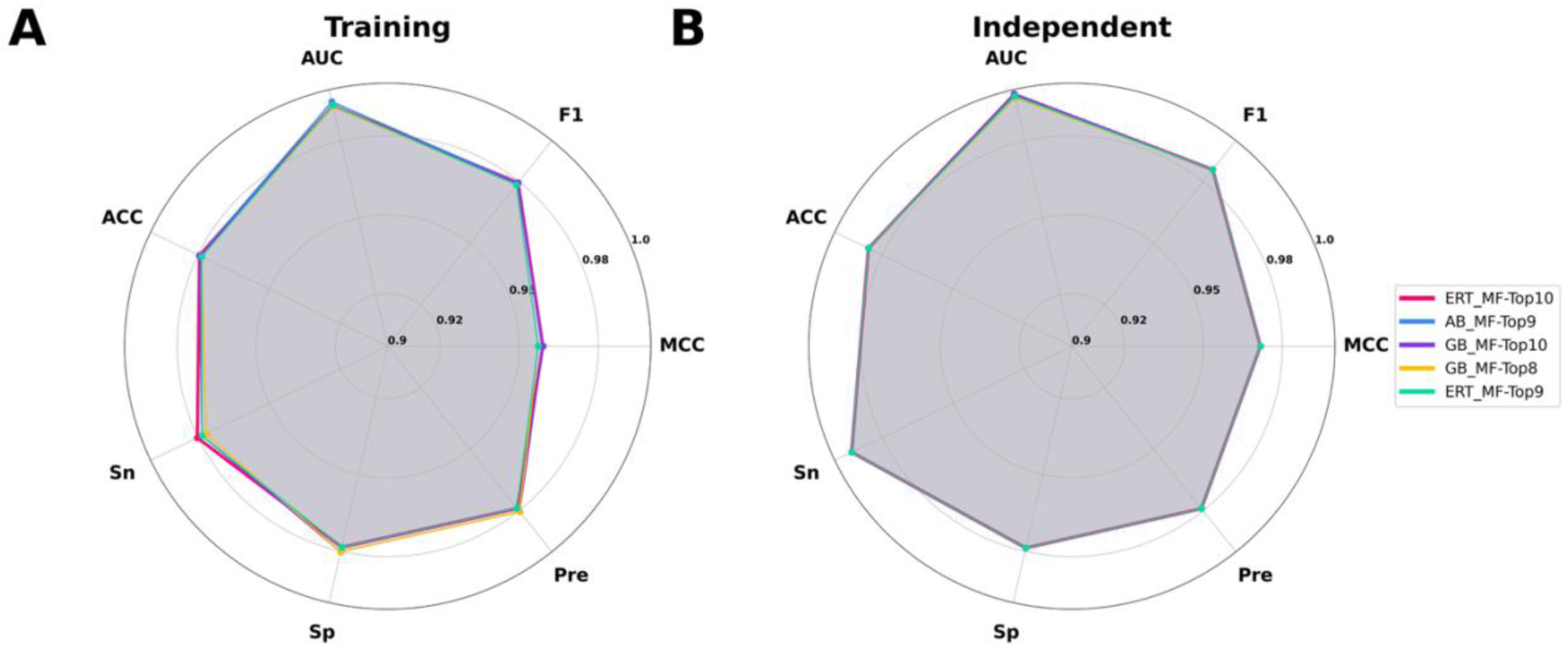
Performance comparison of the top five classifier-level stacking models on the training (A) and independent (B) datasets across seven evaluation metrics: AUC, MCC, ACC, Sn, Sp, Precision, and F1-score.

We further assessed classifier-level integration using the same out-of-fold predicted probabilities. In this strategy, the 10 classifiers were ranked according to their average CV MCC, and their predicted probabilities across all 14 selected features were progressively concatenated to generate configurations ranging from Top-1 to Top-10. The resulting meta-feature representations were subsequently used to train secondary meta-learners. Among the classifier-level models, AB-MF-Top9 and ERT-MF-Top10 achieved the highest CV MCC of 0.9590. Although both models showed identical MCC and ACC values, AB-MF-Top9 achieved a slightly higher AUC than ERT-MF-Top10 on the training dataset (0.9953 vs. 0.9938). Therefore, we further examined the performance of AB-MF-Top9, which achieved AUC, MCC, ACC, Sn, Sp, Precision, and F1-score values of 0.9953, 0.9590, 0.9794, 0.9784, 0.9802, 0.9805, and 0.9794, respectively. The corresponding values on the independent dataset were 0.9976, 0.9717, 0.9858, 0.9929, 0.9787, 0.9790, and 0.9859, respectively. Overall, the classifier-level models showed comparable cross-validation performance. However, SVM-MF-Top3 from the feature-level strategy achieved the highest CV MCC of 0.9626. Therefore, we selected the SVM-MF-Top3 model, which integrates the three highest-ranked feature representations, namely CTDD, KSCTriad, and PTAB, and designated it as StackHPpred.

### Comparison of StackHPpred with top-performing baseline models

We compared StackHPpred with the five top-performing baseline models, including CB-CTDD, XGB-CTDD, RF-CTDD, LGB-CTDD, and LRT-Seq2Vec (Fig. 6A–B). Interestingly, four of the five top-performing baseline models were based on CTDD, suggesting that the positional distribution of physicochemical properties within peptide sequences provides informative signals for distinguishing HPs from non-HPs, whereas LRT-Seq2Vec was the only model based on an NLP-derived embedding. As shown in Fig. 6A, StackHPpred achieved AUC, MCC, ACC, Sn, Sp, Precision, and F1-score values of 0.9914, 0.9626, 0.9812, 0.9767, 0.9856, 0.9856, and 0.9810, respectively, on the training dataset. Compared with the five baseline models, these results represent improvements of 0.37–1.39% in AUC, 5.56–8.14% in MCC, 2.87–4.13% in ACC, 0.45–3.13% in Precision, and 3.01–4.22% in F1-score. To further assess the generalization performance of StackHPpred, we evaluated the models on the independent dataset. As shown in Fig. 6B, StackHPpred achieved AUC, MCC, ACC, Sn, Sp, Precision, and F1-score values of 0.9965, 0.9506, 0.9752, 0.9645, 0.9858, 0.9855, and 0.9749, respectively. Compared with the same five baseline models, StackHPpred showed improvements of 0.33–1.54% in AUC, 0.73–4.76% in MCC, 0.36–2.48% in ACC, 0.03–1.39% in Precision, and 0.33–2.64% in F1-score. Overall, StackHPpred consistently surpassed the five baseline models in terms of AUC, MCC, ACC, Precision, and F1-score on both datasets, demonstrating its strong capability for distinguishing HPs from non-HPs.

**Figure 6.**
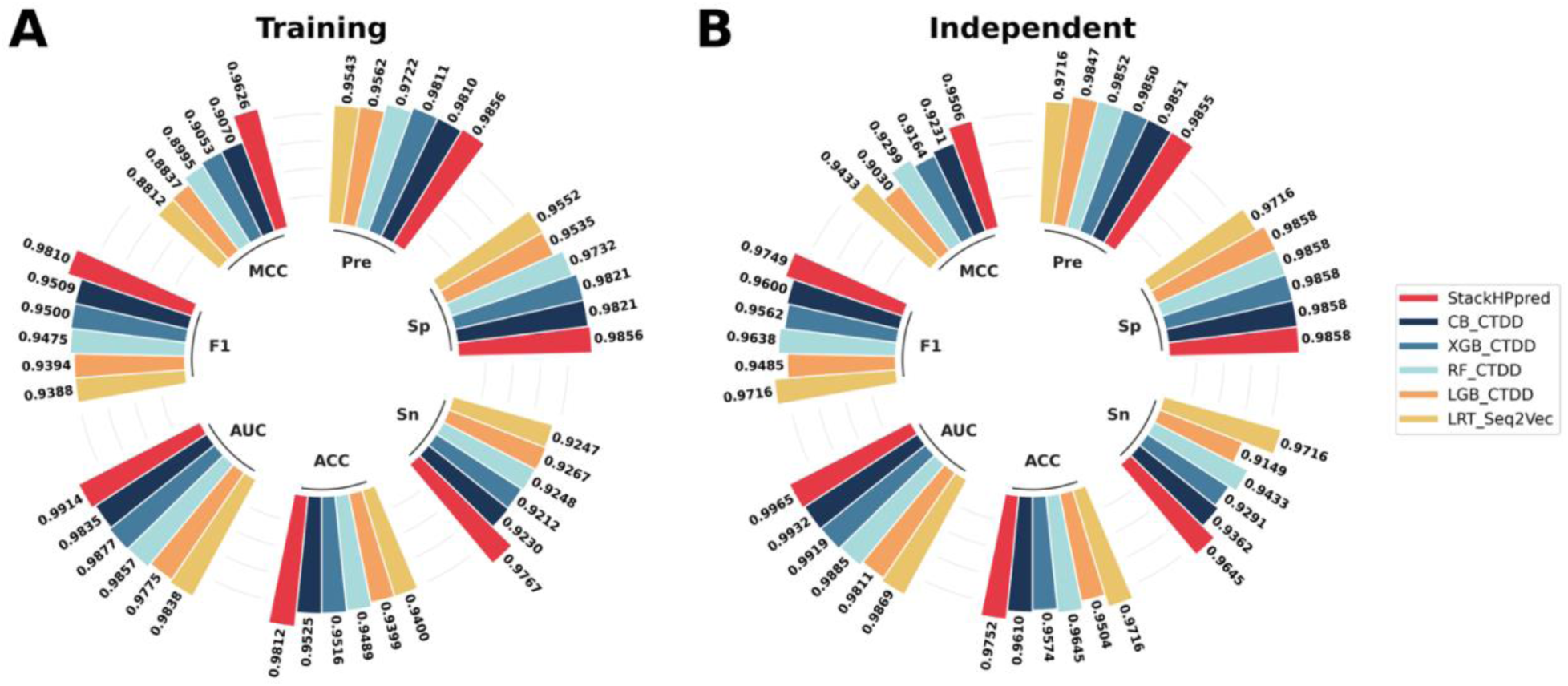
Performance comparison of StackHPpred with the five top-performing baseline models on the training (A) and independent (B) datasets across AUC, MCC, ACC, Sn, Sp, Precision, and F1-score.

### Performance comparison between StackHPpred and publicly available HP predictors using a non-overlapping independent dataset

To benchmark StackHPpred against existing peptide hormone predictors, we compared its performance with HOPPred and mHPpred using a non-overlapping independent dataset (Table 1). At the balanced Pos:Neg ratio of 1:1, StackHPpred achieved AUC, MCC, ACC, Sn, Sp, Precision, and F1-score values of 0.9993, 0.9658, 0.9853, 0.9841, 0.9858, 0.9688, and 0.9764, respectively. In comparison, the HOPPred_Hybrid model achieved 0.8243, 0.4388, 0.7500, 0.6667, 0.7872, 0.5833, and 0.6222, while mHPpred achieved 0.9172, 0.6855, 0.8480, 0.9048, 0.8227, 0.6951, and 0.7862, respectively. These results show that StackHPpred achieved higher performance across all seven-evaluation metrics on the balanced independent dataset.

**Table 1.** Performance comparison of StackHPpred and existing peptide hormone predictors on non-overlapping independent datasets with Pos:Neg ratios of 1:1, 1:10, 1:20, 1:30, 1:40, and 1:50. Performance was evaluated using AUC, MCC, ACC, Sn, Sp, Precision, and F1-score. The best value for each metric is shown in bold.

| Pos:Neg Ratio | Method | Model | AUC | MCC | ACC | Sn | Sp | Precision | F1 |
| --- | --- | --- | --- | --- | --- | --- | --- | --- | --- |
| 1:01 | HOPpred | LR | 0.6829 | 0.2671 | 0.6912 | 0.4762 | 0.7872 | 0.5000 | 0.4878 |
|  |  | Hybrid | 0.8243 | 0.4388 | 0.75 | 0.6667 | 0.7872 | 0.5833 | 0.6222 |
|  | mHPpred | LGB-Meta-20D | 0.9172 | 0.6855 | 0.848 | 0.9048 | 0.8227 | 0.6951 | 0.7862 |
|  | StackHPpred | SVM_MF-Top3 | **0.9993** | **0.9658** | **0.9853** | **0.9841** | **0.9858** | **0.9688** | **0.9764** |
| 1:10 | HOPpred | LR | 0.6671 | 0.1136 | 0.7529 | 0.4762 | 0.7652 | 0.0831 | 0.1415 |
|  |  | Hybrid | 0.8083 | 0.1979 | 0.757 | 0.6667 | 0.761 | 0.1108 | 0.1900 |
|  | mHPpred | LGB-Meta-20D | 0.9342 | 0.4268 | 0.8730 | 0.9048 | 0.8716 | 0.2395 | 0.3787 |
|  | StackHPpred | SVM_MF-Top3 | **0.9992** | **0.8616** | **0.9864** | **0.9841** | **0.9865** | **0.7654** | **0.8611** |
| 1:20 | HOPpred | LR | 0.6799 | 0.0856 | 0.7651 | 0.4762 | 0.7716 | 0.0445 | 0.0814 |
|  |  | Hybrid | 0.8159 | 0.1491 | 0.7668 | 0.6667 | 0.7691 | 0.0606 | 0.1111 |
|  | mHPpred | LGB-Meta-20D | 0.934 | 0.3263 | 0.8754 | 0.9048 | 0.8748 | 0.139 | 0.241 |
|  | StackHPpred | SVM_MF-Top3 | **0.9989** | **0.8098** | **0.9892** | **0.9841** | **0.9894** | **0.6739** | **0.8000** |
| 1:30 | HOPpred | LR | 0.6767 | 0.0701 | 0.7661 | 0.4762 | 0.7704 | 0.0300 | 0.0564 |
|  |  | Hybrid | 0.8145 | 0.1220 | 0.765 | 0.6667 | 0.7664 | 0.0408 | 0.0769 |
|  | mHPpred | LGB-Meta-20D | 0.9338 | 0.2627 | 0.8663 | 0.9048 | 0.8657 | 0.0912 | 0.1657 |
|  | StackHPpred | SVM_MF-Top3 | **0.9993** | **0.7696** | **0.9904** | **0.9841** | **0.9905** | **0.6078** | **0.7515** |
| 1:40 | HOPpred | LR | 0.671 | 0.0597 | 0.7631 | 0.4762 | 0.7663 | 0.0223 | 0.0425 |
|  |  | Hybrid | 0.8112 | 0.1051 | 0.7626 | 0.6667 | 0.7637 | 0.0305 | 0.0584 |
|  | mHPpred | LGB-Meta-20D | 0.9311 | 0.2227 | 0.8581 | 0.9048 | 0.8576 | 0.0663 | 0.1235 |
|  | StackHPpred | SVM_MF-Top3 | **0.9990** | **0.6728** | **0.9874** | **0.9841** | **0.9874** | **0.4662** | **0.6327** |
| 1:50 | HOPpred | LR | 0.6751 | 0.0541 | 0.7655 | 0.4762 | 0.7681 | 0.0180 | 0.0347 |
|  |  | Hybrid | 0.8141 | 0.0949 | 0.7642 | 0.6667 | 0.7651 | 0.0247 | 0.0477 |
|  | mHPpred | LGB-Meta-20D | 0.9314 | 0.2063 | 0.8649 | 0.9048 | 0.8645 | 0.0563 | 0.1060 |
|  | StackHPpred | SVM_MF-Top3 | **0.9991** | **0.6737** | **0.9899** | **0.9841** | **0.9899** | **0.4662** | **0.6327** |

We next examined whether this performance advantage was maintained under increasingly imbalanced conditions using Pos:Neg ratios of 1:10, 1:20, 1:30, 1:40, and 1:50. At the 1:10 ratio, StackHPpred achieved an MCC of 0.8616 and Precision of 0.7654, compared with 0.1979 and 0.1108 for HOPPred_Hybrid model and 0.4268 and 0.2395 for mHPpred, respectively. This advantage was retained as class imbalance became more severe. At the 1:50 ratio, StackHPpred maintained an MCC of 0.6737, F1-score of 0.6327, and Precision of 0.4662, whereas the corresponding values were 0.0949, 0.0477, and 0.0247 for HOPPred_Hybrid model and 0.2063, 0.1060, and 0.0563 for mHPpred. Notably, the Precision of StackHPpred was approximately 18.9- and 8.3-fold higher than that of HOPPred_Hybrid model and mHPpred, respectively.

StackHPpred also showed a smaller overall decline in MCC under increasing class imbalance. From the 1:10 to 1:50 ratios, MCC decreased by 21.8% for StackHPpred, compared with 52.0% for HOPPred_Hybrid model and 51.7% for mHPpred. Moreover, StackHPpred maintained a sensitivity of 0.9841 and specificity ranging from 0.9865 to 0.9905 across the imbalanced datasets, indicating strong control of false-positive predictions while retaining high sensitivity. Together, these results indicate that StackHPpred provides more robust and balanced discrimination of HPs and non-HPs under both balanced and increasingly imbalanced conditions.

### Visualization of discriminative feature space using t-SNE

To investigate how effectively different feature representations separate HPs from non-HPs, we projected the samples into a two-dimensional space using t-distributed stochastic neighbor embedding (t-SNE) [45]. For comparison with StackHPpred, we selected the five highest-ranked features retained after correlation-based filtering, namely CTDD, KSCTriad, PTAB, PseKRAAC-type-5, and PTXLU. These five features were ranked based on their average CV MCC and represent the top-performing non-redundant feature representations. While the individual feature representations showed varying degrees of overlap between HPs and non-HPs on the training dataset (Fig. 7A–E), the StackHPpred representation showed a more pronounced separation between the two classes (Fig. 7F). This clearer separation, with substantially reduced overlap between HPs and non-HPs, highlights the improved discriminative capability of the integrated representation. Importantly, a similar pattern of enhanced separation was consistently observed on the independent dataset (Fig. 7G–K), where the individual feature representations continued to show considerable overlap, whereas StackHPpred maintained a distinct separation between the two classes. These findings suggest that the integrated representation generated by StackHPpred is more effective at differentiating HPs from non-HPs than the individual feature representations. The consistent separation observed across both the training and independent datasets further indicates that StackHPpred effectively captures complementary information from multiple features for distinguishing HPs from non-HPs.

**Figure 7.**
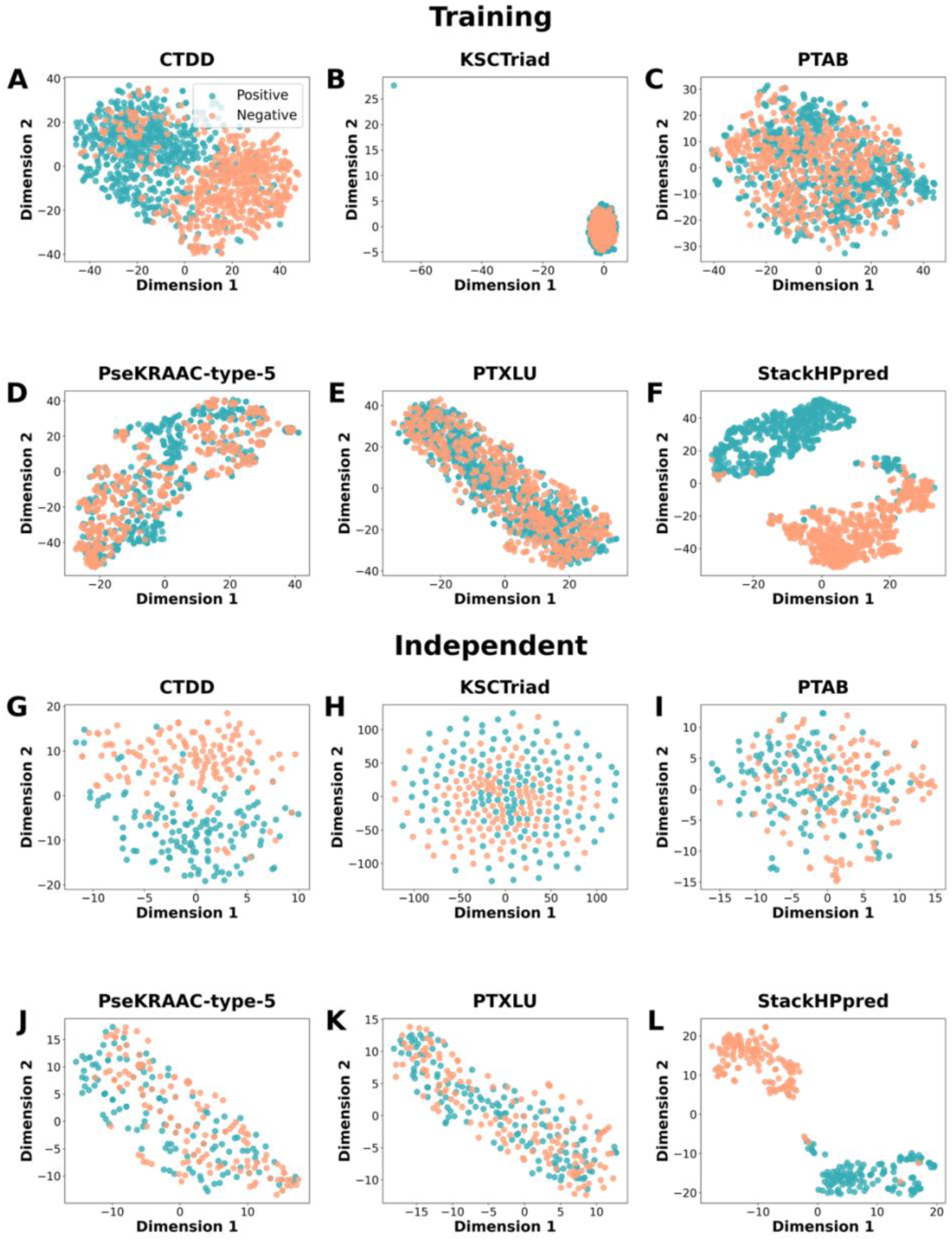
t-SNE visualization of HP and non-HP distributions using the top five non-redundant feature representations and StackHPpred. Panels A–F show CTDD, KSCTriad, PTAB, PseKRAAC-type-5, PTXLU, and StackHPpred, respectively, on the training dataset, while panels G–L show the corresponding distributions on the independent dataset.

### SHAP-based feature contribution analysis

To interpret the contribution of individual meta-features to the final prediction of StackHPpred, we performed SHapley Additive exPlanations (SHAP) analysis [46]. Figure 8A shows the mean absolute SHAP values, which rank the meta-features according to their overall importance to the model output. XGB-CTDD showed the highest contribution, followed by AB-KSCTriad, GB-CTDD, RF-KSCTriad, and LGB-CTDD. Notably, CTDD- and KSCTriad-based meta-features were highly represented among the top-ranked contributors, highlighting their substantial influence on StackHPpred predictions. The frequent occurrence of CTDD-based meta-features further emphasizes the relevance of the positional distribution of physicochemical properties for distinguishing HPs from non-HPs, while PTAB-based meta-features also contributed to the final prediction with comparatively lower importance.

**Figure 8.**
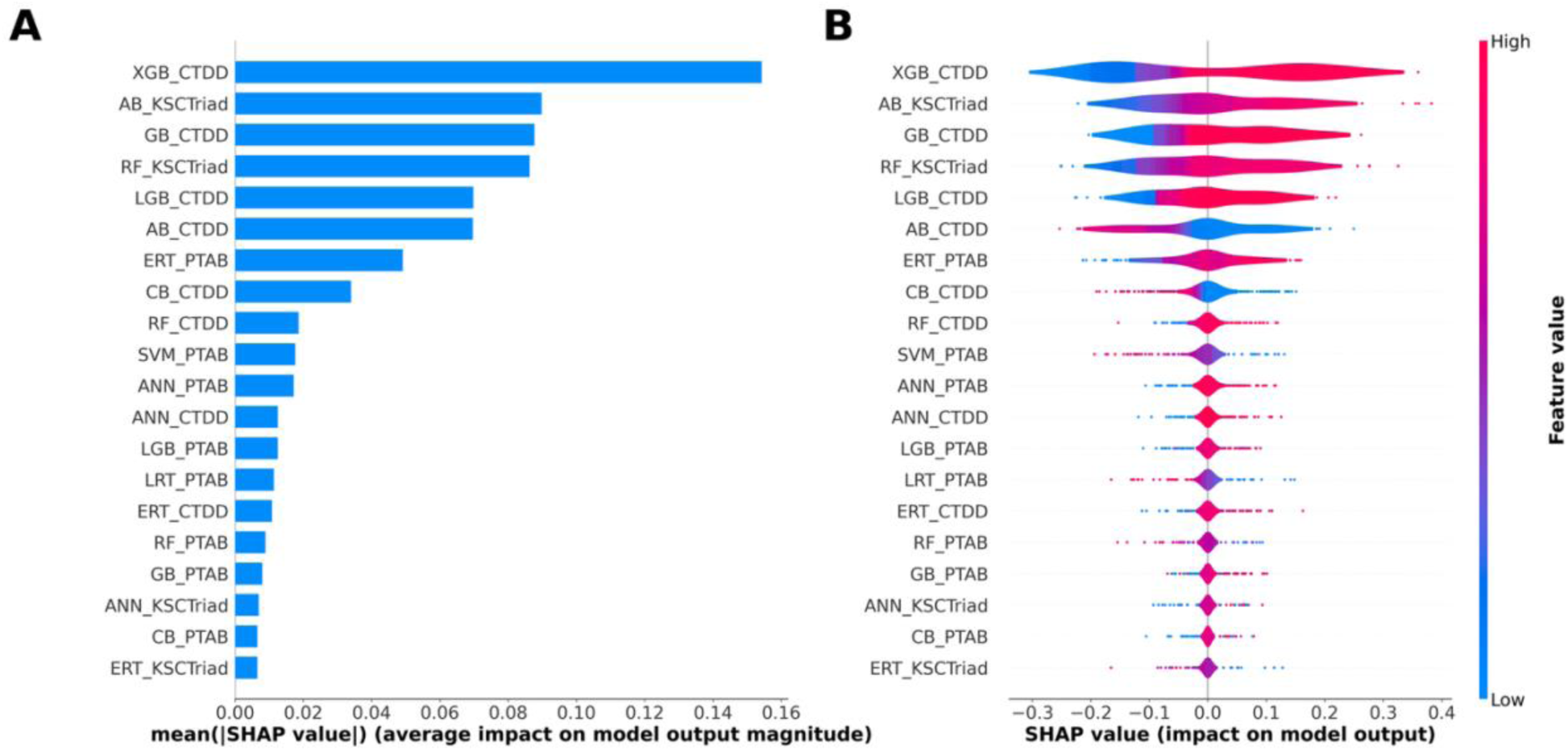
SHAP-based interpretation of StackHPpred. (A) Mean absolute SHAP values ranking the meta-features by their overall contribution to model output. (B) SHAP summary plot showing the magnitude and direction of each meta-feature’s effect, with blue and red representing low and high feature values, respectively. Positive SHAP values favor HP prediction, whereas negative SHAP values favor non-HP prediction.

To further examine the direction of these contributions, we generated a SHAP summary plot (Fig. 8B), where positive and negative SHAP values indicate contributions toward HP and non-HP predictions, respectively, and the color gradient represents the magnitude of each meta-feature value. For XGB-CTDD, higher feature values were generally associated with positive SHAP values, whereas lower values tended to correspond to negative SHAP values. Similar trends were observed for several other highly ranked meta-features, including AB-KSCTriad, GB-CTDD, and RF-KSCTriad. Overall, the SHAP analysis indicates that StackHPpred predictions are influenced by multiple classifier–feature combinations, with CTDD- and KSCTriad-based meta-features providing the strongest contributions to distinguishing HPs from non-HPs.

### Web server development

To facilitate convenient access and practical use of StackHPpred, we developed a freely accessible web server at https://jayasreekirthipati.org/StackHPpred/. The web server provides a user-friendly interface for submitting peptide sequences and obtaining predictions without requiring local installation or programming expertise. Users can perform prediction through the dedicated prediction page (https://jayasreekirthipati.org/StackHPpred/server/) and retrieve previously submitted jobs through the job search page (https://jayasreekirthipati.org/StackHPpred/job_search/). Detailed instructions for input preparation, job submission, and result interpretation are provided on the help page (https://jayasreekirthipati.org/StackHPpred/help/). In addition, the datasets, standalone program, and source code associated with StackHPpred are freely available through the download page (https://jayasreekirthipati.org/StackHPpred/downloads/) and the GitHub repository (https://github.com/JayasreeKirthipati04/StackHPpred), facilitating reproducible research and broader accessibility of the StackHPpred tool.

## Conclusion

In this study, we developed StackHPpred, a stacking-based ensemble framework for accurate identification of hormone peptides. Systematic benchmarking of 56 feature representations with 10 ML classifiers revealed considerable variation in their predictive capabilities, while prediction-level correlation analysis identified a compact set of complementary, non-redundant representations. Feature- and classifier-level stacking further demonstrated that combining complementary predictive information was more effective than relying on individual feature representations, with the integration of CTDD, KSCTriad, and PTAB yielding the optimal configuration. Importantly, StackHPpred showed greater robustness than existing peptide hormone predictors under increasing class imbalance, maintaining superior MCC, F1-score, Precision, and Specificity across the evaluated ratios. At a Pos:Neg ratio of 1:50, StackHPpred achieved a Precision approximately 18.9 and 8.3 times that of HOPPred_Hybrid model and mHPpred, respectively, indicating improved control of false-positive predictions under challenging screening conditions. Moreover, from the 1:10 to 1:50 ratios, its MCC decreased by only 21.8%, compared with 52.0% for HOPPred Hybrid and 51.7% for mHPpred, further demonstrating its robustness to increasing class imbalance. t-SNE visualization further showed clearer separation between HPs and non-HPs after feature integration, while SHAP analysis identified CTDD- and KSCTriad-based meta-features among the strongest contributors to model predictions. Overall, StackHPpred combines complementary sequence representations and classifier outputs through an advanced stacking-based meta-learning framework, providing a robust and interpretable tool for peptide hormone identification. The model is freely available as both a web server and standalone program, facilitating its practical application in peptide hormone identification and related biological studies.

## CRediT authorship contribution statement

**Jayasree Kirthipati:** Conceptualization, Methodology, Data curation, Software, Validation, Visualization, Writing – original draft, Writing – review & editing. **Mohan Krishna Chemarthi Ravi:** Methodology, Data curation, Software, Validation, Visualization, Writing – original draft, Writing – review & editing. **Mounica Kirthipati:** Methodology, Data curation, Software, Validation, Visualization, Writing – original draft, Writing – review & editing.

## Data availability

The StackHPpred web server, along with the newly constructed training and independent datasets, is publicly accessible at https://jayasreekirthipati.org/StackHPpred/. The standalone implementation and source code are freely available at https://github.com/JayasreeKirthipati04/StackHPpred.

## Declaration of competing interest

The authors declare that they have no known competing financial interests or personal relationships that could have appeared to influence the work reported in this paper.

## AI Disclosure Statement

During the preparation of this work, the authors used ChatGPT 5.6 to improve the fluency, clarity, and readability of the manuscript. After using this tool, the authors reviewed and edited the content as necessary and take full responsibility for the final content of the published article.

## Notes

https://jayasreekirthipati.org/StackHPpred/

https://github.com/JayasreeKirthipati04/StackHPpred

### Competing Interest Statement

The authors have declared no competing interest.

