## Supplementary Information for "StackHPpred: A Stacking-based ensemble learning framework for the identification of peptide hormones using multi-view feature representations"

**Author's Information**

| | |
|---|---|
| Jayasree Kirthipati: | ORCID ID: https://orcid.org/0009-0005-1087-3440 |
| Mohan Krishna Chemarthi Ravi: | ORCID ID: https://orcid.org/0009-0002-7919-4735 |
| Mounica Kirthipati: | ORCID ID: https://orcid.org/0009-0006-6333-6486 |

**Author’s Biography**

1. Jayasree Kirthipati received her master’s degree in computer science from University of South Dakota, Vermillion, South Dakota, 57069, United States of America.
2. Mohan Krishna Chemarthi Ravi is currently pursuing a Pharm.D. at the Department of Pharmaceutical sciences, Krishna Teja Pharmacy College (affiliated with Jawaharlal Nehru Technological University Anantapur (JNTUA)), Chadalawada Nagar, Renigunta Road, Tirupati, Andhra Pradesh, 517506, India.
3. Mounica Kirthipati received her bachelor’s degree in Electronics and Communication Engineering from Siddharth Institute of Engineering and Technology, Puttur, Andhra Pradesh, 57069, India.

**Supplementary Figures**

**Figure S1.** Sequence length distributions of hormone and non-hormone peptides in the combined training and independent datasets. Histograms and density curves illustrate the distributions of 699 HPs and 699 non-HPs.

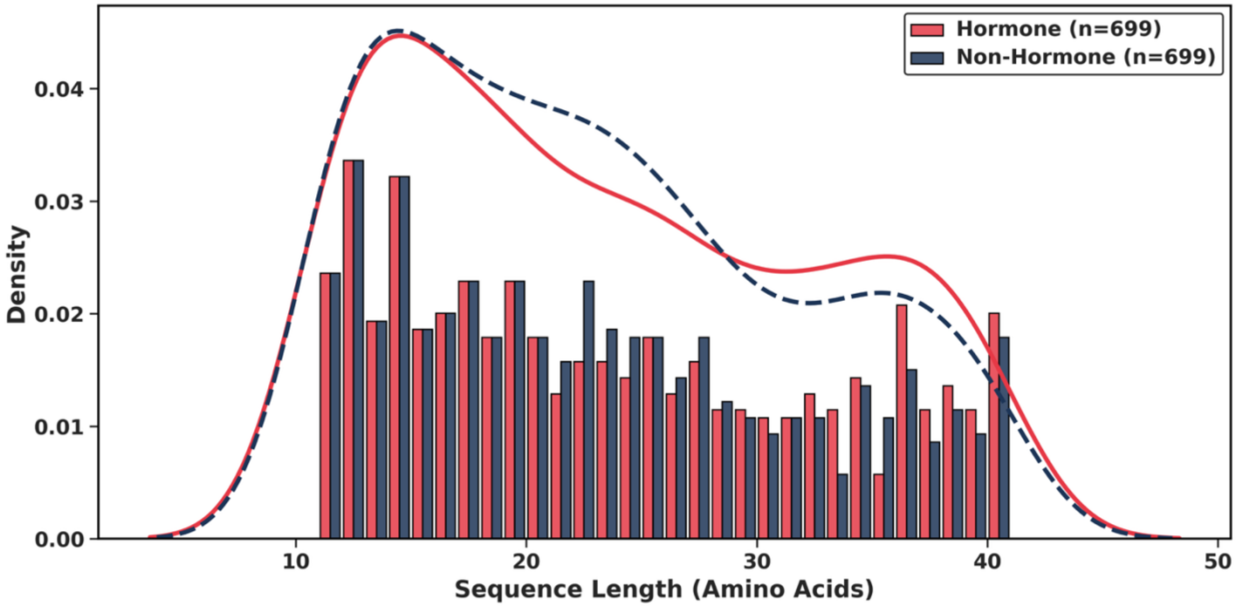

**Figure S2.** Amino acid composition analysis of hormone peptides (HPs) and non-hormone peptides (non-HPs). Average amino acid composition (%) was compared between the two classes. Statistical significance levels: ***$p < 0.001$, *$p < 0.01$, $p < 0.05$, *and ns = not significant.*

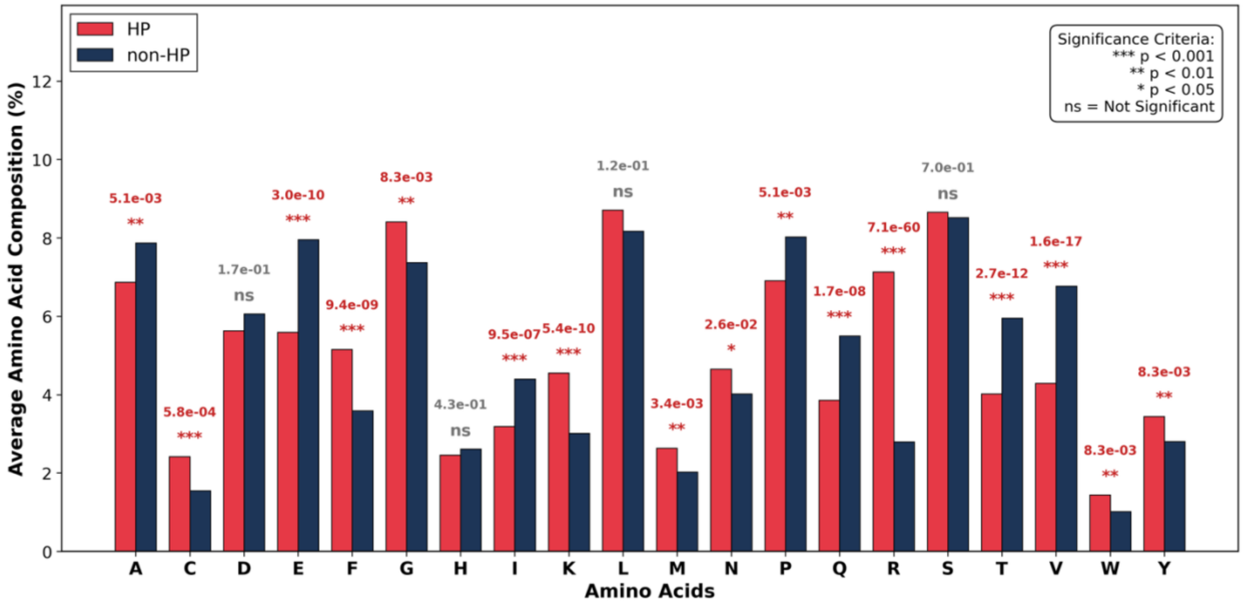

**Figure S3.** Two-sample logo analysis of positional amino acid distribution in hormone peptides (HPs) and non-hormone peptides (non-HPs). The N-terminal (A) and C-terminal (B) 15 residues are shown. Residues above the baseline are enriched in HPs, whereas residues below the baseline are depleted.

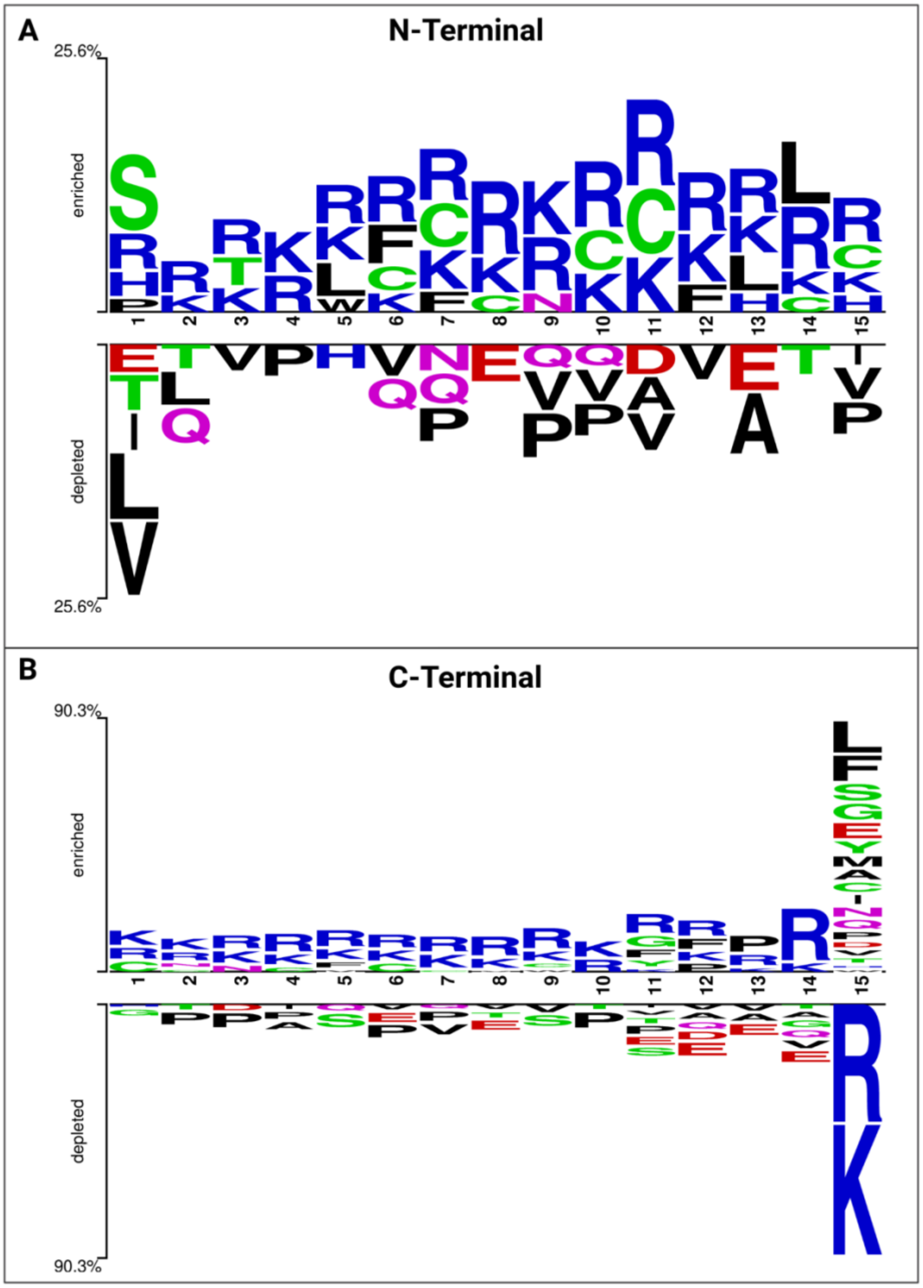

**Figure S4.** Prediction-level Pearson correlation matrix among the 56 feature representations. Pairwise correlations were calculated using the averaged out-of-fold predicted probabilities across the 10 classifiers. Color intensity represents the absolute Pearson correlation coefficient (|r|), while bold labels indicate the 14 non-redundant feature representations retained after correlation-based filtering.

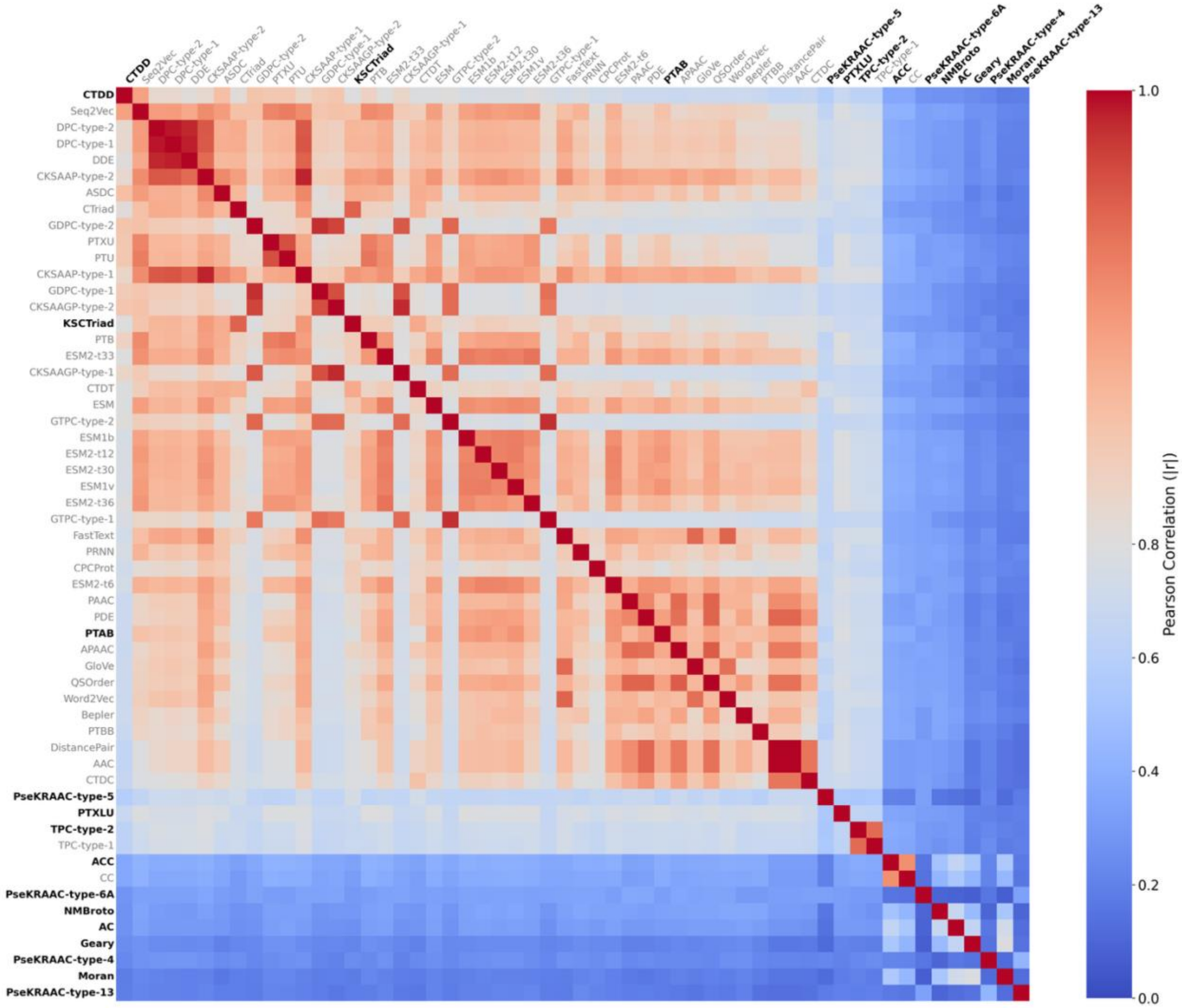

**Supplementary Tables**

**Table S1.** Summary of the 21 PLM/NLP-based embeddings and 35 conventional-based descriptors used in this study, along with their variants and feature dimensions.

| Feature types | Features | Variant | Dimension |
|---|---|---|---|
| PLM/NLP-based embeddings | Bepler | - | 121-D |
| | CPCProt | - | 512-D |
| | ESM2 | esm2-t6-8M_UR50D (ESM2-t6) | 320-D |
| | | esm2-t12-35M_UR50D (ESM2-t12) | 480-D |
| | | esm2-t30-150M_UR50D (ESM2-t30) | 640-D |
| | | esm2-t33-650M_UR50D (ESM2-t33) | 1280-D |
| | | esm2-t36-3B_UR50D (ESM2-t36) | 2560-D |
| | ESM | ESM | 1280-D |
| | | ESM1b | 1280-D |
| | | ESM1v | 1280-D |
| | FastText | - | 512-D |
| | GloVe | - | 512-D |
| | PRNN | - | 1024-D |
| | PTAB | - | 4096-D |
| | PTBB | - | 1024-D |
| | PTB | - | 1024-D |
| | PTU | - | 1024-D |
| | PTXU | - | 1024-D |
| | PTXLU | - | 1024-D |
| | Seq2Vec | - | 1024-D |
| | Word2Vec | - | 512-D |

| | | | |
|---|---|---|---|
| Conventional-based descriptors | AAC | - | 20-D |
| | ACC | - | 192-D |
| | AC | - | 24-D |
| | APAAC | - | 26-D |
| | ASDC | - | 400-D |
| | CC | - | 168-D |
| | CKSAAGP | Type-1 | 100-D |
| | | Type-2 | 100-D |
| | CKSAAP | Type-1 | 1600-D |
| | | Type-2 | 1600-D |
| | CTDC | - | 39-D |
| | CTDD | - | 195-D |
| | CTDT | - | 39-D |
| | CTriad | - | 343-D |
| | DDE | - | 400-D |
| | DistancePair | - | 20-D |
| | DPC | Type-1 | 400-D |
| | | Type-2 | 400-D |
| | GDPC | Type-1 | 25-D |
| | | Type-2 | 25-D |
| | Geary | - | 24-D |
| | GTPC | Type-1 | 125-D |
| | | Type-2 | 125-D |
| | KSCTriad | - | 1372-D |
| | Moran | - | 24-D |
| | NMBroto | - | 24-D |
| | PAAC | - | 23-D |

| | | | |
|---|---|---|---|
| | PDE | - | 48-D |
| | PseKRAAC | Type-4 | 25-D |
| | | Type-5 | 9-D |
| | | Type-6A | 16-D |
| | | Type-13 | 16-D |
| | QSOrder | - | 46-D |
| | TPC | Type-1 | 8000-D |
| | | Type-2 | 8000-D |

**Table S2.** Performance of 560 single-feature baseline models on the training and independent datasets.

| Classifier | Feature | Training | | | | | | | Independent | | | | | | |
|---|---|---|---|---|---|---|---|---|---|---|---|---|---|---|---|
| | | AUC | MCC | ACC | Sn | Sp | Precision | F1 | AUC | MCC | ACC | Sn | Sp | Precision | F1 |
| AB | Bepler | 0.8968 | 0.6256 | 0.8109 | 0.776 | 0.8458 | 0.8359 | 0.8031 | 0.921 | 0.7107 | 0.8546 | 0.8227 | 0.8865 | 0.8788 | 0.8498 |
| AB | CPCProt | 0.9157 | 0.7066 | 0.8513 | 0.819 | 0.8836 | 0.8768 | 0.8451 | 0.9197 | 0.6924 | 0.844 | 0.7872 | 0.9007 | 0.888 | 0.8346 |
| AB | ESM | 0.9348 | 0.7412 | 0.8691 | 0.8332 | 0.9049 | 0.8994 | 0.8644 | 0.9501 | 0.7385 | 0.8688 | 0.844 | 0.8936 | 0.8881 | 0.8655 |
| AB | ESM1b | 0.9167 | 0.707 | 0.8522 | 0.8228 | 0.8817 | 0.875 | 0.8471 | 0.931 | 0.6955 | 0.8475 | 0.8298 | 0.8652 | 0.8603 | 0.8448 |
| AB | ESM1v | 0.9199 | 0.7059 | 0.8522 | 0.8297 | 0.8745 | 0.8692 | 0.8484 | 0.9409 | 0.7199 | 0.8582 | 0.8085 | 0.9078 | 0.8976 | 0.8507 |
| AB | ESM2-t6 | 0.8931 | 0.6541 | 0.8262 | 0.8101 | 0.8424 | 0.8374 | 0.8225 | 0.9175 | 0.647 | 0.8227 | 0.7872 | 0.8582 | 0.8473 | 0.8162 |
| AB | ESM2-t12 | 0.9256 | 0.7243 | 0.8611 | 0.8442 | 0.8781 | 0.8743 | 0.8579 | 0.9421 | 0.7175 | 0.8582 | 0.8298 | 0.8865 | 0.8797 | 0.854 |
| AB | ESM2-t30 | 0.9198 | 0.7211 | 0.8585 | 0.828 | 0.8889 | 0.8831 | 0.8524 | 0.9474 | 0.7541 | 0.8759 | 0.8369 | 0.9149 | 0.9077 | 0.8708 |
| AB | ESM2-t33 | 0.9423 | 0.7563 | 0.8764 | 0.8422 | 0.9104 | 0.9046 | 0.8709 | 0.9563 | 0.775 | 0.8865 | 0.8511 | 0.922 | 0.916 | 0.8824 |
| AB | ESM2-t36 | 0.9478 | 0.7716 | 0.8835 | 0.8369 | 0.93 | 0.9237 | 0.8771 | 0.9677 | 0.8095 | 0.9043 | 0.8794 | 0.9291 | 0.9254 | 0.9018 |

| | | | | | | | | | | | | | | | |
|---|---|---|---|---|---|---|---|---|---|---|---|---|---|---|---|
| AB | FastText | 0.9154 | 0.7226 | 0.8594 | 0.8245 | 0.8944 | 0.8881 | 0.8537 | 0.9271 | 0.717 | 0.8582 | 0.8369 | 0.8794 | 0.8741 | 0.8551 |
| AB | GloVe | 0.9011 | 0.6641 | 0.8316 | 0.8118 | 0.8515 | 0.8453 | 0.8279 | 0.9167 | 0.6456 | 0.8227 | 0.8085 | 0.8369 | 0.8321 | 0.8201 |
| AB | PRNN | 0.9262 | 0.7074 | 0.8521 | 0.8207 | 0.8835 | 0.8771 | 0.8467 | 0.9492 | 0.7879 | 0.8936 | 0.8723 | 0.9149 | 0.9111 | 0.8913 |
| AB | PTAB | 0.9066 | 0.6959 | 0.8468 | 0.8298 | 0.8639 | 0.8604 | 0.8436 | 0.925 | 0.6529 | 0.8262 | 0.8085 | 0.844 | 0.8382 | 0.8231 |
| AB | PTBB | 0.8922 | 0.6181 | 0.8083 | 0.7868 | 0.8299 | 0.8231 | 0.8039 | 0.9444 | 0.7518 | 0.8759 | 0.8723 | 0.8794 | 0.8786 | 0.8754 |
| AB | PTB | 0.9498 | 0.7653 | 0.8809 | 0.8603 | 0.9015 | 0.899 | 0.8773 | 0.9697 | 0.811 | 0.9043 | 0.8652 | 0.9433 | 0.9385 | 0.9004 |
| AB | PTU | 0.9645 | 0.782 | 0.8898 | 0.8728 | 0.9069 | 0.9051 | 0.8874 | 0.9665 | 0.8102 | 0.9043 | 0.8723 | 0.9362 | 0.9318 | 0.9011 |
| AB | PTXU | 0.9668 | 0.7964 | 0.897 | 0.8745 | 0.9194 | 0.9161 | 0.8937 | 0.9677 | 0.7953 | 0.8972 | 0.8723 | 0.922 | 0.9179 | 0.8945 |
| AB | PTXLU | 0.879 | 0.6012 | 0.7993 | 0.7884 | 0.8101 | 0.8074 | 0.7959 | 0.8799 | 0.5466 | 0.773 | 0.7943 | 0.7518 | 0.7619 | 0.7778 |
| AB | Seq2Vec | 0.9801 | 0.8677 | 0.9328 | 0.905 | 0.9606 | 0.9586 | 0.9305 | 0.9905 | 0.9434 | 0.9716 | 0.9645 | 0.9787 | 0.9784 | 0.9714 |
| AB | Word2Vec | 0.8971 | 0.6834 | 0.8396 | 0.7974 | 0.8818 | 0.8717 | 0.8313 | 0.9293 | 0.6838 | 0.8404 | 0.7943 | 0.8865 | 0.875 | 0.8327 |
| AB | AAC | 0.8699 | 0.6033 | 0.8002 | 0.7617 | 0.8387 | 0.8258 | 0.7915 | 0.8949 | 0.6335 | 0.8156 | 0.773 | 0.8582 | 0.845 | 0.8074 |
| AB | ACC | 0.6979 | 0.3095 | 0.6542 | 0.6615 | 0.647 | 0.6528 | 0.656 | 0.6974 | 0.2838 | 0.6418 | 0.6312 | 0.6525 | 0.6449 | 0.638 |

| | | | | | | | | | | | | | | | |
|---|---|---|---|---|---|---|---|---|---|---|---|---|---|---|---|
| AB | AC | 0.6509 | 0.2427 | 0.621 | 0.604 | 0.6381 | 0.626 | 0.6141 | 0.6265 | 0.1718 | 0.5851 | 0.5177 | 0.6525 | 0.5984 | 0.5551 |
| AB | APAAC | 0.9127 | 0.6653 | 0.8307 | 0.7941 | 0.8673 | 0.8584 | 0.8234 | 0.9222 | 0.6675 | 0.8333 | 0.8085 | 0.8582 | 0.8507 | 0.8291 |
| AB | ASDC | 0.9534 | 0.784 | 0.8907 | 0.8673 | 0.9139 | 0.9112 | 0.8875 | 0.9734 | 0.8447 | 0.922 | 0.9007 | 0.9433 | 0.9407 | 0.9203 |
| AB | CC | 0.6919 | 0.2857 | 0.6425 | 0.6364 | 0.6487 | 0.6445 | 0.6397 | 0.6912 | 0.3269 | 0.6631 | 0.6312 | 0.695 | 0.6742 | 0.652 |
| AB | CKSAAGP-type-1 | 0.9432 | 0.7381 | 0.8683 | 0.8549 | 0.8818 | 0.8786 | 0.8658 | 0.9818 | 0.88 | 0.9397 | 0.922 | 0.9574 | 0.9559 | 0.9386 |
| AB | CKSAAGP-type-2 | 0.9515 | 0.7854 | 0.8916 | 0.8746 | 0.9087 | 0.9063 | 0.8891 | 0.9832 | 0.8795 | 0.9397 | 0.9362 | 0.9433 | 0.9429 | 0.9395 |
| AB | CKSAAP-type-1 | 0.9662 | 0.8462 | 0.9194 | 0.864 | 0.9749 | 0.9732 | 0.9137 | 0.9732 | 0.8559 | 0.9255 | 0.8723 | 0.9787 | 0.9762 | 0.9213 |
| AB | CKSAAP-type-2 | 0.961 | 0.8263 | 0.9095 | 0.8514 | 0.9678 | 0.9643 | 0.9031 | 0.9754 | 0.8652 | 0.9291 | 0.8652 | 0.9929 | 0.9919 | 0.9242 |
| AB | CTDC | 0.8706 | 0.6209 | 0.8092 | 0.7723 | 0.846 | 0.835 | 0.8019 | 0.9016 | 0.6695 | 0.8333 | 0.7872 | 0.8794 | 0.8672 | 0.8253 |
| AB | CTDD | 0.9766 | 0.8671 | 0.9328 | 0.957 | 0.9086 | 0.9131 | 0.9343 | 0.9845 | 0.88 | 0.9397 | 0.9574 | 0.922 | 0.9247 | 0.9408 |
| AB | CTDT | 0.9099 | 0.7137 | 0.8539 | 0.8028 | 0.9051 | 0.8966 | 0.8455 | 0.9213 | 0.7352 | 0.8652 | 0.8085 | 0.922 | 0.912 | 0.8571 |
| AB | CTriad | 0.9423 | 0.8006 | 0.8961 | 0.8297 | 0.9624 | 0.9568 | 0.8873 | 0.9583 | 0.808 | 0.9007 | 0.8369 | 0.9645 | 0.9593 | 0.8939 |
| AB | DDE | 0.9684 | 0.8715 | 0.932 | 0.8711 | 0.9929 | 0.992 | 0.9267 | 0.9848 | 0.8669 | 0.9291 | 0.8582 | 1 | 1 | 0.9237 |

| | | | | | | | | | | | | | | | |
|---|---|---|---|---|---|---|---|---|---|---|---|---|---|---|---|
| AB | DistancePair | 0.8699 | 0.6033 | 0.8002 | 0.7617 | 0.8387 | 0.8258 | 0.7915 | 0.8949 | 0.6335 | 0.8156 | 0.773 | 0.8582 | 0.845 | 0.8074 |
| AB | DPC-type-1 | 0.966 | 0.8646 | 0.9284 | 0.8657 | 0.9911 | 0.9898 | 0.9227 | 0.9815 | 0.8795 | 0.9362 | 0.8723 | 1 | 1 | 0.9318 |
| AB | DPC-type-2 | 0.9699 | 0.8791 | 0.9356 | 0.8729 | 0.9982 | 0.9981 | 0.9303 | 0.9781 | 0.8732 | 0.9326 | 0.8652 | 1 | 1 | 0.9278 |
| AB | GDPC-type-1 | 0.9596 | 0.8049 | 0.9015 | 0.889 | 0.914 | 0.9125 | 0.8996 | 0.9897 | 0.8795 | 0.9397 | 0.9362 | 0.9433 | 0.9429 | 0.9395 |
| AB | GDPC-type-2 | 0.9676 | 0.8493 | 0.9239 | 0.9212 | 0.9266 | 0.9269 | 0.9232 | 0.9887 | 0.9078 | 0.9539 | 0.9504 | 0.9574 | 0.9571 | 0.9537 |
| AB | Geary | 0.6243 | 0.1834 | 0.5914 | 0.586 | 0.5968 | 0.5929 | 0.5886 | 0.5874 | 0.1137 | 0.5567 | 0.5248 | 0.5887 | 0.5606 | 0.5421 |
| AB | GTPC-type-1 | 0.9081 | 0.6697 | 0.8343 | 0.8263 | 0.8423 | 0.84 | 0.8324 | 0.9446 | 0.8002 | 0.8972 | 0.8369 | 0.9574 | 0.9516 | 0.8906 |
| AB | GTPC-type-2 | 0.9258 | 0.7198 | 0.8594 | 0.8478 | 0.871 | 0.8687 | 0.8576 | 0.98 | 0.8511 | 0.9255 | 0.922 | 0.9291 | 0.9286 | 0.9253 |
| AB | KSCTriad | 0.9461 | 0.7893 | 0.8907 | 0.8262 | 0.9552 | 0.9494 | 0.8823 | 0.9498 | 0.7809 | 0.8865 | 0.8156 | 0.9574 | 0.9504 | 0.8779 |
| AB | Moran | 0.5907 | 0.1564 | 0.578 | 0.5789 | 0.577 | 0.5772 | 0.5771 | 0.5533 | 0.0788 | 0.539 | 0.4681 | 0.6099 | 0.5455 | 0.5038 |
| AB | NMBroto | 0.6476 | 0.2484 | 0.6236 | 0.6218 | 0.6255 | 0.6257 | 0.6221 | 0.6249 | 0.1561 | 0.578 | 0.5674 | 0.5887 | 0.5797 | 0.5735 |
| AB | PAAC | 0.9037 | 0.6676 | 0.8315 | 0.7994 | 0.8637 | 0.8574 | 0.8251 | 0.9131 | 0.6687 | 0.8333 | 0.7943 | 0.8723 | 0.8615 | 0.8266 |
| AB | PDE | 0.8992 | 0.6414 | 0.8181 | 0.7723 | 0.8636 | 0.8519 | 0.8084 | 0.922 | 0.674 | 0.8369 | 0.8227 | 0.8511 | 0.8467 | 0.8345 |

| | | | | | | | | | | | | | | | |
|---|---|---|---|---|---|---|---|---|---|---|---|---|---|---|---|
| AB | PseKRAAC-type-4 | 0.6085 | 0.1764 | 0.5877 | 0.5771 | 0.5984 | 0.5882 | 0.5813 | 0.5703 | 0.1068 | 0.5532 | 0.5106 | 0.5957 | 0.5581 | 0.5333 |
| AB | PseKRAAC-type-5 | 0.8403 | 0.5945 | 0.7895 | 0.6847 | 0.8943 | 0.8696 | 0.7637 | 0.8404 | 0.5254 | 0.7589 | 0.6738 | 0.844 | 0.812 | 0.7364 |
| AB | PseKRAAC-type-6A | 0.6618 | 0.2578 | 0.628 | 0.6056 | 0.6504 | 0.6353 | 0.6176 | 0.6699 | 0.2491 | 0.6241 | 0.5816 | 0.6667 | 0.6357 | 0.6074 |
| AB | PseKRAAC-type-13 | 0.58 | 0.1388 | 0.569 | 0.5376 | 0.6002 | 0.5743 | 0.5535 | 0.5923 | 0.1135 | 0.5567 | 0.5461 | 0.5674 | 0.558 | 0.552 |
| AB | QSOrder | 0.8939 | 0.6294 | 0.8128 | 0.7743 | 0.8513 | 0.8408 | 0.8047 | 0.9218 | 0.6959 | 0.8475 | 0.8227 | 0.8723 | 0.8657 | 0.8436 |
| AB | TPC-type-1 | 0.793 | 0.5054 | 0.7393 | 0.5807 | 0.8978 | 0.8512 | 0.689 | 0.8871 | 0.6562 | 0.8085 | 0.6383 | 0.9787 | 0.9677 | 0.7692 |
| AB | TPC-type-2 | 0.7993 | 0.5011 | 0.7384 | 0.5879 | 0.8889 | 0.8427 | 0.6909 | 0.8545 | 0.5524 | 0.766 | 0.6312 | 0.9007 | 0.8641 | 0.7295 |
| ANN | Bepler | 0.9009 | 0.6519 | 0.8234 | 0.837 | 0.8098 | 0.8199 | 0.8255 | 0.9273 | 0.7163 | 0.8582 | 0.8582 | 0.8582 | 0.8582 | 0.8582 |
| ANN | CPCProt | 0.9045 | 0.6912 | 0.844 | 0.8278 | 0.8601 | 0.8588 | 0.8412 | 0.8893 | 0.6533 | 0.8262 | 0.8014 | 0.8511 | 0.8433 | 0.8218 |
| ANN | ESM | 0.9404 | 0.7649 | 0.8817 | 0.8709 | 0.8925 | 0.8907 | 0.8799 | 0.9606 | 0.7873 | 0.8936 | 0.9007 | 0.8865 | 0.8881 | 0.8944 |
| ANN | ESM1b | 0.9296 | 0.7238 | 0.8611 | 0.8531 | 0.8692 | 0.8673 | 0.8591 | 0.9624 | 0.8015 | 0.9007 | 0.8936 | 0.9078 | 0.9065 | 0.9 |
| ANN | ESM1v | 0.9347 | 0.7306 | 0.8639 | 0.8568 | 0.871 | 0.8706 | 0.8619 | 0.9549 | 0.7385 | 0.8688 | 0.844 | 0.8936 | 0.8881 | 0.8655 |
| ANN | ESM2-t6 | 0.9225 | 0.7214 | 0.8593 | 0.8387 | 0.8801 | 0.8773 | 0.8562 | 0.9528 | 0.7977 | 0.8972 | 0.8511 | 0.9433 | 0.9375 | 0.8922 |

| | | | | | | | | | | | | | | | |
|---|---|---|---|---|---|---|---|---|---|---|---|---|---|---|---|
| ANN | ESM2-t12 | 0.9321 | 0.7293 | 0.8638 | 0.8585 | 0.8692 | 0.8679 | 0.8621 | 0.9479 | 0.7448 | 0.8723 | 0.8652 | 0.8794 | 0.8777 | 0.8714 |
| ANN | ESM2-t30 | 0.9283 | 0.7162 | 0.8567 | 0.8496 | 0.8639 | 0.8637 | 0.8548 | 0.9556 | 0.7731 | 0.8865 | 0.8936 | 0.8794 | 0.8811 | 0.8873 |
| ANN | ESM2-t33 | 0.9481 | 0.7796 | 0.8889 | 0.8708 | 0.907 | 0.9042 | 0.8864 | 0.9782 | 0.8795 | 0.9397 | 0.9362 | 0.9433 | 0.9429 | 0.9395 |
| ANN | ESM2-t36 | 0.9528 | 0.7666 | 0.8825 | 0.8709 | 0.8942 | 0.893 | 0.8809 | 0.9664 | 0.8369 | 0.9184 | 0.9149 | 0.922 | 0.9214 | 0.9181 |
| ANN | FastText | 0.907 | 0.7275 | 0.8621 | 0.8334 | 0.8907 | 0.8849 | 0.8567 | 0.9256 | 0.6951 | 0.8475 | 0.8511 | 0.844 | 0.8451 | 0.8481 |
| ANN | GloVe | 0.9013 | 0.6924 | 0.845 | 0.8263 | 0.8639 | 0.86 | 0.8416 | 0.8989 | 0.6328 | 0.8156 | 0.7801 | 0.8511 | 0.8397 | 0.8088 |
| ANN | PRNN | 0.9415 | 0.7399 | 0.8692 | 0.8637 | 0.8745 | 0.8738 | 0.8678 | 0.9567 | 0.7943 | 0.8972 | 0.9007 | 0.8936 | 0.8944 | 0.8975 |
| ANN | PTAB | 0.9051 | 0.6871 | 0.8423 | 0.8206 | 0.8639 | 0.8593 | 0.8383 | 0.8981 | 0.6168 | 0.805 | 0.8794 | 0.7305 | 0.7654 | 0.8185 |
| ANN | PTBB | 0.8968 | 0.6618 | 0.8306 | 0.8244 | 0.837 | 0.8355 | 0.8296 | 0.931 | 0.7306 | 0.8652 | 0.8582 | 0.8723 | 0.8705 | 0.8643 |
| ANN | PTB | 0.9632 | 0.8074 | 0.9033 | 0.8944 | 0.9123 | 0.9111 | 0.9023 | 0.9809 | 0.8369 | 0.9184 | 0.9149 | 0.922 | 0.9214 | 0.9181 |
| ANN | PTU | 0.9688 | 0.8307 | 0.9149 | 0.9014 | 0.9283 | 0.9267 | 0.9134 | 0.9788 | 0.8727 | 0.9362 | 0.922 | 0.9504 | 0.9489 | 0.9353 |
| ANN | PTXU | 0.978 | 0.8603 | 0.9292 | 0.914 | 0.9445 | 0.9439 | 0.9279 | 0.983 | 0.8865 | 0.9433 | 0.9433 | 0.9433 | 0.9433 | 0.9433 |
| ANN | PTXLU | 0.8982 | 0.6577 | 0.8279 | 0.8262 | 0.8296 | 0.832 | 0.8278 | 0.9037 | 0.681 | 0.8404 | 0.8511 | 0.8298 | 0.8333 | 0.8421 |

| | | | | | | | | | | | | | | | |
|---|---|---|---|---|---|---|---|---|---|---|---|---|---|---|---|
| ANN | Seq2Vec | 0.9804 | 0.8776 | 0.9382 | 0.9266 | 0.9499 | 0.949 | 0.9371 | 0.9889 | 0.9364 | 0.9681 | 0.9787 | 0.9574 | 0.9583 | 0.9684 |
| ANN | Word2Vec | 0.899 | 0.6804 | 0.8396 | 0.8298 | 0.8494 | 0.8467 | 0.8376 | 0.935 | 0.7317 | 0.8652 | 0.8369 | 0.8936 | 0.8872 | 0.8613 |
| ANN | AAC | 0.8611 | 0.5927 | 0.7949 | 0.7527 | 0.8369 | 0.8235 | 0.7859 | 0.8698 | 0.6171 | 0.8085 | 0.8014 | 0.8156 | 0.8129 | 0.8071 |
| ANN | ACC | 0.6559 | 0.2628 | 0.6307 | 0.6256 | 0.6363 | 0.6318 | 0.6274 | 0.6361 | 0.2201 | 0.6099 | 0.6312 | 0.5887 | 0.6054 | 0.6181 |
| ANN | AC | 0.6477 | 0.2158 | 0.6074 | 0.6022 | 0.6128 | 0.6117 | 0.6059 | 0.6045 | 0.206 | 0.6028 | 0.5745 | 0.6312 | 0.609 | 0.5912 |
| ANN | APAAC | 0.8833 | 0.6185 | 0.8065 | 0.7636 | 0.8495 | 0.8376 | 0.7962 | 0.8938 | 0.6464 | 0.8227 | 0.8511 | 0.7943 | 0.8054 | 0.8276 |
| ANN | ASDC | 0.9295 | 0.7374 | 0.8674 | 0.8584 | 0.8762 | 0.877 | 0.8661 | 0.9543 | 0.7519 | 0.8759 | 0.8652 | 0.8865 | 0.8841 | 0.8746 |
| ANN | CC | 0.6531 | 0.2364 | 0.6174 | 0.6345 | 0.6002 | 0.6174 | 0.6235 | 0.6132 | 0.1631 | 0.5816 | 0.5745 | 0.5887 | 0.5827 | 0.5786 |
| ANN | CKSAAGP-type-1 | 0.9523 | 0.785 | 0.8916 | 0.8799 | 0.9031 | 0.9022 | 0.8899 | 0.986 | 0.9007 | 0.9504 | 0.9504 | 0.9504 | 0.9504 | 0.9504 |
| ANN | CKSAAGP-type-2 | 0.9566 | 0.8033 | 0.9006 | 0.8817 | 0.9194 | 0.9175 | 0.8981 | 0.9651 | 0.8723 | 0.9362 | 0.9362 | 0.9362 | 0.9362 | 0.9362 |
| ANN | CKSAAP-type-1 | 0.9421 | 0.7705 | 0.8844 | 0.8638 | 0.9051 | 0.9013 | 0.8815 | 0.9364 | 0.7664 | 0.883 | 0.8652 | 0.9007 | 0.8971 | 0.8809 |
| ANN | CKSAAP-type-2 | 0.9389 | 0.7676 | 0.8808 | 0.8369 | 0.9249 | 0.92 | 0.8745 | 0.9464 | 0.7692 | 0.883 | 0.8369 | 0.9291 | 0.9219 | 0.8773 |
| ANN | CTDC | 0.8395 | 0.5748 | 0.785 | 0.7618 | 0.8084 | 0.8046 | 0.7797 | 0.8427 | 0.5823 | 0.7908 | 0.766 | 0.8156 | 0.806 | 0.7855 |

| | | | | | | | | | | | | | | | |
|---|---|---|---|---|---|---|---|---|---|---|---|---|---|---|---|
| ANN | CTDD | 0.9735 | 0.8791 | 0.9391 | 0.9391 | 0.939 | 0.9397 | 0.9389 | 0.9715 | 0.9078 | 0.9539 | 0.9504 | 0.9574 | 0.9571 | 0.9537 |
| ANN | CTDT | 0.8804 | 0.6946 | 0.845 | 0.792 | 0.8979 | 0.8859 | 0.8357 | 0.8794 | 0.6526 | 0.8262 | 0.8156 | 0.8369 | 0.8333 | 0.8244 |
| ANN | CTriad | 0.9352 | 0.7609 | 0.8781 | 0.8403 | 0.9159 | 0.9106 | 0.8722 | 0.9385 | 0.7589 | 0.8794 | 0.8723 | 0.8865 | 0.8849 | 0.8786 |
| ANN | DDE | 0.9252 | 0.7439 | 0.871 | 0.853 | 0.8889 | 0.8854 | 0.8679 | 0.9452 | 0.7398 | 0.8688 | 0.8298 | 0.9078 | 0.9 | 0.8635 |
| ANN | DistancePair | 0.8611 | 0.5927 | 0.7949 | 0.7527 | 0.8369 | 0.8235 | 0.7859 | 0.8698 | 0.6171 | 0.8085 | 0.8014 | 0.8156 | 0.8129 | 0.8071 |
| ANN | DPC-type-1 | 0.9451 | 0.789 | 0.8934 | 0.8764 | 0.9104 | 0.909 | 0.8912 | 0.9321 | 0.7039 | 0.8511 | 0.8156 | 0.8865 | 0.8779 | 0.8456 |
| ANN | DPC-type-2 | 0.9392 | 0.7626 | 0.88 | 0.8566 | 0.9033 | 0.9001 | 0.8766 | 0.9427 | 0.7312 | 0.8652 | 0.844 | 0.8865 | 0.8815 | 0.8623 |
| ANN | GDPC-type-1 | 0.9684 | 0.8515 | 0.9248 | 0.9141 | 0.9356 | 0.9352 | 0.9235 | 0.9716 | 0.8865 | 0.9433 | 0.9433 | 0.9433 | 0.9433 | 0.9433 |
| ANN | GDPC-type-2 | 0.9777 | 0.8725 | 0.9355 | 0.9302 | 0.9409 | 0.9413 | 0.9349 | 0.978 | 0.9011 | 0.9504 | 0.9362 | 0.9645 | 0.9635 | 0.9496 |
| ANN | Geary | 0.6368 | 0.2182 | 0.6084 | 0.6057 | 0.6108 | 0.6089 | 0.6051 | 0.5663 | 0.0787 | 0.539 | 0.4752 | 0.6028 | 0.5447 | 0.5076 |
| ANN | GTPC-type-1 | 0.9397 | 0.7726 | 0.8854 | 0.8656 | 0.905 | 0.9013 | 0.8822 | 0.9753 | 0.8595 | 0.9291 | 0.9007 | 0.9574 | 0.9549 | 0.927 |
| ANN | GTPC-type-2 | 0.9504 | 0.7842 | 0.8916 | 0.8746 | 0.9087 | 0.9051 | 0.8893 | 0.9819 | 0.8796 | 0.9397 | 0.9291 | 0.9504 | 0.9493 | 0.9391 |
| ANN | KSCTriad | 0.9296 | 0.7522 | 0.8746 | 0.8495 | 0.8997 | 0.8956 | 0.8706 | 0.9351 | 0.7669 | 0.883 | 0.8582 | 0.9078 | 0.903 | 0.88 |

| | | | | | | | | | | | | | | | |
|---|---|---|---|---|---|---|---|---|---|---|---|---|---|---|---|
| ANN | Moran | 0.6013 | 0.1959 | 0.595 | 0.6686 | 0.5217 | 0.5845 | 0.6198 | 0.582 | 0.1502 | 0.5745 | 0.5106 | 0.6383 | 0.5854 | 0.5455 |
| ANN | NMBroto | 0.6298 | 0.2168 | 0.6066 | 0.6542 | 0.559 | 0.5978 | 0.6218 | 0.5848 | 0.1348 | 0.5674 | 0.5674 | 0.5674 | 0.5674 | 0.5674 |
| ANN | PAAC | 0.8902 | 0.6424 | 0.819 | 0.7831 | 0.8548 | 0.8453 | 0.811 | 0.9283 | 0.7236 | 0.8617 | 0.8511 | 0.8723 | 0.8696 | 0.8602 |
| ANN | PDE | 0.8944 | 0.6664 | 0.8307 | 0.7867 | 0.8745 | 0.8633 | 0.8211 | 0.9273 | 0.7175 | 0.8582 | 0.8865 | 0.8298 | 0.8389 | 0.8621 |
| ANN | PseKRAAC-type-4 | 0.6255 | 0.2216 | 0.6101 | 0.6182 | 0.6021 | 0.6102 | 0.6125 | 0.515 | 0.0213 | 0.5106 | 0.4752 | 0.5461 | 0.5115 | 0.4926 |
| ANN | PseKRAAC-type-5 | 0.8447 | 0.6047 | 0.7922 | 0.6757 | 0.9085 | 0.8857 | 0.762 | 0.8514 | 0.5525 | 0.7695 | 0.6596 | 0.8794 | 0.8455 | 0.741 |
| ANN | PseKRAAC-type-6A | 0.6645 | 0.2416 | 0.6165 | 0.6093 | 0.6238 | 0.6251 | 0.6061 | 0.6765 | 0.2843 | 0.6418 | 0.6099 | 0.6738 | 0.6515 | 0.63 |
| ANN | PseKRAAC-type-13 | 0.6152 | 0.1719 | 0.5852 | 0.5788 | 0.5913 | 0.5884 | 0.5812 | 0.5686 | 0.1207 | 0.5603 | 0.5816 | 0.539 | 0.5578 | 0.5694 |
| ANN | QSOrder | 0.8999 | 0.6486 | 0.8235 | 0.8065 | 0.8406 | 0.8357 | 0.8201 | 0.9137 | 0.6529 | 0.8262 | 0.844 | 0.8085 | 0.8151 | 0.8293 |
| ANN | TPC-type-1 | 0.8369 | 0.5544 | 0.7768 | 0.7849 | 0.7687 | 0.7738 | 0.7788 | 0.9112 | 0.6335 | 0.8156 | 0.773 | 0.8582 | 0.845 | 0.8074 |
| ANN | TPC-type-2 | 0.8441 | 0.5578 | 0.7769 | 0.8226 | 0.7312 | 0.7555 | 0.7865 | 0.915 | 0.6748 | 0.8369 | 0.8085 | 0.8652 | 0.8571 | 0.8321 |
| CB | Bepler | 0.9143 | 0.6855 | 0.8387 | 0.7832 | 0.8942 | 0.8836 | 0.8271 | 0.9312 | 0.7107 | 0.8546 | 0.8227 | 0.8865 | 0.8788 | 0.8498 |
| CB | CPCProt | 0.8989 | 0.6548 | 0.8262 | 0.7956 | 0.8567 | 0.8477 | 0.82 | 0.8913 | 0.6101 | 0.805 | 0.7943 | 0.8156 | 0.8116 | 0.8029 |

| | | | | | | | | | | | | | | | |
|---|---|---|---|---|---|---|---|---|---|---|---|---|---|---|---|
| CB | ESM | 0.9088 | 0.6912 | 0.8449 | 0.8297 | 0.86 | 0.8572 | 0.8425 | 0.9251 | 0.6817 | 0.8404 | 0.8156 | 0.8652 | 0.8582 | 0.8364 |
| CB | ESM1b | 0.8991 | 0.6603 | 0.8289 | 0.8084 | 0.8494 | 0.8453 | 0.8252 | 0.9207 | 0.6891 | 0.844 | 0.8156 | 0.8723 | 0.8647 | 0.8394 |
| CB | ESM1v | 0.8976 | 0.6448 | 0.8217 | 0.8028 | 0.8405 | 0.8344 | 0.8175 | 0.8868 | 0.6254 | 0.8121 | 0.7801 | 0.844 | 0.8333 | 0.8059 |
| CB | ESM2-t6 | 0.8852 | 0.6335 | 0.8154 | 0.8011 | 0.8294 | 0.8266 | 0.812 | 0.8854 | 0.6031 | 0.8014 | 0.7872 | 0.8156 | 0.8102 | 0.7986 |
| CB | ESM2-t12 | 0.9046 | 0.6608 | 0.8289 | 0.7977 | 0.8604 | 0.8506 | 0.8221 | 0.9132 | 0.6391 | 0.8191 | 0.7943 | 0.844 | 0.8358 | 0.8145 |
| CB | ESM2-t30 | 0.8934 | 0.655 | 0.8254 | 0.8085 | 0.8424 | 0.8396 | 0.8211 | 0.9087 | 0.6529 | 0.8262 | 0.8085 | 0.844 | 0.8382 | 0.8231 |
| CB | ESM2-t33 | 0.9163 | 0.7026 | 0.8494 | 0.8206 | 0.8781 | 0.872 | 0.8436 | 0.9169 | 0.7047 | 0.8511 | 0.8085 | 0.8936 | 0.8837 | 0.8444 |
| CB | ESM2-t36 | 0.9179 | 0.7016 | 0.8495 | 0.8262 | 0.8728 | 0.8675 | 0.8451 | 0.9413 | 0.7376 | 0.8688 | 0.8723 | 0.8652 | 0.8662 | 0.8693 |
| CB | FastText | 0.8966 | 0.6649 | 0.8315 | 0.8029 | 0.8602 | 0.8528 | 0.8267 | 0.8838 | 0.5892 | 0.7943 | 0.8156 | 0.773 | 0.7823 | 0.7986 |
| CB | GloVe | 0.8916 | 0.6253 | 0.8118 | 0.8101 | 0.8136 | 0.814 | 0.8111 | 0.8892 | 0.6044 | 0.8014 | 0.766 | 0.8369 | 0.8244 | 0.7941 |
| CB | PRNN | 0.9118 | 0.6675 | 0.8325 | 0.8155 | 0.8495 | 0.845 | 0.8285 | 0.9089 | 0.646 | 0.8227 | 0.8014 | 0.844 | 0.837 | 0.8188 |
| CB | PTAB | 0.8828 | 0.6516 | 0.8253 | 0.8047 | 0.8459 | 0.8398 | 0.8216 | 0.903 | 0.6667 | 0.8333 | 0.8298 | 0.8369 | 0.8357 | 0.8327 |
| CB | PTBB | 0.8773 | 0.5969 | 0.7975 | 0.7904 | 0.8047 | 0.8032 | 0.7955 | 0.903 | 0.646 | 0.8227 | 0.8014 | 0.844 | 0.837 | 0.8188 |

| | | | | | | | | | | | | | | | |
|---|---|---|---|---|---|---|---|---|---|---|---|---|---|---|---|
| CB | PTB | 0.9269 | 0.7146 | 0.8558 | 0.83 | 0.8819 | 0.8766 | 0.8513 | 0.9249 | 0.6667 | 0.8333 | 0.8369 | 0.8298 | 0.831 | 0.8339 |
| CB | PTU | 0.9438 | 0.7557 | 0.8771 | 0.8692 | 0.8851 | 0.8849 | 0.8761 | 0.9484 | 0.7101 | 0.8546 | 0.8298 | 0.8794 | 0.8731 | 0.8509 |
| CB | PTXU | 0.9365 | 0.729 | 0.8638 | 0.8512 | 0.8764 | 0.8741 | 0.8618 | 0.9523 | 0.7459 | 0.8723 | 0.844 | 0.9007 | 0.8947 | 0.8686 |
| CB | PTXLU | 0.8275 | 0.5109 | 0.7545 | 0.7526 | 0.7562 | 0.7569 | 0.7532 | 0.8072 | 0.4898 | 0.7447 | 0.766 | 0.7234 | 0.7347 | 0.75 |
| CB | Seq2Vec | 0.9655 | 0.83 | 0.914 | 0.8854 | 0.9426 | 0.9394 | 0.9112 | 0.9741 | 0.8513 | 0.9255 | 0.9149 | 0.9362 | 0.9348 | 0.9247 |
| CB | Word2Vec | 0.8632 | 0.5946 | 0.7966 | 0.7938 | 0.7992 | 0.7999 | 0.7959 | 0.8962 | 0.662 | 0.8298 | 0.7872 | 0.8723 | 0.8605 | 0.8222 |
| CB | AAC | 0.8849 | 0.6657 | 0.8306 | 0.7848 | 0.8762 | 0.8658 | 0.822 | 0.9183 | 0.6628 | 0.8298 | 0.7801 | 0.8794 | 0.8661 | 0.8209 |
| CB | ACC | 0.6543 | 0.2604 | 0.6298 | 0.629 | 0.6307 | 0.633 | 0.6302 | 0.5956 | 0.1704 | 0.5851 | 0.5603 | 0.6099 | 0.5896 | 0.5745 |
| CB | AC | 0.6263 | 0.2167 | 0.6075 | 0.6219 | 0.5931 | 0.6042 | 0.6113 | 0.6182 | 0.2137 | 0.6064 | 0.5603 | 0.6525 | 0.6172 | 0.5874 |
| CB | APAAC | 0.9101 | 0.6972 | 0.8477 | 0.8332 | 0.862 | 0.8588 | 0.8448 | 0.9091 | 0.7164 | 0.8582 | 0.8652 | 0.8511 | 0.8531 | 0.8592 |
| CB | ASDC | 0.9641 | 0.8044 | 0.9005 | 0.8765 | 0.9247 | 0.9229 | 0.8975 | 0.971 | 0.8232 | 0.9113 | 0.9291 | 0.8936 | 0.8973 | 0.9129 |
| CB | CC | 0.662 | 0.2451 | 0.6218 | 0.6309 | 0.6128 | 0.622 | 0.6247 | 0.6527 | 0.2489 | 0.6241 | 0.6596 | 0.5887 | 0.6159 | 0.637 |
| CB | CKSAAGP-type-1 | 0.9245 | 0.7245 | 0.8611 | 0.8657 | 0.8567 | 0.8598 | 0.8615 | 0.9598 | 0.7589 | 0.8794 | 0.8794 | 0.8794 | 0.8794 | 0.8794 |

| | | | | | | | | | | | | | | | |
|---|---|---|---|---|---|---|---|---|---|---|---|---|---|---|---|
| CB | CKSAAGP-type-2 | 0.942 | 0.7748 | 0.8862 | 0.8889 | 0.8836 | 0.8869 | 0.8865 | 0.9703 | 0.8085 | 0.9043 | 0.9007 | 0.9078 | 0.9071 | 0.9039 |
| CB | CKSAAP-type-1 | 0.9596 | 0.8097 | 0.9006 | 0.837 | 0.9641 | 0.9595 | 0.8922 | 0.964 | 0.8066 | 0.9007 | 0.844 | 0.9574 | 0.952 | 0.8947 |
| CB | CKSAAP-type-2 | 0.9522 | 0.8245 | 0.9068 | 0.8353 | 0.9785 | 0.9761 | 0.898 | 0.97 | 0.8352 | 0.9149 | 0.8582 | 0.9716 | 0.968 | 0.9098 |
| CB | CTDC | 0.8727 | 0.6349 | 0.8164 | 0.7831 | 0.8495 | 0.8388 | 0.8094 | 0.8863 | 0.6667 | 0.8333 | 0.8298 | 0.8369 | 0.8357 | 0.8327 |
| CB | CTDD | 0.9835 | 0.907 | 0.9525 | 0.923 | 0.9821 | 0.981 | 0.9509 | 0.9932 | 0.9231 | 0.961 | 0.9362 | 0.9858 | 0.9851 | 0.96 |
| CB | CTDT | 0.9422 | 0.7936 | 0.8961 | 0.8835 | 0.9088 | 0.9075 | 0.8946 | 0.9489 | 0.7879 | 0.8936 | 0.8723 | 0.9149 | 0.9111 | 0.8913 |
| CB | CTriad | 0.9265 | 0.7576 | 0.8728 | 0.7886 | 0.957 | 0.9487 | 0.86 | 0.9411 | 0.7809 | 0.8865 | 0.8156 | 0.9574 | 0.9504 | 0.8779 |
| CB | DDE | 0.9578 | 0.8276 | 0.9096 | 0.8424 | 0.9768 | 0.9734 | 0.9022 | 0.9665 | 0.8223 | 0.9078 | 0.844 | 0.9716 | 0.9675 | 0.9015 |
| CB | DistancePair | 0.8849 | 0.6657 | 0.8306 | 0.7848 | 0.8762 | 0.8658 | 0.822 | 0.9183 | 0.6628 | 0.8298 | 0.7801 | 0.8794 | 0.8661 | 0.8209 |
| CB | DPC-type-1 | 0.9599 | 0.8415 | 0.9168 | 0.855 | 0.9785 | 0.976 | 0.9102 | 0.974 | 0.843 | 0.9184 | 0.8582 | 0.9787 | 0.9758 | 0.9132 |
| CB | DPC-type-2 | 0.9643 | 0.8664 | 0.9293 | 0.8675 | 0.991 | 0.9903 | 0.9237 | 0.9686 | 0.8547 | 0.9255 | 0.8794 | 0.9716 | 0.9688 | 0.9219 |
| CB | GDPC-type-1 | 0.9502 | 0.7828 | 0.8907 | 0.8924 | 0.8889 | 0.8904 | 0.8906 | 0.9809 | 0.8447 | 0.922 | 0.9433 | 0.9007 | 0.9048 | 0.9236 |
| CB | GDPC-type-2 | 0.9604 | 0.8271 | 0.9131 | 0.9103 | 0.9158 | 0.9155 | 0.9125 | 0.9897 | 0.9008 | 0.9504 | 0.9433 | 0.9574 | 0.9568 | 0.95 |

| | | | | | | | | | | | | | | | |
|---|---|---|---|---|---|---|---|---|---|---|---|---|---|---|---|
| CB | Geary | 0.6222 | 0.1976 | 0.5985 | 0.6076 | 0.5894 | 0.5973 | 0.6015 | 0.5787 | 0.1064 | 0.5532 | 0.5532 | 0.5532 | 0.5532 | 0.5532 |
| CB | GTPC-type-1 | 0.8727 | 0.6312 | 0.8145 | 0.8048 | 0.8243 | 0.8224 | 0.8122 | 0.9322 | 0.7666 | 0.8794 | 0.8085 | 0.9504 | 0.9421 | 0.8702 |
| CB | GTPC-type-2 | 0.9016 | 0.6592 | 0.8289 | 0.8119 | 0.8458 | 0.8411 | 0.8254 | 0.9607 | 0.7876 | 0.8936 | 0.8794 | 0.9078 | 0.9051 | 0.8921 |
| CB | KSCTriad | 0.9306 | 0.7409 | 0.8647 | 0.7832 | 0.9462 | 0.9373 | 0.8518 | 0.9446 | 0.7541 | 0.8759 | 0.8369 | 0.9149 | 0.9077 | 0.8708 |
| CB | Moran | 0.5906 | 0.1733 | 0.586 | 0.5933 | 0.5787 | 0.5858 | 0.5875 | 0.5278 | 0.0356 | 0.5177 | 0.4752 | 0.5603 | 0.5194 | 0.4963 |
| CB | NMBroto | 0.6344 | 0.225 | 0.6119 | 0.6021 | 0.6219 | 0.6168 | 0.608 | 0.5993 | 0.1915 | 0.5957 | 0.5887 | 0.6028 | 0.5971 | 0.5929 |
| CB | PAAC | 0.9186 | 0.6921 | 0.845 | 0.8261 | 0.8637 | 0.8593 | 0.8412 | 0.9333 | 0.7519 | 0.8759 | 0.8865 | 0.8652 | 0.8681 | 0.8772 |
| CB | PDE | 0.9162 | 0.6913 | 0.8432 | 0.8155 | 0.8709 | 0.8658 | 0.8373 | 0.9388 | 0.7243 | 0.8617 | 0.8369 | 0.8865 | 0.8806 | 0.8582 |
| CB | PseKRAAC-type-4 | 0.6136 | 0.1959 | 0.5968 | 0.5702 | 0.6238 | 0.6034 | 0.5833 | 0.5832 | 0.0787 | 0.539 | 0.4752 | 0.6028 | 0.5447 | 0.5076 |
| CB | PseKRAAC-type-5 | 0.8405 | 0.599 | 0.7895 | 0.6722 | 0.9069 | 0.883 | 0.7598 | 0.8455 | 0.5816 | 0.7837 | 0.6738 | 0.8936 | 0.8636 | 0.757 |
| CB | PseKRAAC-type-6A | 0.6584 | 0.2888 | 0.6424 | 0.6182 | 0.6664 | 0.6546 | 0.6312 | 0.6477 | 0.1917 | 0.5957 | 0.5745 | 0.617 | 0.6 | 0.587 |
| CB | PseKRAAC-type-13 | 0.577 | 0.1606 | 0.5797 | 0.5773 | 0.5821 | 0.5829 | 0.5781 | 0.6281 | 0.1846 | 0.5922 | 0.617 | 0.5674 | 0.5878 | 0.6021 |
| CB | QSOrder | 0.9096 | 0.6785 | 0.8369 | 0.8012 | 0.8727 | 0.8655 | 0.8301 | 0.9254 | 0.7164 | 0.8582 | 0.8511 | 0.8652 | 0.8633 | 0.8571 |

| | | | | | | | | | | | | | | | |
|---|---|---|---|---|---|---|---|---|---|---|---|---|---|---|---|
| CB | TPC-type-1 | 0.7491 | 0.4103 | 0.6971 | 0.5664 | 0.8281 | 0.7704 | 0.6512 | 0.8341 | 0.5061 | 0.7447 | 0.617 | 0.8723 | 0.8286 | 0.7073 |
| CB | TPC-type-2 | 0.7815 | 0.4504 | 0.7196 | 0.6202 | 0.8191 | 0.7764 | 0.6867 | 0.8139 | 0.4871 | 0.734 | 0.5957 | 0.8723 | 0.8235 | 0.6914 |
| ERT | Bepler | 0.8515 | 0.5573 | 0.776 | 0.735 | 0.8171 | 0.8002 | 0.7628 | 0.874 | 0.5747 | 0.7872 | 0.773 | 0.8014 | 0.7956 | 0.7842 |
| ERT | CPCProt | 0.8919 | 0.6595 | 0.828 | 0.7974 | 0.8584 | 0.8508 | 0.8217 | 0.8934 | 0.647 | 0.8227 | 0.7872 | 0.8582 | 0.8473 | 0.8162 |
| ERT | ESM | 0.9084 | 0.6956 | 0.8441 | 0.7869 | 0.9015 | 0.8916 | 0.8338 | 0.9159 | 0.6419 | 0.8191 | 0.766 | 0.8723 | 0.8571 | 0.809 |
| ERT | ESM1b | 0.8993 | 0.6881 | 0.8414 | 0.7975 | 0.8854 | 0.8758 | 0.8329 | 0.9269 | 0.6847 | 0.8404 | 0.7872 | 0.8936 | 0.881 | 0.8315 |
| ERT | ESM1v | 0.8883 | 0.6484 | 0.8217 | 0.7742 | 0.8692 | 0.8557 | 0.8112 | 0.9034 | 0.652 | 0.8227 | 0.7518 | 0.8936 | 0.876 | 0.8092 |
| ERT | ESM2-t6 | 0.8864 | 0.643 | 0.8199 | 0.7903 | 0.8496 | 0.8417 | 0.8138 | 0.9007 | 0.5892 | 0.7943 | 0.773 | 0.8156 | 0.8074 | 0.7899 |
| ERT | ESM2-t12 | 0.8922 | 0.6589 | 0.8279 | 0.7975 | 0.8583 | 0.8511 | 0.822 | 0.9132 | 0.6277 | 0.8121 | 0.7589 | 0.8652 | 0.8492 | 0.8015 |
| ERT | ESM2-t30 | 0.8905 | 0.6631 | 0.8288 | 0.7958 | 0.8618 | 0.8549 | 0.821 | 0.9006 | 0.6628 | 0.8298 | 0.7801 | 0.8794 | 0.8661 | 0.8209 |
| ERT | ESM2-t33 | 0.9124 | 0.703 | 0.8486 | 0.7957 | 0.9015 | 0.8897 | 0.8383 | 0.9214 | 0.6965 | 0.8475 | 0.8156 | 0.8794 | 0.8712 | 0.8425 |
| ERT | ESM2-t36 | 0.9088 | 0.6897 | 0.8396 | 0.7633 | 0.9157 | 0.9037 | 0.8255 | 0.9227 | 0.6573 | 0.8262 | 0.766 | 0.8865 | 0.871 | 0.8151 |
| ERT | FastText | 0.8769 | 0.624 | 0.8101 | 0.7725 | 0.8478 | 0.8375 | 0.8021 | 0.8847 | 0.6201 | 0.8085 | 0.7589 | 0.8582 | 0.8425 | 0.7985 |

| | | | | | | | | | | | | | | | |
|---|---|---|---|---|---|---|---|---|---|---|---|---|---|---|---|
| ERT | GloVe | 0.8721 | 0.5892 | 0.7939 | 0.7903 | 0.7975 | 0.7987 | 0.7935 | 0.8846 | 0.5817 | 0.7908 | 0.7801 | 0.8014 | 0.7971 | 0.7885 |
| ERT | PRNN | 0.8562 | 0.5697 | 0.783 | 0.7597 | 0.8065 | 0.8001 | 0.7769 | 0.8785 | 0.609 | 0.8014 | 0.7305 | 0.8723 | 0.8512 | 0.7863 |
| ERT | PTAB | 0.8737 | 0.6224 | 0.8099 | 0.7938 | 0.8262 | 0.8217 | 0.806 | 0.8918 | 0.6396 | 0.8191 | 0.7872 | 0.8511 | 0.8409 | 0.8132 |
| ERT | PTBB | 0.8636 | 0.586 | 0.7921 | 0.7652 | 0.819 | 0.8089 | 0.7857 | 0.9043 | 0.6809 | 0.8404 | 0.8369 | 0.844 | 0.8429 | 0.8399 |
| ERT | PTB | 0.9054 | 0.6702 | 0.8325 | 0.785 | 0.8799 | 0.8685 | 0.8225 | 0.9283 | 0.6914 | 0.844 | 0.7943 | 0.8936 | 0.8819 | 0.8358 |
| ERT | PTU | 0.9293 | 0.7236 | 0.8602 | 0.8332 | 0.8872 | 0.8822 | 0.8555 | 0.9205 | 0.6936 | 0.844 | 0.7801 | 0.9078 | 0.8943 | 0.8333 |
| ERT | PTXU | 0.9323 | 0.7184 | 0.8575 | 0.8297 | 0.8853 | 0.8804 | 0.8528 | 0.9317 | 0.7341 | 0.8652 | 0.8156 | 0.9149 | 0.9055 | 0.8582 |
| ERT | PTXLU | 0.7712 | 0.4329 | 0.716 | 0.7293 | 0.7026 | 0.711 | 0.7194 | 0.765 | 0.3699 | 0.6844 | 0.7234 | 0.6454 | 0.6711 | 0.6962 |
| ERT | Seq2Vec | 0.9451 | 0.7758 | 0.8862 | 0.8531 | 0.9194 | 0.9155 | 0.8821 | 0.9584 | 0.7664 | 0.883 | 0.8652 | 0.9007 | 0.8971 | 0.8809 |
| ERT | Word2Vec | 0.862 | 0.5839 | 0.7903 | 0.767 | 0.8136 | 0.8077 | 0.785 | 0.8982 | 0.6201 | 0.8085 | 0.7589 | 0.8582 | 0.8425 | 0.7985 |
| ERT | AAC | 0.895 | 0.6535 | 0.8253 | 0.7831 | 0.8673 | 0.8556 | 0.8173 | 0.9247 | 0.7317 | 0.8652 | 0.8369 | 0.8936 | 0.8872 | 0.8613 |
| ERT | ACC | 0.6608 | 0.2591 | 0.629 | 0.6041 | 0.6542 | 0.6353 | 0.6185 | 0.6274 | 0.1782 | 0.5887 | 0.539 | 0.6383 | 0.5984 | 0.5672 |
| ERT | AC | 0.6387 | 0.2525 | 0.6255 | 0.6077 | 0.6435 | 0.6292 | 0.6162 | 0.6085 | 0.1874 | 0.5922 | 0.5035 | 0.6809 | 0.6121 | 0.5525 |

| | | | | | | | | | | | | | | | |
|---|---|---|---|---|---|---|---|---|---|---|---|---|---|---|---|
| ERT | APAAC | 0.9214 | 0.6912 | 0.8432 | 0.8082 | 0.8781 | 0.8716 | 0.8364 | 0.936 | 0.7589 | 0.8794 | 0.8794 | 0.8794 | 0.8794 | 0.8794 |
| ERT | ASDC | 0.9403 | 0.7887 | 0.8925 | 0.853 | 0.9318 | 0.9269 | 0.8875 | 0.9565 | 0.8054 | 0.9007 | 0.8511 | 0.9504 | 0.9449 | 0.8955 |
| ERT | CC | 0.6499 | 0.2416 | 0.6201 | 0.6185 | 0.622 | 0.6199 | 0.6172 | 0.6199 | 0.2129 | 0.6064 | 0.5887 | 0.6241 | 0.6103 | 0.5993 |
| ERT | CKSAAGP-type-1 | 0.8965 | 0.6714 | 0.8342 | 0.8369 | 0.8316 | 0.8368 | 0.835 | 0.9358 | 0.6951 | 0.8475 | 0.8511 | 0.844 | 0.8451 | 0.8481 |
| ERT | CKSAAGP-type-2 | 0.9067 | 0.6695 | 0.8334 | 0.8389 | 0.828 | 0.833 | 0.8343 | 0.9519 | 0.7448 | 0.8723 | 0.8652 | 0.8794 | 0.8777 | 0.8714 |
| ERT | CKSAAP-type-1 | 0.9503 | 0.8051 | 0.8997 | 0.8459 | 0.9535 | 0.9486 | 0.8935 | 0.9573 | 0.7977 | 0.8972 | 0.8511 | 0.9433 | 0.9375 | 0.8922 |
| ERT | CKSAAP-type-2 | 0.9472 | 0.835 | 0.9149 | 0.8675 | 0.9624 | 0.9594 | 0.9101 | 0.9551 | 0.8177 | 0.9078 | 0.8723 | 0.9433 | 0.9389 | 0.9044 |
| ERT | CTDC | 0.8604 | 0.5932 | 0.7948 | 0.7454 | 0.8442 | 0.827 | 0.7834 | 0.8846 | 0.6268 | 0.8121 | 0.766 | 0.8582 | 0.8438 | 0.803 |
| ERT | CTDD | 0.9789 | 0.8794 | 0.9391 | 0.9212 | 0.957 | 0.9561 | 0.9379 | 0.9807 | 0.9011 | 0.9504 | 0.9362 | 0.9645 | 0.9635 | 0.9496 |
| ERT | CTDT | 0.9095 | 0.7332 | 0.863 | 0.8047 | 0.9212 | 0.9121 | 0.853 | 0.9245 | 0.7157 | 0.8546 | 0.7872 | 0.922 | 0.9098 | 0.8441 |
| ERT | CTriad | 0.9373 | 0.8011 | 0.8961 | 0.8316 | 0.9606 | 0.9563 | 0.8878 | 0.9465 | 0.7746 | 0.883 | 0.8085 | 0.9574 | 0.95 | 0.8736 |
| ERT | DDE | 0.954 | 0.8248 | 0.9087 | 0.8496 | 0.9677 | 0.9648 | 0.9022 | 0.9681 | 0.8303 | 0.9113 | 0.844 | 0.9787 | 0.9754 | 0.9049 |
| ERT | DistancePair | 0.895 | 0.6535 | 0.8253 | 0.7831 | 0.8673 | 0.8556 | 0.8173 | 0.9247 | 0.7317 | 0.8652 | 0.8369 | 0.8936 | 0.8872 | 0.8613 |

| | | | | | | | | | | | | | | | |
|---|---|---|---|---|---|---|---|---|---|---|---|---|---|---|---|
| ERT | DPC-type-1 | 0.9575 | 0.8264 | 0.9104 | 0.8568 | 0.9642 | 0.9602 | 0.905 | 0.9708 | 0.8352 | 0.9149 | 0.8582 | 0.9716 | 0.968 | 0.9098 |
| ERT | DPC-type-2 | 0.96 | 0.8476 | 0.9212 | 0.8711 | 0.9714 | 0.9686 | 0.9165 | 0.9654 | 0.8689 | 0.9326 | 0.8865 | 0.9787 | 0.9766 | 0.9294 |
| ERT | GDPC-type-1 | 0.8969 | 0.662 | 0.8289 | 0.8424 | 0.8153 | 0.8243 | 0.831 | 0.9458 | 0.7381 | 0.8688 | 0.8511 | 0.8865 | 0.8824 | 0.8664 |
| ERT | GDPC-type-2 | 0.9029 | 0.6646 | 0.8307 | 0.821 | 0.8404 | 0.8392 | 0.828 | 0.9552 | 0.7743 | 0.8865 | 0.8582 | 0.9149 | 0.9098 | 0.8832 |
| ERT | Geary | 0.6315 | 0.2354 | 0.6174 | 0.6021 | 0.6326 | 0.6217 | 0.611 | 0.5918 | 0.156 | 0.578 | 0.5816 | 0.5745 | 0.5775 | 0.5795 |
| ERT | GTPC-type-1 | 0.8851 | 0.6547 | 0.8262 | 0.8209 | 0.8316 | 0.8318 | 0.8249 | 0.9258 | 0.7175 | 0.8582 | 0.8298 | 0.8865 | 0.8797 | 0.854 |
| ERT | GTPC-type-2 | 0.8765 | 0.6508 | 0.8245 | 0.8264 | 0.8227 | 0.8241 | 0.8241 | 0.925 | 0.7028 | 0.8511 | 0.8298 | 0.8723 | 0.8667 | 0.8478 |
| ERT | KSCTriad | 0.9306 | 0.7764 | 0.8835 | 0.8119 | 0.9553 | 0.949 | 0.874 | 0.9257 | 0.7093 | 0.8511 | 0.7801 | 0.922 | 0.9091 | 0.8397 |
| ERT | Moran | 0.6115 | 0.2084 | 0.604 | 0.5951 | 0.6129 | 0.6054 | 0.5996 | 0.5531 | 0.0501 | 0.5248 | 0.461 | 0.5887 | 0.5285 | 0.4924 |
| ERT | NMBroto | 0.6553 | 0.2612 | 0.6298 | 0.62 | 0.6397 | 0.6325 | 0.6241 | 0.6262 | 0.1703 | 0.5851 | 0.5674 | 0.6028 | 0.5882 | 0.5776 |
| ERT | PAAC | 0.9087 | 0.6854 | 0.8414 | 0.8046 | 0.8781 | 0.8683 | 0.8346 | 0.9164 | 0.6959 | 0.8475 | 0.8227 | 0.8723 | 0.8657 | 0.8436 |
| ERT | PDE | 0.8972 | 0.6461 | 0.819 | 0.7561 | 0.8814 | 0.868 | 0.8054 | 0.9085 | 0.641 | 0.8191 | 0.773 | 0.8652 | 0.8516 | 0.8104 |
| ERT | PseKRAAC-type-4 | 0.6269 | 0.2081 | 0.603 | 0.6094 | 0.5967 | 0.6031 | 0.6034 | 0.5564 | 0.0568 | 0.5284 | 0.5106 | 0.5461 | 0.5294 | 0.5199 |

| | | | | | | | | | | | | | | | |
|---|---|---|---|---|---|---|---|---|---|---|---|---|---|---|---|
| ERT | PseKRAAC-type-5 | 0.8238 | 0.5242 | 0.759 | 0.7063 | 0.8118 | 0.7936 | 0.7444 | 0.8217 | 0.5001 | 0.7482 | 0.6879 | 0.8085 | 0.7823 | 0.7321 |
| ERT | PseKRAAC-type-6A | 0.6519 | 0.2486 | 0.6237 | 0.6328 | 0.6147 | 0.6212 | 0.6255 | 0.6975 | 0.2982 | 0.6489 | 0.6738 | 0.6241 | 0.6419 | 0.6574 |
| ERT | PseKRAAC-type-13 | 0.5696 | 0.1115 | 0.5555 | 0.5645 | 0.5463 | 0.5575 | 0.5595 | 0.6029 | 0.1633 | 0.5816 | 0.6028 | 0.5603 | 0.5782 | 0.5903 |
| ERT | QSOrder | 0.904 | 0.6718 | 0.8342 | 0.794 | 0.8745 | 0.8641 | 0.8266 | 0.9232 | 0.6972 | 0.8475 | 0.8085 | 0.8865 | 0.8769 | 0.8413 |
| ERT | TPC-type-1 | 0.8741 | 0.6048 | 0.7921 | 0.6683 | 0.9157 | 0.8901 | 0.7605 | 0.9263 | 0.7188 | 0.8546 | 0.773 | 0.9362 | 0.9237 | 0.8417 |
| ERT | TPC-type-2 | 0.8645 | 0.5962 | 0.7859 | 0.647 | 0.9247 | 0.8966 | 0.7494 | 0.9224 | 0.6675 | 0.8227 | 0.695 | 0.9504 | 0.9333 | 0.7967 |
| GB | Bepler | 0.8794 | 0.5955 | 0.7966 | 0.7796 | 0.8135 | 0.8076 | 0.7919 | 0.9133 | 0.6667 | 0.8333 | 0.8369 | 0.8298 | 0.831 | 0.8339 |
| GB | CPCProt | 0.895 | 0.6607 | 0.8289 | 0.7939 | 0.8637 | 0.8545 | 0.8221 | 0.8958 | 0.6335 | 0.8156 | 0.773 | 0.8582 | 0.845 | 0.8074 |
| GB | ESM | 0.9173 | 0.7144 | 0.8539 | 0.794 | 0.9138 | 0.9037 | 0.8441 | 0.9312 | 0.6762 | 0.8369 | 0.7943 | 0.8794 | 0.8682 | 0.8296 |
| GB | ESM1b | 0.9048 | 0.6923 | 0.8441 | 0.7994 | 0.8889 | 0.8789 | 0.8362 | 0.9188 | 0.6671 | 0.8333 | 0.8156 | 0.8511 | 0.8456 | 0.8303 |
| GB | ESM1v | 0.8973 | 0.6664 | 0.8315 | 0.7922 | 0.871 | 0.861 | 0.8243 | 0.9111 | 0.6442 | 0.8191 | 0.7518 | 0.8865 | 0.8689 | 0.8061 |
| GB | ESM2-t6 | 0.8865 | 0.641 | 0.819 | 0.7885 | 0.8495 | 0.8413 | 0.8128 | 0.8956 | 0.6118 | 0.805 | 0.766 | 0.844 | 0.8308 | 0.797 |
| GB | ESM2-t12 | 0.9025 | 0.6827 | 0.8396 | 0.8155 | 0.8638 | 0.8587 | 0.8347 | 0.9257 | 0.6838 | 0.8404 | 0.7943 | 0.8865 | 0.875 | 0.8327 |

| | | | | | | | | | | | | | | | |
|---|---|---|---|---|---|---|---|---|---|---|---|---|---|---|---|
| GB | ESM2-t30 | 0.9053 | 0.6817 | 0.8388 | 0.8065 | 0.8708 | 0.8631 | 0.8315 | 0.9173 | 0.7123 | 0.8546 | 0.8085 | 0.9007 | 0.8906 | 0.8476 |
| GB | ESM2-t33 | 0.9234 | 0.7316 | 0.8629 | 0.8171 | 0.9086 | 0.901 | 0.8549 | 0.9253 | 0.7249 | 0.8617 | 0.8298 | 0.8936 | 0.8864 | 0.8571 |
| GB | ESM2-t36 | 0.9225 | 0.7197 | 0.8557 | 0.7902 | 0.921 | 0.9119 | 0.845 | 0.9417 | 0.7301 | 0.8617 | 0.7943 | 0.9291 | 0.918 | 0.8517 |
| GB | FastText | 0.8911 | 0.6536 | 0.8253 | 0.7886 | 0.862 | 0.8522 | 0.8183 | 0.9096 | 0.6538 | 0.8262 | 0.7943 | 0.8582 | 0.8485 | 0.8205 |
| GB | GloVe | 0.8846 | 0.6208 | 0.8101 | 0.8029 | 0.8173 | 0.8149 | 0.8085 | 0.8864 | 0.6241 | 0.8121 | 0.8156 | 0.8085 | 0.8099 | 0.8127 |
| GB | PRNN | 0.8858 | 0.6171 | 0.8072 | 0.7849 | 0.8297 | 0.8228 | 0.8018 | 0.9056 | 0.6904 | 0.844 | 0.8014 | 0.8865 | 0.876 | 0.837 |
| GB | PTAB | 0.8806 | 0.6345 | 0.8163 | 0.8154 | 0.8172 | 0.8182 | 0.8156 | 0.8962 | 0.6099 | 0.805 | 0.8014 | 0.8085 | 0.8071 | 0.8043 |
| GB | PTBB | 0.8762 | 0.6006 | 0.7993 | 0.7743 | 0.8244 | 0.816 | 0.7937 | 0.9226 | 0.7039 | 0.8511 | 0.8156 | 0.8865 | 0.8779 | 0.8456 |
| GB | PTB | 0.9111 | 0.6929 | 0.8441 | 0.7994 | 0.8889 | 0.8787 | 0.8356 | 0.9401 | 0.7045 | 0.8475 | 0.766 | 0.9291 | 0.9153 | 0.834 |
| GB | PTU | 0.934 | 0.7305 | 0.8629 | 0.8316 | 0.8943 | 0.8906 | 0.8579 | 0.9221 | 0.663 | 0.8262 | 0.7376 | 0.9149 | 0.8966 | 0.8093 |
| GB | PTXU | 0.9396 | 0.732 | 0.8638 | 0.8244 | 0.9032 | 0.897 | 0.8576 | 0.9389 | 0.7133 | 0.8546 | 0.8014 | 0.9078 | 0.8968 | 0.8464 |
| GB | PTXLU | 0.7987 | 0.4649 | 0.7321 | 0.7276 | 0.7366 | 0.7352 | 0.7308 | 0.8047 | 0.4403 | 0.7199 | 0.695 | 0.7447 | 0.7313 | 0.7127 |
| GB | Seq2Vec | 0.9551 | 0.7983 | 0.8978 | 0.8674 | 0.9283 | 0.9244 | 0.8941 | 0.9652 | 0.7953 | 0.8972 | 0.8723 | 0.922 | 0.9179 | 0.8945 |

| | | | | | | | | | | | | | | | |
|---|---|---|---|---|---|---|---|---|---|---|---|---|---|---|---|
| GB | Word2Vec | 0.8768 | 0.609 | 0.8028 | 0.7742 | 0.8315 | 0.8236 | 0.7965 | 0.9039 | 0.647 | 0.8227 | 0.7872 | 0.8582 | 0.8473 | 0.8162 |
| GB | AAC | 0.884 | 0.6089 | 0.8029 | 0.7597 | 0.8458 | 0.8307 | 0.7928 | 0.9194 | 0.6891 | 0.844 | 0.8156 | 0.8723 | 0.8647 | 0.8394 |
| GB | ACC | 0.6827 | 0.2755 | 0.6371 | 0.6184 | 0.6558 | 0.6423 | 0.6284 | 0.6562 | 0.2553 | 0.6277 | 0.6312 | 0.6241 | 0.6268 | 0.629 |
| GB | AC | 0.6331 | 0.2318 | 0.6156 | 0.6236 | 0.6075 | 0.6157 | 0.6189 | 0.629 | 0.1865 | 0.5922 | 0.5177 | 0.6667 | 0.6083 | 0.5594 |
| GB | APAAC | 0.8942 | 0.6323 | 0.8146 | 0.7762 | 0.8531 | 0.8406 | 0.806 | 0.917 | 0.7101 | 0.8546 | 0.8298 | 0.8794 | 0.8731 | 0.8509 |
| GB | ASDC | 0.9601 | 0.7962 | 0.897 | 0.8889 | 0.905 | 0.9054 | 0.8957 | 0.9739 | 0.8021 | 0.9007 | 0.922 | 0.8794 | 0.8844 | 0.9028 |
| GB | CC | 0.6613 | 0.2555 | 0.6085 | 0.3479 | 0.8692 | 0.7296 | 0.4696 | 0.6483 | 0.268 | 0.6135 | 0.3475 | 0.8794 | 0.7424 | 0.4734 |
| GB | CKSAAGP-type-1 | 0.9102 | 0.6824 | 0.8396 | 0.8603 | 0.8189 | 0.8291 | 0.8428 | 0.9528 | 0.7589 | 0.8794 | 0.8723 | 0.8865 | 0.8849 | 0.8786 |
| GB | CKSAAGP-type-2 | 0.9164 | 0.6769 | 0.837 | 0.8388 | 0.8351 | 0.8387 | 0.8369 | 0.9667 | 0.7945 | 0.8972 | 0.8865 | 0.9078 | 0.9058 | 0.8961 |
| GB | CKSAAP-type-1 | 0.959 | 0.8267 | 0.9113 | 0.8729 | 0.9498 | 0.9469 | 0.9074 | 0.9739 | 0.8379 | 0.9184 | 0.8936 | 0.9433 | 0.9403 | 0.9164 |
| GB | CKSAAP-type-2 | 0.9569 | 0.8358 | 0.9149 | 0.8639 | 0.966 | 0.9628 | 0.9094 | 0.9666 | 0.8186 | 0.9078 | 0.8652 | 0.9504 | 0.9457 | 0.9037 |
| GB | CTDC | 0.8682 | 0.6145 | 0.8065 | 0.7742 | 0.8388 | 0.8273 | 0.7996 | 0.9009 | 0.6891 | 0.844 | 0.8156 | 0.8723 | 0.8647 | 0.8394 |
| GB | CTDD | 0.9884 | 0.8796 | 0.9373 | 0.8872 | 0.9875 | 0.9861 | 0.9336 | 0.9861 | 0.8963 | 0.9468 | 0.9078 | 0.9858 | 0.9846 | 0.9446 |

| | | | | | | | | | | | | | | | |
|---|---|---|---|---|---|---|---|---|---|---|---|---|---|---|---|
| GB | CTDT | 0.9257 | 0.7483 | 0.8719 | 0.8244 | 0.9195 | 0.9117 | 0.8649 | 0.9084 | 0.7181 | 0.8582 | 0.8227 | 0.8936 | 0.8855 | 0.8529 |
| GB | CTriad | 0.9379 | 0.8071 | 0.8997 | 0.8369 | 0.9624 | 0.9575 | 0.8919 | 0.9525 | 0.773 | 0.883 | 0.8156 | 0.9504 | 0.9426 | 0.8745 |
| GB | DDE | 0.9522 | 0.8053 | 0.9006 | 0.864 | 0.9373 | 0.9329 | 0.8958 | 0.9646 | 0.808 | 0.9007 | 0.8369 | 0.9645 | 0.9593 | 0.8939 |
| GB | DistancePair | 0.884 | 0.6089 | 0.8029 | 0.7597 | 0.8458 | 0.8307 | 0.7928 | 0.9194 | 0.6891 | 0.844 | 0.8156 | 0.8723 | 0.8647 | 0.8394 |
| GB | DPC-type-1 | 0.9634 | 0.8447 | 0.9203 | 0.8818 | 0.9588 | 0.9561 | 0.9163 | 0.9817 | 0.8832 | 0.9397 | 0.8936 | 0.9858 | 0.9844 | 0.9368 |
| GB | DPC-type-2 | 0.9626 | 0.8565 | 0.9257 | 0.8765 | 0.975 | 0.9727 | 0.9213 | 0.9783 | 0.8767 | 0.9362 | 0.8865 | 0.9858 | 0.9843 | 0.9328 |
| GB | GDPC-type-1 | 0.9005 | 0.6596 | 0.828 | 0.8388 | 0.817 | 0.8249 | 0.8297 | 0.9502 | 0.7601 | 0.8794 | 0.8511 | 0.9078 | 0.9023 | 0.8759 |
| GB | GDPC-type-2 | 0.9178 | 0.7064 | 0.8522 | 0.8604 | 0.8441 | 0.8477 | 0.853 | 0.9702 | 0.7953 | 0.8972 | 0.8723 | 0.922 | 0.9179 | 0.8945 |
| GB | Geary | 0.6277 | 0.207 | 0.6031 | 0.5932 | 0.613 | 0.6048 | 0.5981 | 0.5624 | 0.1077 | 0.5532 | 0.4752 | 0.6312 | 0.563 | 0.5154 |
| GB | GTPC-type-1 | 0.8784 | 0.6348 | 0.8163 | 0.8209 | 0.8117 | 0.8172 | 0.8176 | 0.9277 | 0.7407 | 0.8688 | 0.8227 | 0.9149 | 0.9062 | 0.8625 |
| GB | GTPC-type-2 | 0.8819 | 0.6467 | 0.82 | 0.7619 | 0.8781 | 0.8629 | 0.8073 | 0.943 | 0.7316 | 0.8617 | 0.7872 | 0.9362 | 0.925 | 0.8506 |
| GB | KSCTriad | 0.9369 | 0.7807 | 0.8871 | 0.8298 | 0.9445 | 0.9383 | 0.8797 | 0.9407 | 0.755 | 0.8759 | 0.8298 | 0.922 | 0.9141 | 0.8699 |
| GB | Moran | 0.6054 | 0.1854 | 0.5924 | 0.5951 | 0.5898 | 0.5918 | 0.5925 | 0.5466 | 0.114 | 0.5567 | 0.5106 | 0.6028 | 0.5625 | 0.5353 |

| | | | | | | | | | | | | | | | |
|---|---|---|---|---|---|---|---|---|---|---|---|---|---|---|---|
| GB | NMBroto | 0.6571 | 0.2516 | 0.6253 | 0.6164 | 0.6343 | 0.6283 | 0.6213 | 0.6171 | 0.2058 | 0.6028 | 0.5887 | 0.617 | 0.6058 | 0.5971 |
| GB | PAAC | 0.8956 | 0.6394 | 0.8181 | 0.776 | 0.8602 | 0.8476 | 0.8094 | 0.902 | 0.6822 | 0.8404 | 0.8085 | 0.8723 | 0.8636 | 0.8352 |
| GB | PDE | 0.8961 | 0.6362 | 0.8154 | 0.774 | 0.8565 | 0.8462 | 0.8059 | 0.9184 | 0.6533 | 0.8262 | 0.8014 | 0.8511 | 0.8433 | 0.8218 |
| GB | PseKRAAC-type-4 | 0.6288 | 0.204 | 0.6012 | 0.6023 | 0.6003 | 0.6051 | 0.6016 | 0.5503 | 0.0426 | 0.5213 | 0.4965 | 0.5461 | 0.5224 | 0.5091 |
| GB | PseKRAAC-type-5 | 0.8162 | 0.516 | 0.7554 | 0.6972 | 0.8136 | 0.7914 | 0.7399 | 0.8164 | 0.4867 | 0.7411 | 0.6738 | 0.8085 | 0.7787 | 0.7224 |
| GB | PseKRAAC-type-6A | 0.64 | 0.234 | 0.6164 | 0.6094 | 0.6235 | 0.6192 | 0.6127 | 0.6917 | 0.2784 | 0.6383 | 0.5816 | 0.695 | 0.656 | 0.6165 |
| GB | PseKRAAC-type-13 | 0.5811 | 0.1426 | 0.5707 | 0.5612 | 0.5805 | 0.5741 | 0.5662 | 0.605 | 0.1071 | 0.5532 | 0.6099 | 0.4965 | 0.5478 | 0.5772 |
| GB | QSOrder | 0.8894 | 0.6354 | 0.8154 | 0.7778 | 0.853 | 0.8431 | 0.807 | 0.9148 | 0.7028 | 0.8511 | 0.8298 | 0.8723 | 0.8667 | 0.8478 |
| GB | TPC-type-1 | 0.837 | 0.5601 | 0.7777 | 0.7275 | 0.828 | 0.8099 | 0.7647 | 0.8842 | 0.5916 | 0.7943 | 0.7447 | 0.844 | 0.8268 | 0.7836 |
| GB | TPC-type-2 | 0.8668 | 0.5919 | 0.7904 | 0.6953 | 0.8855 | 0.858 | 0.7672 | 0.9166 | 0.6702 | 0.8227 | 0.6879 | 0.9574 | 0.9417 | 0.7951 |
| LGB | Bepler | 0.9079 | 0.6795 | 0.8369 | 0.8047 | 0.869 | 0.8646 | 0.8304 | 0.9205 | 0.7107 | 0.8546 | 0.8227 | 0.8865 | 0.8788 | 0.8498 |
| LGB | CPCProt | 0.889 | 0.6417 | 0.8181 | 0.7705 | 0.8656 | 0.8549 | 0.8087 | 0.9115 | 0.7144 | 0.8546 | 0.7943 | 0.9149 | 0.9032 | 0.8453 |
| LGB | ESM | 0.9124 | 0.6894 | 0.8432 | 0.81 | 0.8761 | 0.8696 | 0.8375 | 0.9321 | 0.6847 | 0.8404 | 0.7872 | 0.8936 | 0.881 | 0.8315 |

| | | | | | | | | | | | | | | | |
|---|---|---|---|---|---|---|---|---|---|---|---|---|---|---|---|
| LGB | ESM1b | 0.9054 | 0.6676 | 0.8324 | 0.7994 | 0.8656 | 0.8566 | 0.8266 | 0.9128 | 0.6562 | 0.8262 | 0.773 | 0.8794 | 0.8651 | 0.8165 |
| LGB | ESM1v | 0.8841 | 0.6312 | 0.8145 | 0.7815 | 0.8478 | 0.8364 | 0.8074 | 0.9029 | 0.6545 | 0.8262 | 0.7872 | 0.8652 | 0.8538 | 0.8192 |
| LGB | ESM2-t6 | 0.8797 | 0.5942 | 0.7957 | 0.7633 | 0.8278 | 0.8175 | 0.7881 | 0.8972 | 0.6545 | 0.8262 | 0.7872 | 0.8652 | 0.8538 | 0.8192 |
| LGB | ESM2-t12 | 0.8952 | 0.6322 | 0.8154 | 0.7958 | 0.8351 | 0.8291 | 0.8115 | 0.9156 | 0.6886 | 0.844 | 0.8227 | 0.8652 | 0.8593 | 0.8406 |
| LGB | ESM2-t30 | 0.8817 | 0.6293 | 0.8137 | 0.7885 | 0.8388 | 0.8311 | 0.8083 | 0.9131 | 0.6687 | 0.8333 | 0.7943 | 0.8723 | 0.8615 | 0.8266 |
| LGB | ESM2-t33 | 0.9078 | 0.6847 | 0.8405 | 0.803 | 0.8781 | 0.8691 | 0.8334 | 0.8832 | 0.641 | 0.8191 | 0.773 | 0.8652 | 0.8516 | 0.8104 |
| LGB | ESM2-t36 | 0.9028 | 0.66 | 0.828 | 0.7886 | 0.8674 | 0.8577 | 0.8202 | 0.9215 | 0.7097 | 0.8546 | 0.8369 | 0.8723 | 0.8676 | 0.852 |
| LGB | FastText | 0.8926 | 0.6622 | 0.8288 | 0.7778 | 0.88 | 0.8677 | 0.8195 | 0.8895 | 0.6813 | 0.8404 | 0.8227 | 0.8582 | 0.8529 | 0.8375 |
| LGB | GloVe | 0.878 | 0.6193 | 0.8092 | 0.7956 | 0.8226 | 0.8184 | 0.8063 | 0.893 | 0.6103 | 0.805 | 0.7872 | 0.8227 | 0.8162 | 0.8014 |
| LGB | PRNN | 0.8947 | 0.6592 | 0.828 | 0.7867 | 0.8693 | 0.8569 | 0.8194 | 0.9216 | 0.6585 | 0.8262 | 0.7589 | 0.8936 | 0.877 | 0.8137 |
| LGB | PTAB | 0.8893 | 0.6331 | 0.8154 | 0.8047 | 0.8263 | 0.8244 | 0.813 | 0.9117 | 0.6245 | 0.8121 | 0.7943 | 0.8298 | 0.8235 | 0.8087 |
| LGB | PTBB | 0.8776 | 0.6006 | 0.7993 | 0.7779 | 0.8209 | 0.8142 | 0.7946 | 0.9123 | 0.6671 | 0.8333 | 0.8511 | 0.8156 | 0.8219 | 0.8362 |
| LGB | PTB | 0.924 | 0.7027 | 0.8495 | 0.8155 | 0.8836 | 0.8771 | 0.8437 | 0.9315 | 0.6924 | 0.844 | 0.7872 | 0.9007 | 0.888 | 0.8346 |

| | | | | | | | | | | | | | | | |
|---|---|---|---|---|---|---|---|---|---|---|---|---|---|---|---|
| LGB | PTU | 0.9453 | 0.7582 | 0.8772 | 0.8495 | 0.905 | 0.9021 | 0.8735 | 0.9368 | 0.6829 | 0.8404 | 0.8014 | 0.8794 | 0.8692 | 0.8339 |
| LGB | PTXU | 0.9377 | 0.7287 | 0.8629 | 0.8333 | 0.8926 | 0.8872 | 0.8584 | 0.9356 | 0.7181 | 0.8582 | 0.8227 | 0.8936 | 0.8855 | 0.8529 |
| LGB | PTXLU | 0.8122 | 0.4971 | 0.7482 | 0.7456 | 0.7508 | 0.7503 | 0.7474 | 0.8288 | 0.4824 | 0.7411 | 0.7518 | 0.7305 | 0.7361 | 0.7439 |
| LGB | Seq2Vec | 0.9529 | 0.7824 | 0.8898 | 0.862 | 0.9176 | 0.9147 | 0.8865 | 0.9674 | 0.7945 | 0.8972 | 0.8865 | 0.9078 | 0.9058 | 0.8961 |
| LGB | AAC | 0.887 | 0.6428 | 0.819 | 0.7777 | 0.8601 | 0.8503 | 0.8106 | 0.9165 | 0.7317 | 0.8652 | 0.8369 | 0.8936 | 0.8872 | 0.8613 |
| LGB | ACC | 0.694 | 0.2909 | 0.6451 | 0.6505 | 0.6397 | 0.6424 | 0.6457 | 0.6209 | 0.2129 | 0.6064 | 0.5887 | 0.6241 | 0.6103 | 0.5993 |
| LGB | AC | 0.6064 | 0.2192 | 0.6048 | 0.7364 | 0.4731 | 0.5847 | 0.6496 | 0.4841 | -0.0298 | 0.4858 | 0.6383 | 0.3333 | 0.4891 | 0.5538 |
| LGB | APAAC | 0.9034 | 0.6398 | 0.8173 | 0.7689 | 0.8655 | 0.853 | 0.8068 | 0.9241 | 0.6952 | 0.8475 | 0.8369 | 0.8582 | 0.8551 | 0.8459 |
| LGB | ASDC | 0.966 | 0.8113 | 0.905 | 0.9086 | 0.9014 | 0.9041 | 0.9056 | 0.9695 | 0.8663 | 0.9326 | 0.9574 | 0.9078 | 0.9122 | 0.9343 |
| LGB | CC | 0.6701 | 0.2624 | 0.6309 | 0.6291 | 0.6327 | 0.6316 | 0.6297 | 0.6264 | 0.1986 | 0.5993 | 0.5957 | 0.6028 | 0.6 | 0.5979 |
| LGB | CKSAAGP-type-1 | 0.9174 | 0.684 | 0.8405 | 0.8584 | 0.8227 | 0.8318 | 0.8434 | 0.9661 | 0.7806 | 0.8901 | 0.9078 | 0.8723 | 0.8767 | 0.892 |
| LGB | CKSAAGP-type-2 | 0.9287 | 0.7145 | 0.8557 | 0.8566 | 0.8549 | 0.859 | 0.856 | 0.9702 | 0.8159 | 0.9078 | 0.8936 | 0.922 | 0.9197 | 0.9065 |

| | | | | | | | | | | | | | | | |
|---|---|---|---|---|---|---|---|---|---|---|---|---|---|---|---|
| LGB | CKSAAP-type-1 | 0.9581 | 0.8013 | 0.8934 | 0.8031 | 0.9838 | 0.9811 | 0.8819 | 0.9705 | 0.8544 | 0.922 | 0.844 | 1 | 1 | 0.9154 |
| LGB | CKSAAP-type-2 | 0.955 | 0.8082 | 0.8961 | 0.8012 | 0.991 | 0.9899 | 0.8842 | 0.974 | 0.8526 | 0.922 | 0.8511 | 0.9929 | 0.9917 | 0.916 |
| LGB | CTDC | 0.8802 | 0.6528 | 0.8244 | 0.7777 | 0.8711 | 0.858 | 0.8148 | 0.9034 | 0.6606 | 0.8298 | 0.8014 | 0.8582 | 0.8496 | 0.8248 |
| LGB | CTDD | 0.9775 | 0.8837 | 0.9399 | 0.9267 | 0.9535 | 0.9562 | 0.9394 | 0.9811 | 0.903 | 0.9504 | 0.9149 | 0.9858 | 0.9847 | 0.9485 |
| LGB | CTDT | 0.9443 | 0.8011 | 0.8996 | 0.8869 | 0.9123 | 0.9119 | 0.8983 | 0.9425 | 0.7661 | 0.883 | 0.8936 | 0.8723 | 0.875 | 0.8842 |
| LGB | CTriad | 0.9148 | 0.7435 | 0.8665 | 0.7903 | 0.9427 | 0.9341 | 0.8545 | 0.9402 | 0.7301 | 0.8617 | 0.7943 | 0.9291 | 0.918 | 0.8517 |
| LGB | DDE | 0.9516 | 0.7997 | 0.8898 | 0.7796 | 1 | 1 | 0.8755 | 0.9751 | 0.7938 | 0.8865 | 0.773 | 1 | 1 | 0.872 |
| LGB | DistancePair | 0.887 | 0.6428 | 0.819 | 0.7777 | 0.8601 | 0.8503 | 0.8106 | 0.9165 | 0.7317 | 0.8652 | 0.8369 | 0.8936 | 0.8872 | 0.8613 |
| LGB | DPC-type-1 | 0.9498 | 0.7968 | 0.888 | 0.7761 | 1 | 1 | 0.8731 | 0.9791 | 0.8117 | 0.8972 | 0.7943 | 1 | 1 | 0.8854 |
| LGB | DPC-type-2 | 0.9533 | 0.7993 | 0.8898 | 0.7815 | 0.9982 | 0.9979 | 0.8757 | 0.9721 | 0.8238 | 0.9043 | 0.8085 | 1 | 1 | 0.8941 |
| LGB | GDPC-type-1 | 0.9218 | 0.678 | 0.8378 | 0.8566 | 0.8189 | 0.8274 | 0.8407 | 0.9642 | 0.8085 | 0.9043 | 0.9007 | 0.9078 | 0.9071 | 0.9039 |
| LGB | GDPC-type-2 | 0.923 | 0.715 | 0.8567 | 0.871 | 0.8423 | 0.8477 | 0.8585 | 0.9775 | 0.8229 | 0.9113 | 0.922 | 0.9007 | 0.9028 | 0.9123 |
| LGB | Geary | 0.6258 | 0.2184 | 0.6084 | 0.5734 | 0.6433 | 0.618 | 0.5929 | 0.5363 | 0.0569 | 0.5284 | 0.4894 | 0.5674 | 0.5308 | 0.5092 |

| | | | | | | | | | | | | | |
|---|---|---|---|---|---|---|---|---|---|---|---|---|---|
| LGB | GTPC-type-1 | 0.8653 | 0.6052 | 0.8003 | 0.7582 | 0.8423 | 0.8303 | 0.7905 | 0.9091 | 0.698 | 0.8475 | 0.8014 | 0.8936 | 0.8828 | 0.8401 |
| LGB | GTPC-type-2 | 0.8783 | 0.6174 | 0.8074 | 0.7993 | 0.8153 | 0.8152 | 0.8054 | 0.9344 | 0.7039 | 0.8511 | 0.8156 | 0.8865 | 0.8779 | 0.8456 |
| LGB | KSCTriad | 0.9283 | 0.7267 | 0.8576 | 0.7743 | 0.9408 | 0.9291 | 0.843 | 0.9494 | 0.7236 | 0.8582 | 0.7872 | 0.9291 | 0.9174 | 0.8473 |
| LGB | Moran | 0.5748 | 0.1475 | 0.5736 | 0.5791 | 0.5681 | 0.5725 | 0.575 | 0.5382 | 0.0427 | 0.5213 | 0.4823 | 0.5603 | 0.5231 | 0.5018 |
| LGB | NMBroto | 0.6468 | 0.2396 | 0.6191 | 0.5861 | 0.6522 | 0.6272 | 0.6047 | 0.6204 | 0.192 | 0.5957 | 0.5603 | 0.6312 | 0.6031 | 0.5809 |
| LGB | PAAC | 0.8897 | 0.6406 | 0.8181 | 0.7742 | 0.862 | 0.8498 | 0.8086 | 0.9038 | 0.6353 | 0.8156 | 0.7589 | 0.8723 | 0.856 | 0.8045 |
| LGB | PDE | 0.9063 | 0.6515 | 0.8235 | 0.7777 | 0.8691 | 0.8579 | 0.8144 | 0.9275 | 0.6955 | 0.8475 | 0.8652 | 0.8298 | 0.8356 | 0.8502 |
| LGB | PseKRAAC-type-4 | 0.6023 | 0.191 | 0.5941 | 0.5807 | 0.6075 | 0.6042 | 0.5885 | 0.5523 | 0.0922 | 0.5461 | 0.5461 | 0.5461 | 0.5461 | 0.5461 |
| LGB | PseKRAAC-type-5 | 0.822 | 0.5918 | 0.7904 | 0.6991 | 0.8817 | 0.8563 | 0.7685 | 0.8398 | 0.5385 | 0.766 | 0.6879 | 0.844 | 0.8151 | 0.7462 |
| LGB | PseKRAAC-type-6A | 0.6414 | 0.2482 | 0.6236 | 0.6091 | 0.6379 | 0.6265 | 0.6161 | 0.6818 | 0.2413 | 0.6206 | 0.6383 | 0.6028 | 0.6164 | 0.6272 |
| LGB | PseKRAAC-type-13 | 0.5919 | 0.1466 | 0.5725 | 0.5503 | 0.5949 | 0.5786 | 0.562 | 0.5975 | 0.1137 | 0.5567 | 0.5887 | 0.5248 | 0.5533 | 0.5704 |
| LGB | QSOrder | 0.8911 | 0.6185 | 0.8064 | 0.7544 | 0.8583 | 0.8436 | 0.7946 | 0.9231 | 0.745 | 0.8723 | 0.8582 | 0.8865 | 0.8832 | 0.8705 |
| LGB | TPC-type-1 | 0.5045 | 0.0432 | 0.5036 | 0.4089 | 0.6018 | 0.7158 | 0.2923 | 0.5035 | 0.06 | 0.5071 | 0.0213 | 0.9929 | 0.75 | 0.0414 |

| | | | | | | | | | | | | | | | |
|---|---|---|---|---|---|---|---|---|---|---|---|---|---|---|---|
| LGB | TPC-type-2 | 0.5045 | 0.0432 | 0.5036 | 0.4089 | 0.6018 | 0.7158 | 0.2923 | 0.5035 | 0.06 | 0.5071 | 0.0213 | 0.9929 | 0.75 | 0.0414 |
| LRT | Bepler | 0.8788 | 0.6111 | 0.8046 | 0.7866 | 0.8224 | 0.817 | 0.8004 | 0.9099 | 0.6886 | 0.844 | 0.8227 | 0.8652 | 0.8593 | 0.8406 |
| LRT | CPCProt | 0.9105 | 0.6954 | 0.8459 | 0.8296 | 0.8619 | 0.8595 | 0.8421 | 0.8994 | 0.6533 | 0.8262 | 0.8014 | 0.8511 | 0.8433 | 0.8218 |
| LRT | ESM | 0.9532 | 0.7882 | 0.8933 | 0.8799 | 0.907 | 0.9065 | 0.8923 | 0.9671 | 0.809 | 0.9043 | 0.922 | 0.8865 | 0.8904 | 0.9059 |
| LRT | ESM1b | 0.9576 | 0.7994 | 0.8978 | 0.8871 | 0.9085 | 0.9095 | 0.8959 | 0.9511 | 0.8017 | 0.9007 | 0.9149 | 0.8865 | 0.8897 | 0.9021 |
| LRT | ESM1v | 0.9549 | 0.7892 | 0.8925 | 0.8747 | 0.9102 | 0.91 | 0.8894 | 0.9677 | 0.8085 | 0.9043 | 0.9078 | 0.9007 | 0.9014 | 0.9046 |
| LRT | ESM2-t6 | 0.9337 | 0.7523 | 0.8755 | 0.8745 | 0.8764 | 0.8784 | 0.8757 | 0.9728 | 0.8441 | 0.922 | 0.9149 | 0.9291 | 0.9281 | 0.9214 |
| LRT | ESM2-t12 | 0.9443 | 0.7943 | 0.8961 | 0.8927 | 0.8997 | 0.9 | 0.8951 | 0.9464 | 0.811 | 0.9043 | 0.9433 | 0.8652 | 0.875 | 0.9078 |
| LRT | ESM2-t30 | 0.9558 | 0.7741 | 0.8862 | 0.8745 | 0.8979 | 0.8967 | 0.8846 | 0.9724 | 0.8447 | 0.922 | 0.9007 | 0.9433 | 0.9407 | 0.9203 |
| LRT | ESM2-t33 | 0.9701 | 0.8392 | 0.9185 | 0.8924 | 0.9445 | 0.9416 | 0.9155 | 0.9874 | 0.9008 | 0.9504 | 0.9574 | 0.9433 | 0.9441 | 0.9507 |
| LRT | ESM2-t36 | 0.9658 | 0.8186 | 0.9086 | 0.887 | 0.9301 | 0.9273 | 0.9063 | 0.9822 | 0.88 | 0.9397 | 0.9574 | 0.922 | 0.9247 | 0.9408 |
| LRT | FastText | 0.916 | 0.7097 | 0.8531 | 0.8172 | 0.889 | 0.8812 | 0.8467 | 0.9188 | 0.7163 | 0.8582 | 0.8582 | 0.8582 | 0.8582 | 0.8582 |
| LRT | GloVe | 0.9067 | 0.6657 | 0.8316 | 0.7956 | 0.8676 | 0.858 | 0.825 | 0.902 | 0.6402 | 0.8191 | 0.7801 | 0.8582 | 0.8462 | 0.8118 |

| | | | | | | | | | | | | | | | |
|---|---|---|---|---|---|---|---|---|---|---|---|---|---|---|---|
| LRT | PRNN | 0.9415 | 0.7459 | 0.8719 | 0.8422 | 0.9015 | 0.8959 | 0.8676 | 0.9586 | 0.8087 | 0.9043 | 0.8936 | 0.9149 | 0.913 | 0.9032 |
| LRT | PTAB | 0.9137 | 0.7121 | 0.8548 | 0.8424 | 0.8674 | 0.8659 | 0.8525 | 0.9436 | 0.7519 | 0.8759 | 0.8652 | 0.8865 | 0.8841 | 0.8746 |
| LRT | PTBB | 0.9058 | 0.6661 | 0.8325 | 0.8263 | 0.8388 | 0.8369 | 0.8309 | 0.9503 | 0.7731 | 0.8865 | 0.8794 | 0.8936 | 0.8921 | 0.8857 |
| LRT | PTB | 0.9718 | 0.8341 | 0.9167 | 0.9086 | 0.9249 | 0.9238 | 0.9159 | 0.9813 | 0.8653 | 0.9326 | 0.9291 | 0.9362 | 0.9357 | 0.9324 |
| LRT | PTU | 0.9763 | 0.8401 | 0.9194 | 0.9103 | 0.9283 | 0.9276 | 0.9182 | 0.9861 | 0.8724 | 0.9362 | 0.9433 | 0.9291 | 0.9301 | 0.9366 |
| LRT | PTXU | 0.9819 | 0.8614 | 0.9301 | 0.9211 | 0.9391 | 0.9388 | 0.9293 | 0.9855 | 0.8866 | 0.9433 | 0.9362 | 0.9504 | 0.9496 | 0.9429 |
| LRT | PTXLU | 0.9219 | 0.7093 | 0.8539 | 0.8423 | 0.8656 | 0.8627 | 0.8516 | 0.9324 | 0.7033 | 0.8511 | 0.8794 | 0.8227 | 0.8322 | 0.8552 |
| LRT | Seq2Vec | 0.9838 | 0.8812 | 0.94 | 0.9247 | 0.9552 | 0.9543 | 0.9388 | 0.9869 | 0.9433 | 0.9716 | 0.9716 | 0.9716 | 0.9716 | 0.9716 |
| LRT | Word2Vec | 0.9093 | 0.6735 | 0.8352 | 0.8063 | 0.8639 | 0.857 | 0.8294 | 0.9173 | 0.7256 | 0.8617 | 0.8227 | 0.9007 | 0.8923 | 0.8561 |
| LRT | AAC | 0.8449 | 0.5687 | 0.7822 | 0.7454 | 0.819 | 0.808 | 0.7733 | 0.8554 | 0.6125 | 0.805 | 0.7589 | 0.8511 | 0.8359 | 0.7955 |
| LRT | ACC | 0.6431 | 0.2101 | 0.6047 | 0.6056 | 0.6039 | 0.6039 | 0.6039 | 0.6182 | 0.1425 | 0.5709 | 0.617 | 0.5248 | 0.5649 | 0.5898 |
| LRT | AC | 0.6119 | 0.1905 | 0.595 | 0.604 | 0.5858 | 0.594 | 0.5977 | 0.5919 | 0.1349 | 0.5674 | 0.5461 | 0.5887 | 0.5704 | 0.558 |
| LRT | APAAC | 0.8733 | 0.6036 | 0.7993 | 0.7563 | 0.8422 | 0.831 | 0.7898 | 0.8693 | 0.5754 | 0.7872 | 0.7589 | 0.8156 | 0.8045 | 0.781 |

| | | | | | | | | | | | | | | | |
|---|---|---|---|---|---|---|---|---|---|---|---|---|---|---|---|
| LRT | ASDC | 0.9329 | 0.7542 | 0.8754 | 0.8458 | 0.905 | 0.9006 | 0.871 | 0.9631 | 0.8305 | 0.9149 | 0.8936 | 0.9362 | 0.9333 | 0.913 |
| LRT | CC | 0.6467 | 0.2231 | 0.6111 | 0.6219 | 0.6003 | 0.6095 | 0.6143 | 0.6253 | 0.1988 | 0.5993 | 0.6241 | 0.5745 | 0.5946 | 0.609 |
| LRT | CKSAAGP-type-1 | 0.962 | 0.8351 | 0.9167 | 0.9032 | 0.9302 | 0.9294 | 0.9153 | 0.9808 | 0.8727 | 0.9362 | 0.922 | 0.9504 | 0.9489 | 0.9353 |
| LRT | CKSAAGP-type-2 | 0.9709 | 0.8456 | 0.9221 | 0.9158 | 0.9284 | 0.9285 | 0.9213 | 0.9905 | 0.8869 | 0.9433 | 0.9291 | 0.9574 | 0.9562 | 0.9424 |
| LRT | CKSAAP-type-1 | 0.9438 | 0.7666 | 0.8817 | 0.8477 | 0.9159 | 0.9104 | 0.877 | 0.9362 | 0.7466 | 0.8723 | 0.8369 | 0.9078 | 0.9008 | 0.8676 |
| LRT | CKSAAP-type-2 | 0.9442 | 0.7723 | 0.8844 | 0.853 | 0.9159 | 0.9124 | 0.8805 | 0.9508 | 0.7817 | 0.8901 | 0.8582 | 0.922 | 0.9167 | 0.8864 |
| LRT | CTDC | 0.8444 | 0.5719 | 0.784 | 0.7472 | 0.8207 | 0.81 | 0.7756 | 0.8532 | 0.6118 | 0.805 | 0.766 | 0.844 | 0.8308 | 0.797 |
| LRT | CTDD | 0.973 | 0.8684 | 0.9337 | 0.9355 | 0.9319 | 0.9329 | 0.9337 | 0.9686 | 0.8795 | 0.9397 | 0.9433 | 0.9362 | 0.9366 | 0.9399 |
| LRT | CTDT | 0.8796 | 0.6639 | 0.8306 | 0.7902 | 0.871 | 0.8599 | 0.8232 | 0.9092 | 0.6859 | 0.8404 | 0.7801 | 0.9007 | 0.8871 | 0.8302 |
| LRT | CTriad | 0.9497 | 0.7891 | 0.8933 | 0.8709 | 0.9157 | 0.9131 | 0.8904 | 0.9683 | 0.8298 | 0.9149 | 0.9149 | 0.9149 | 0.9149 | 0.9149 |
| LRT | DDE | 0.9413 | 0.7527 | 0.8755 | 0.8656 | 0.8853 | 0.8846 | 0.874 | 0.9476 | 0.7454 | 0.8723 | 0.8511 | 0.8936 | 0.8889 | 0.8696 |
| LRT | DistancePair | 0.8449 | 0.5687 | 0.7822 | 0.7454 | 0.819 | 0.808 | 0.7733 | 0.8554 | 0.6125 | 0.805 | 0.7589 | 0.8511 | 0.8359 | 0.7955 |
| LRT | DPC-type-1 | 0.9475 | 0.7997 | 0.8988 | 0.8819 | 0.9159 | 0.9139 | 0.8965 | 0.9472 | 0.7317 | 0.8652 | 0.8369 | 0.8936 | 0.8872 | 0.8613 |

| | | | | | | | | | | | | | | | |
|---|---|---|---|---|---|---|---|---|---|---|---|---|---|---|---|
| LRT | DPC-type-2 | 0.9465 | 0.7903 | 0.8934 | 0.8639 | 0.923 | 0.92 | 0.8896 | 0.9481 | 0.7522 | 0.8759 | 0.8582 | 0.8936 | 0.8897 | 0.8736 |
| LRT | GDPC-type-1 | 0.9717 | 0.8578 | 0.9283 | 0.9159 | 0.9409 | 0.9399 | 0.9272 | 0.9915 | 0.922 | 0.961 | 0.9645 | 0.9574 | 0.9577 | 0.9611 |
| LRT | GDPC-type-2 | 0.9691 | 0.8225 | 0.9096 | 0.8746 | 0.9446 | 0.9409 | 0.9057 | 0.9919 | 0.8805 | 0.9397 | 0.9149 | 0.9645 | 0.9627 | 0.9382 |
| LRT | Geary | 0.6404 | 0.2289 | 0.6137 | 0.6218 | 0.6057 | 0.6111 | 0.6153 | 0.5933 | 0.0923 | 0.5461 | 0.5177 | 0.5745 | 0.5489 | 0.5328 |
| LRT | GTPC-type-1 | 0.9564 | 0.8106 | 0.9042 | 0.8997 | 0.9086 | 0.9084 | 0.9027 | 0.9819 | 0.8679 | 0.9326 | 0.8936 | 0.9716 | 0.9692 | 0.9299 |
| LRT | GTPC-type-2 | 0.9661 | 0.8486 | 0.9239 | 0.9212 | 0.9266 | 0.9261 | 0.9233 | 0.9913 | 0.9231 | 0.961 | 0.9362 | 0.9858 | 0.9851 | 0.96 |
| LRT | KSCTriad | 0.9346 | 0.7503 | 0.8719 | 0.8172 | 0.9266 | 0.9188 | 0.8633 | 0.9361 | 0.7189 | 0.8582 | 0.8156 | 0.9007 | 0.8915 | 0.8519 |
| LRT | Moran | 0.6065 | 0.1661 | 0.5825 | 0.5913 | 0.5734 | 0.5835 | 0.5852 | 0.5609 | 0.1277 | 0.5638 | 0.5745 | 0.5532 | 0.5625 | 0.5684 |
| LRT | NMBroto | 0.5952 | 0.1598 | 0.5797 | 0.5627 | 0.5965 | 0.5827 | 0.5716 | 0.5963 | 0.1918 | 0.5957 | 0.5674 | 0.6241 | 0.6015 | 0.5839 |
| LRT | PAAC | 0.874 | 0.6257 | 0.8109 | 0.7706 | 0.8513 | 0.8387 | 0.8016 | 0.8634 | 0.6254 | 0.8121 | 0.7801 | 0.844 | 0.8333 | 0.8059 |
| LRT | PDE | 0.8743 | 0.618 | 0.8073 | 0.7723 | 0.8421 | 0.8326 | 0.7999 | 0.8788 | 0.6044 | 0.8014 | 0.766 | 0.8369 | 0.8244 | 0.7941 |
| LRT | PseKRAAC-type-4 | 0.591 | 0.1128 | 0.5556 | 0.5288 | 0.5827 | 0.5623 | 0.5422 | 0.5885 | 0.1207 | 0.5603 | 0.5816 | 0.539 | 0.5578 | 0.5694 |
| LRT | PseKRAAC-type-5 | 0.8437 | 0.5799 | 0.7814 | 0.6704 | 0.8925 | 0.8652 | 0.7524 | 0.8491 | 0.5412 | 0.766 | 0.6738 | 0.8582 | 0.8261 | 0.7422 |

| | | | | | | | | | | | | | |
|---|---|---|---|---|---|---|---|---|---|---|---|---|---|
| LRT | PseKRAAC-type-6A | 0.6373 | 0.2146 | 0.6066 | 0.5789 | 0.6343 | 0.6127 | 0.593 | 0.6317 | 0.0784 | 0.539 | 0.4894 | 0.5887 | 0.5433 | 0.5149 |
| LRT | PseKRAAC-type-13 | 0.5686 | 0.1195 | 0.5591 | 0.5539 | 0.5644 | 0.5589 | 0.553 | 0.5939 | 0.1277 | 0.5638 | 0.5461 | 0.5816 | 0.5662 | 0.556 |
| LRT | QSOrder | 0.8807 | 0.6347 | 0.8164 | 0.7994 | 0.8335 | 0.8284 | 0.8125 | 0.9033 | 0.6667 | 0.8333 | 0.8369 | 0.8298 | 0.831 | 0.8339 |
| LRT | TPC-type-1 | 0.8708 | 0.5805 | 0.7876 | 0.7312 | 0.844 | 0.8255 | 0.7741 | 0.9071 | 0.6249 | 0.8121 | 0.7872 | 0.8369 | 0.8284 | 0.8073 |
| LRT | TPC-type-2 | 0.8661 | 0.5633 | 0.7787 | 0.7095 | 0.8476 | 0.8229 | 0.7612 | 0.9096 | 0.6716 | 0.8333 | 0.773 | 0.8936 | 0.879 | 0.8226 |
| RF | Bepler | 0.8652 | 0.5781 | 0.7876 | 0.769 | 0.8064 | 0.7999 | 0.7821 | 0.8911 | 0.6456 | 0.8227 | 0.8085 | 0.8369 | 0.8321 | 0.8201 |
| RF | CPCProt | 0.8958 | 0.6562 | 0.8262 | 0.7832 | 0.8692 | 0.8575 | 0.8176 | 0.8937 | 0.6464 | 0.8227 | 0.7943 | 0.8511 | 0.8421 | 0.8175 |
| RF | ESM | 0.9154 | 0.6911 | 0.8431 | 0.7975 | 0.8887 | 0.8807 | 0.8356 | 0.9245 | 0.6859 | 0.8404 | 0.7801 | 0.9007 | 0.8871 | 0.8302 |
| RF | ESM1b | 0.8992 | 0.6817 | 0.8387 | 0.7939 | 0.8835 | 0.8732 | 0.8304 | 0.9201 | 0.6829 | 0.8404 | 0.8014 | 0.8794 | 0.8692 | 0.8339 |
| RF | ESM1v | 0.8895 | 0.6585 | 0.8271 | 0.785 | 0.8692 | 0.8587 | 0.8185 | 0.9052 | 0.6442 | 0.8191 | 0.7518 | 0.8865 | 0.8689 | 0.8061 |
| RF | ESM2-t6 | 0.8861 | 0.6364 | 0.8164 | 0.7886 | 0.8441 | 0.837 | 0.8101 | 0.9059 | 0.647 | 0.8227 | 0.7872 | 0.8582 | 0.8473 | 0.8162 |
| RF | ESM2-t12 | 0.9029 | 0.6591 | 0.8279 | 0.8047 | 0.8512 | 0.8459 | 0.8231 | 0.9237 | 0.6695 | 0.8333 | 0.7872 | 0.8794 | 0.8672 | 0.8253 |
| RF | ESM2-t30 | 0.897 | 0.6625 | 0.8289 | 0.7922 | 0.8656 | 0.858 | 0.8214 | 0.9083 | 0.6639 | 0.8298 | 0.773 | 0.8865 | 0.872 | 0.8195 |

| | | | | | | | | | | | | | | | |
|---|---|---|---|---|---|---|---|---|---|---|---|---|---|---|---|
| RF | ESM2-t33 | 0.9101 | 0.7058 | 0.8504 | 0.8011 | 0.8996 | 0.8891 | 0.8413 | 0.9156 | 0.6533 | 0.8262 | 0.8014 | 0.8511 | 0.8433 | 0.8218 |
| RF | ESM2-t36 | 0.914 | 0.7041 | 0.8486 | 0.792 | 0.905 | 0.8952 | 0.8384 | 0.9256 | 0.7236 | 0.8582 | 0.7872 | 0.9291 | 0.9174 | 0.8473 |
| RF | FastText | 0.8846 | 0.6401 | 0.8181 | 0.7832 | 0.8531 | 0.8445 | 0.8111 | 0.8972 | 0.5892 | 0.7943 | 0.773 | 0.8156 | 0.8074 | 0.7899 |
| RF | GloVe | 0.8844 | 0.6124 | 0.8055 | 0.7957 | 0.8153 | 0.813 | 0.8035 | 0.8911 | 0.6596 | 0.8298 | 0.8227 | 0.8369 | 0.8345 | 0.8286 |
| RF | PRNN | 0.868 | 0.5904 | 0.7929 | 0.7723 | 0.8136 | 0.8065 | 0.7855 | 0.8874 | 0.5759 | 0.7872 | 0.7518 | 0.8227 | 0.8092 | 0.7794 |
| RF | PTAB | 0.8777 | 0.6306 | 0.8145 | 0.7993 | 0.8298 | 0.8256 | 0.8113 | 0.9077 | 0.6176 | 0.8085 | 0.7872 | 0.8298 | 0.8222 | 0.8043 |
| RF | PTBB | 0.8608 | 0.5876 | 0.793 | 0.7669 | 0.819 | 0.8092 | 0.7868 | 0.8986 | 0.6118 | 0.805 | 0.766 | 0.844 | 0.8308 | 0.797 |
| RF | PTB | 0.9143 | 0.6842 | 0.8396 | 0.7994 | 0.8799 | 0.8713 | 0.8317 | 0.9282 | 0.698 | 0.8475 | 0.8014 | 0.8936 | 0.8828 | 0.8401 |
| RF | PTU | 0.9338 | 0.7211 | 0.8584 | 0.8244 | 0.8925 | 0.8875 | 0.8531 | 0.935 | 0.7236 | 0.8582 | 0.7872 | 0.9291 | 0.9174 | 0.8473 |
| RF | PTXU | 0.9313 | 0.7286 | 0.862 | 0.8209 | 0.9032 | 0.8966 | 0.8556 | 0.9285 | 0.6663 | 0.8298 | 0.7589 | 0.9007 | 0.8843 | 0.8168 |
| RF | PTXLU | 0.7939 | 0.4577 | 0.7276 | 0.7457 | 0.7096 | 0.7214 | 0.7318 | 0.7889 | 0.4114 | 0.7057 | 0.695 | 0.7163 | 0.7101 | 0.7025 |
| RF | Seq2Vec | 0.9505 | 0.7862 | 0.8916 | 0.8603 | 0.923 | 0.9192 | 0.8877 | 0.9652 | 0.8021 | 0.9007 | 0.8794 | 0.922 | 0.9185 | 0.8986 |
| RF | Word2Vec | 0.8632 | 0.5914 | 0.7948 | 0.7724 | 0.8171 | 0.8101 | 0.7899 | 0.8864 | 0.6322 | 0.8156 | 0.7872 | 0.844 | 0.8346 | 0.8102 |

| | | | | | | | | | | | | | | | |
|---|---|---|---|---|---|---|---|---|---|---|---|---|---|---|---|
| RF | AAC | 0.8815 | 0.6073 | 0.8011 | 0.7437 | 0.8583 | 0.8418 | 0.7889 | 0.9184 | 0.6822 | 0.8404 | 0.8085 | 0.8723 | 0.8636 | 0.8352 |
| RF | ACC | 0.6588 | 0.2675 | 0.6335 | 0.6201 | 0.6469 | 0.6361 | 0.6274 | 0.6343 | 0.2272 | 0.6135 | 0.5887 | 0.6383 | 0.6194 | 0.6036 |
| RF | AC | 0.6453 | 0.2537 | 0.6264 | 0.6345 | 0.6184 | 0.6258 | 0.6292 | 0.6312 | 0.1934 | 0.5957 | 0.5248 | 0.6667 | 0.6116 | 0.5649 |
| RF | APAAC | 0.891 | 0.627 | 0.8119 | 0.7833 | 0.8403 | 0.833 | 0.8057 | 0.906 | 0.6882 | 0.844 | 0.8298 | 0.8582 | 0.854 | 0.8417 |
| RF | ASDC | 0.9492 | 0.7911 | 0.8943 | 0.8692 | 0.9193 | 0.9165 | 0.8912 | 0.959 | 0.7803 | 0.8901 | 0.9007 | 0.8794 | 0.8819 | 0.8912 |
| RF | CC | 0.6548 | 0.2597 | 0.6291 | 0.5989 | 0.6596 | 0.637 | 0.6162 | 0.638 | 0.2067 | 0.6028 | 0.5532 | 0.6525 | 0.6142 | 0.5821 |
| RF | CKSAAGP-type-1 | 0.9024 | 0.6738 | 0.8352 | 0.8442 | 0.8261 | 0.8332 | 0.8366 | 0.9466 | 0.7234 | 0.8617 | 0.8652 | 0.8582 | 0.8592 | 0.8622 |
| RF | CKSAAGP-type-2 | 0.9065 | 0.67 | 0.8334 | 0.8317 | 0.8351 | 0.8377 | 0.8327 | 0.95 | 0.7811 | 0.8901 | 0.8652 | 0.9149 | 0.9104 | 0.8873 |
| RF | CKSAAP-type-1 | 0.9563 | 0.8174 | 0.9059 | 0.8567 | 0.9552 | 0.9514 | 0.9005 | 0.9636 | 0.8034 | 0.9007 | 0.8652 | 0.9362 | 0.9313 | 0.8971 |
| RF | CKSAAP-type-2 | 0.9553 | 0.8324 | 0.9132 | 0.8604 | 0.966 | 0.9625 | 0.9074 | 0.9609 | 0.8252 | 0.9113 | 0.8723 | 0.9504 | 0.9462 | 0.9077 |
| RF | CTDC | 0.8707 | 0.6005 | 0.7975 | 0.74 | 0.8548 | 0.8381 | 0.7848 | 0.8916 | 0.626 | 0.8121 | 0.773 | 0.8511 | 0.8385 | 0.8044 |
| RF | CTDD | 0.9857 | 0.8995 | 0.9489 | 0.9248 | 0.9732 | 0.9722 | 0.9475 | 0.9885 | 0.9299 | 0.9645 | 0.9433 | 0.9858 | 0.9852 | 0.9638 |
| RF | CTDT | 0.9173 | 0.7393 | 0.8665 | 0.8119 | 0.9212 | 0.9135 | 0.8582 | 0.9153 | 0.7181 | 0.8582 | 0.8227 | 0.8936 | 0.8855 | 0.8529 |

| | | | | | | | | | | | | | | | |
|---|---|---|---|---|---|---|---|---|---|---|---|---|---|---|---|
| RF | CTriad | 0.9385 | 0.8007 | 0.8952 | 0.8226 | 0.9678 | 0.9631 | 0.8856 | 0.9556 | 0.7764 | 0.883 | 0.8014 | 0.9645 | 0.9576 | 0.8726 |
| RF | DDE | 0.9351 | 0.7743 | 0.8836 | 0.83 | 0.9373 | 0.9307 | 0.8755 | 0.9457 | 0.7701 | 0.8794 | 0.7943 | 0.9645 | 0.9573 | 0.8682 |
| RF | DistancePair | 0.8815 | 0.6073 | 0.8011 | 0.7437 | 0.8583 | 0.8418 | 0.7889 | 0.9184 | 0.6822 | 0.8404 | 0.8085 | 0.8723 | 0.8636 | 0.8352 |
| RF | DPC-type-1 | 0.9593 | 0.828 | 0.9105 | 0.8514 | 0.9695 | 0.9657 | 0.9039 | 0.9742 | 0.8559 | 0.9255 | 0.8723 | 0.9787 | 0.9762 | 0.9213 |
| RF | DPC-type-2 | 0.9628 | 0.8438 | 0.9185 | 0.8604 | 0.9767 | 0.9737 | 0.9127 | 0.9719 | 0.8755 | 0.9362 | 0.8936 | 0.9787 | 0.9767 | 0.9333 |
| RF | GDPC-type-1 | 0.8909 | 0.6559 | 0.8262 | 0.8244 | 0.8278 | 0.8314 | 0.8255 | 0.9392 | 0.7243 | 0.8617 | 0.8369 | 0.8865 | 0.8806 | 0.8582 |
| RF | GDPC-type-2 | 0.8983 | 0.6511 | 0.8244 | 0.8226 | 0.8261 | 0.8272 | 0.8234 | 0.9467 | 0.7885 | 0.8936 | 0.8652 | 0.922 | 0.9173 | 0.8905 |
| RF | Geary | 0.6396 | 0.2183 | 0.6084 | 0.5824 | 0.6344 | 0.6148 | 0.5962 | 0.5715 | 0.1349 | 0.5674 | 0.5461 | 0.5887 | 0.5704 | 0.558 |
| RF | GTPC-type-1 | 0.8794 | 0.6393 | 0.8182 | 0.8067 | 0.8297 | 0.8288 | 0.8157 | 0.9247 | 0.717 | 0.8582 | 0.8369 | 0.8794 | 0.8741 | 0.8551 |
| RF | GTPC-type-2 | 0.8799 | 0.6397 | 0.8182 | 0.7975 | 0.8387 | 0.8341 | 0.8134 | 0.9311 | 0.717 | 0.8582 | 0.8369 | 0.8794 | 0.8741 | 0.8551 |
| RF | KSCTriad | 0.9375 | 0.7743 | 0.8835 | 0.8208 | 0.9463 | 0.9393 | 0.8752 | 0.9461 | 0.7287 | 0.8617 | 0.8014 | 0.922 | 0.9113 | 0.8528 |
| RF | Moran | 0.5917 | 0.1958 | 0.5977 | 0.5931 | 0.6023 | 0.5983 | 0.5949 | 0.5266 | 0.0645 | 0.5319 | 0.461 | 0.6028 | 0.5372 | 0.4962 |
| RF | NMBroto | 0.6592 | 0.2559 | 0.6272 | 0.6272 | 0.6271 | 0.6278 | 0.6254 | 0.6045 | 0.1493 | 0.5745 | 0.539 | 0.6099 | 0.5802 | 0.5588 |

| | | | | | | | | | | | | | | | |
|---|---|---|---|---|---|---|---|---|---|---|---|---|---|---|---|
| RF | PAAC | 0.8891 | 0.626 | 0.8109 | 0.7651 | 0.8565 | 0.8433 | 0.8009 | 0.897 | 0.641 | 0.8191 | 0.773 | 0.8652 | 0.8516 | 0.8104 |
| RF | PDE | 0.893 | 0.6291 | 0.8118 | 0.7633 | 0.86 | 0.847 | 0.801 | 0.8996 | 0.6243 | 0.8121 | 0.8014 | 0.8227 | 0.8188 | 0.81 |
| RF | PseKRAAC-type-4 | 0.6209 | 0.2188 | 0.6084 | 0.588 | 0.6291 | 0.614 | 0.5984 | 0.5406 | 0.0572 | 0.5284 | 0.4681 | 0.5887 | 0.5323 | 0.4981 |
| RF | PseKRAAC-type-5 | 0.8173 | 0.5117 | 0.7527 | 0.6972 | 0.8082 | 0.7884 | 0.7373 | 0.8217 | 0.4944 | 0.7447 | 0.6738 | 0.8156 | 0.7851 | 0.7252 |
| RF | PseKRAAC-type-6A | 0.6586 | 0.2445 | 0.6218 | 0.6201 | 0.6237 | 0.6224 | 0.6203 | 0.6848 | 0.2412 | 0.6206 | 0.6099 | 0.6312 | 0.6232 | 0.6165 |
| RF | PseKRAAC-type-13 | 0.5616 | 0.1057 | 0.5528 | 0.5628 | 0.5428 | 0.5545 | 0.5579 | 0.6172 | 0.1632 | 0.5816 | 0.5957 | 0.5674 | 0.5793 | 0.5874 |
| RF | QSOrder | 0.8889 | 0.6277 | 0.8119 | 0.7708 | 0.853 | 0.8414 | 0.8031 | 0.9121 | 0.7033 | 0.8511 | 0.8227 | 0.8794 | 0.8722 | 0.8467 |
| RF | TPC-type-1 | 0.8666 | 0.5854 | 0.7867 | 0.6952 | 0.8781 | 0.8537 | 0.7643 | 0.9211 | 0.695 | 0.844 | 0.773 | 0.9149 | 0.9008 | 0.8321 |
| RF | TPC-type-2 | 0.8671 | 0.5886 | 0.7876 | 0.6881 | 0.887 | 0.86 | 0.7622 | 0.9155 | 0.6895 | 0.8369 | 0.7305 | 0.9433 | 0.9279 | 0.8175 |
| SVM | Bepler | 0.8943 | 0.6604 | 0.8288 | 0.8064 | 0.8512 | 0.8457 | 0.8241 | 0.9201 | 0.6891 | 0.844 | 0.8156 | 0.8723 | 0.8647 | 0.8394 |
| SVM | CPCProt | 0.9075 | 0.7031 | 0.8477 | 0.7814 | 0.914 | 0.9014 | 0.8357 | 0.906 | 0.6885 | 0.8404 | 0.766 | 0.9149 | 0.9 | 0.8276 |
| SVM | ESM | 0.9541 | 0.8105 | 0.9041 | 0.8781 | 0.9302 | 0.9279 | 0.9016 | 0.9741 | 0.8017 | 0.9007 | 0.9149 | 0.8865 | 0.8897 | 0.9021 |
| SVM | ESM1b | 0.9537 | 0.7755 | 0.8853 | 0.8781 | 0.8924 | 0.8957 | 0.8837 | 0.9497 | 0.7943 | 0.8972 | 0.8936 | 0.9007 | 0.9 | 0.8968 |

| | | | | | | | | | | | | | | | |
|---|---|---|---|---|---|---|---|---|---|---|---|---|---|---|---|
| SVM | ESM1v | 0.9584 | 0.7772 | 0.8871 | 0.88 | 0.8941 | 0.8943 | 0.8851 | 0.9716 | 0.8232 | 0.9113 | 0.9291 | 0.8936 | 0.8973 | 0.9129 |
| SVM | ESM2-t6 | 0.9339 | 0.7591 | 0.879 | 0.8798 | 0.8781 | 0.8796 | 0.8791 | 0.9734 | 0.8443 | 0.922 | 0.9362 | 0.9078 | 0.9103 | 0.9231 |
| SVM | ESM2-t12 | 0.9501 | 0.8101 | 0.9042 | 0.8944 | 0.914 | 0.9124 | 0.9023 | 0.9504 | 0.8043 | 0.9007 | 0.9433 | 0.8582 | 0.8693 | 0.9048 |
| SVM | ESM2-t30 | 0.9537 | 0.7837 | 0.8907 | 0.88 | 0.9014 | 0.9012 | 0.8891 | 0.9676 | 0.8441 | 0.922 | 0.9149 | 0.9291 | 0.9281 | 0.9214 |
| SVM | ESM2-t33 | 0.9526 | 0.7994 | 0.8979 | 0.8672 | 0.9283 | 0.9245 | 0.8935 | 0.9729 | 0.8305 | 0.9149 | 0.8936 | 0.9362 | 0.9333 | 0.913 |
| SVM | ESM2-t36 | 0.5 | 0.326 | 0.6607 | 0.6643 | 0.6607 | 0.561 | 0.5949 | 0.4965 | 0.6028 | 0.8014 | 0.8014 | 0.8014 | 0.8014 | 0.8014 |
| SVM | FastText | 0.9303 | 0.7593 | 0.8773 | 0.8316 | 0.923 | 0.9152 | 0.8702 | 0.9371 | 0.7381 | 0.8688 | 0.8511 | 0.8865 | 0.8824 | 0.8664 |
| SVM | GloVe | 0.9124 | 0.6973 | 0.8477 | 0.828 | 0.8676 | 0.8632 | 0.8443 | 0.911 | 0.668 | 0.8333 | 0.8014 | 0.8652 | 0.8561 | 0.8278 |
| SVM | PRNN | 0.9461 | 0.7594 | 0.879 | 0.8656 | 0.8924 | 0.8899 | 0.8767 | 0.9621 | 0.7589 | 0.8794 | 0.8794 | 0.8794 | 0.8794 | 0.8794 |
| SVM | PTAB | 0.8472 | 0.5443 | 0.767 | 0.7728 | 0.762 | 0.7766 | 0.7676 | 0.8671 | 0.5122 | 0.7553 | 0.7943 | 0.7163 | 0.7368 | 0.7645 |
| SVM | PTBB | 0.9043 | 0.6655 | 0.8325 | 0.8281 | 0.837 | 0.835 | 0.8313 | 0.9467 | 0.7589 | 0.8794 | 0.8794 | 0.8794 | 0.8794 | 0.8794 |
| SVM | PTB | 0.9675 | 0.8238 | 0.9113 | 0.8979 | 0.9249 | 0.9236 | 0.91 | 0.9837 | 0.8658 | 0.9326 | 0.9504 | 0.9149 | 0.9178 | 0.9338 |
| SVM | PTU | 0.9749 | 0.8364 | 0.9176 | 0.9104 | 0.9248 | 0.9241 | 0.9165 | 0.988 | 0.8938 | 0.9468 | 0.9574 | 0.9362 | 0.9375 | 0.9474 |

| | | | | | | | | | | | | | | | |
|---|---|---|---|---|---|---|---|---|---|---|---|---|---|---|---|
| SVM | PTXU | 0.9822 | 0.8618 | 0.9301 | 0.9158 | 0.9445 | 0.9437 | 0.9288 | 0.99 | 0.8938 | 0.9468 | 0.9362 | 0.9574 | 0.9565 | 0.9462 |
| SVM | PTXLU | 0.9255 | 0.7186 | 0.8584 | 0.8422 | 0.8745 | 0.8713 | 0.8557 | 0.9242 | 0.6822 | 0.8404 | 0.8723 | 0.8085 | 0.82 | 0.8454 |
| SVM | Seq2Vec | 0.982 | 0.8734 | 0.9355 | 0.9086 | 0.9624 | 0.9607 | 0.9332 | 0.9962 | 0.9291 | 0.9645 | 0.9645 | 0.9645 | 0.9645 | 0.9645 |
| SVM | Word2Vec | 0.918 | 0.6936 | 0.8459 | 0.8207 | 0.8709 | 0.8642 | 0.8412 | 0.9436 | 0.7608 | 0.8794 | 0.844 | 0.9149 | 0.9084 | 0.875 |
| SVM | AAC | 0.8452 | 0.5876 | 0.7913 | 0.7401 | 0.8423 | 0.8277 | 0.78 | 0.875 | 0.6598 | 0.8262 | 0.7518 | 0.9007 | 0.8833 | 0.8123 |
| SVM | ACC | 0.6827 | 0.2767 | 0.638 | 0.6453 | 0.6309 | 0.6363 | 0.6402 | 0.6492 | 0.2483 | 0.6241 | 0.617 | 0.6312 | 0.6259 | 0.6214 |
| SVM | AC | 0.6242 | 0.221 | 0.6102 | 0.6291 | 0.5911 | 0.6085 | 0.6175 | 0.59 | 0.1421 | 0.5709 | 0.6028 | 0.539 | 0.5667 | 0.5842 |
| SVM | APAAC | 0.8711 | 0.6154 | 0.8047 | 0.76 | 0.8494 | 0.8378 | 0.7943 | 0.86 | 0.5901 | 0.7943 | 0.7589 | 0.8298 | 0.8168 | 0.7868 |
| SVM | ASDC | 0.9335 | 0.7685 | 0.8817 | 0.8315 | 0.932 | 0.9245 | 0.8746 | 0.9609 | 0.8244 | 0.9113 | 0.8794 | 0.9433 | 0.9394 | 0.9084 |
| SVM | CC | 0.6711 | 0.2768 | 0.638 | 0.6613 | 0.6148 | 0.6324 | 0.6459 | 0.6383 | 0.1773 | 0.5887 | 0.5887 | 0.5887 | 0.5887 | 0.5887 |
| SVM | CKSAAGP-type-1 | 0.9693 | 0.8702 | 0.9346 | 0.9247 | 0.9445 | 0.944 | 0.9338 | 0.9892 | 0.915 | 0.9574 | 0.9645 | 0.9504 | 0.951 | 0.9577 |
| SVM | CKSAAGP-type-2 | 0.9725 | 0.8794 | 0.9391 | 0.9302 | 0.9481 | 0.948 | 0.9384 | 0.9891 | 0.9149 | 0.9574 | 0.9574 | 0.9574 | 0.9574 | 0.9574 |
| SVM | CKSAAP-type-1 | 0.8437 | 0.4696 | 0.723 | 0.6244 | 0.823 | 0.7947 | 0.6835 | 0.8442 | 0.4704 | 0.727 | 0.5957 | 0.8582 | 0.8077 | 0.6857 |

| | | | | | | | | | | | | | | | |
|---|---|---|---|---|---|---|---|---|---|---|---|---|---|---|---|
| SVM | CKSAAP-type-2 | 0.8696 | 0.5476 | 0.7492 | 0.5468 | 0.9516 | 0.9234 | 0.682 | 0.8496 | 0.5524 | 0.766 | 0.6312 | 0.9007 | 0.8641 | 0.7295 |
| SVM | CTDC | 0.8465 | 0.5902 | 0.7921 | 0.7382 | 0.8459 | 0.8294 | 0.7788 | 0.8584 | 0.6068 | 0.8014 | 0.7447 | 0.8582 | 0.84 | 0.7895 |
| SVM | CTDD | 0.9724 | 0.8751 | 0.9373 | 0.9373 | 0.9373 | 0.9382 | 0.9375 | 0.9739 | 0.8938 | 0.9468 | 0.9362 | 0.9574 | 0.9565 | 0.9462 |
| SVM | CTDT | 0.8848 | 0.7143 | 0.8539 | 0.792 | 0.9158 | 0.9045 | 0.8437 | 0.9033 | 0.7365 | 0.8652 | 0.8014 | 0.9291 | 0.9187 | 0.8561 |
| SVM | CTriad | 0.954 | 0.8114 | 0.9041 | 0.8763 | 0.9319 | 0.9292 | 0.9007 | 0.9589 | 0.8301 | 0.9149 | 0.9291 | 0.9007 | 0.9034 | 0.9161 |
| SVM | DDE | 0.9444 | 0.778 | 0.888 | 0.8763 | 0.8997 | 0.8992 | 0.8865 | 0.9541 | 0.7519 | 0.8759 | 0.8652 | 0.8865 | 0.8841 | 0.8746 |
| SVM | DistancePair | 0.8452 | 0.5876 | 0.7913 | 0.7401 | 0.8423 | 0.8277 | 0.78 | 0.875 | 0.6598 | 0.8262 | 0.7518 | 0.9007 | 0.8833 | 0.8123 |
| SVM | DPC-type-1 | 0.9549 | 0.8142 | 0.9059 | 0.88 | 0.9319 | 0.9291 | 0.9031 | 0.9489 | 0.7341 | 0.8652 | 0.8156 | 0.9149 | 0.9055 | 0.8582 |
| SVM | DPC-type-2 | 0.9433 | 0.7842 | 0.8898 | 0.8459 | 0.9337 | 0.9288 | 0.8843 | 0.9568 | 0.7885 | 0.8936 | 0.8652 | 0.922 | 0.9173 | 0.8905 |
| SVM | GDPC-type-1 | 0.9654 | 0.8792 | 0.9391 | 0.9428 | 0.9356 | 0.9365 | 0.9391 | 0.9867 | 0.9368 | 0.9681 | 0.9858 | 0.9504 | 0.9521 | 0.9686 |
| SVM | GDPC-type-2 | 0.9719 | 0.8793 | 0.9391 | 0.9319 | 0.9463 | 0.9459 | 0.9384 | 0.9917 | 0.9292 | 0.9645 | 0.9716 | 0.9574 | 0.958 | 0.9648 |
| SVM | Geary | 0.6434 | 0.2442 | 0.6218 | 0.6255 | 0.6183 | 0.6216 | 0.623 | 0.5891 | 0.0997 | 0.5496 | 0.5035 | 0.5957 | 0.5547 | 0.5279 |
| SVM | GTPC-type-1 | 0.9595 | 0.837 | 0.9176 | 0.9087 | 0.9265 | 0.9265 | 0.9165 | 0.9862 | 0.9011 | 0.9504 | 0.9362 | 0.9645 | 0.9635 | 0.9496 |

| | | | | | | | | | | | | | | | |
|---|---|---|---|---|---|---|---|---|---|---|---|---|---|---|---|
| SVM | GTPC-type-2 | 0.9684 | 0.877 | 0.9382 | 0.9409 | 0.9356 | 0.9363 | 0.9383 | 0.99 | 0.9506 | 0.9752 | 0.9645 | 0.9858 | 0.9855 | 0.9749 |
| SVM | KSCTriad | 0.9304 | 0.7421 | 0.8665 | 0.792 | 0.9409 | 0.9305 | 0.8545 | 0.923 | 0.7172 | 0.8546 | 0.7801 | 0.9291 | 0.9167 | 0.8429 |
| SVM | Moran | 0.6108 | 0.1978 | 0.5985 | 0.6165 | 0.5805 | 0.5962 | 0.6048 | 0.5541 | 0.0993 | 0.5496 | 0.5461 | 0.5532 | 0.55 | 0.548 |
| SVM | NMBroto | 0.6344 | 0.2247 | 0.6119 | 0.6253 | 0.5984 | 0.6093 | 0.616 | 0.6091 | 0.1773 | 0.5887 | 0.5887 | 0.5887 | 0.5887 | 0.5887 |
| SVM | PAAC | 0.8855 | 0.6474 | 0.8208 | 0.7653 | 0.8764 | 0.8625 | 0.8095 | 0.9003 | 0.6755 | 0.8369 | 0.8014 | 0.8723 | 0.8626 | 0.8309 |
| SVM | PDE | 0.8951 | 0.6526 | 0.8235 | 0.7724 | 0.8745 | 0.862 | 0.8131 | 0.9105 | 0.6695 | 0.8333 | 0.7872 | 0.8794 | 0.8672 | 0.8253 |
| SVM | PseKRAAC-type-4 | 0.6258 | 0.2196 | 0.6084 | 0.5611 | 0.6561 | 0.622 | 0.5876 | 0.6061 | 0.1776 | 0.5887 | 0.5603 | 0.617 | 0.594 | 0.5766 |
| SVM | PseKRAAC-type-5 | 0.8321 | 0.5951 | 0.7877 | 0.6668 | 0.9087 | 0.8823 | 0.7573 | 0.8514 | 0.5798 | 0.7837 | 0.6809 | 0.8865 | 0.8571 | 0.7589 |
| SVM | PseKRAAC-type-6A | 0.6577 | 0.2492 | 0.6237 | 0.5791 | 0.6683 | 0.6348 | 0.6037 | 0.6852 | 0.2646 | 0.6312 | 0.5674 | 0.695 | 0.6504 | 0.6061 |
| SVM | PseKRAAC-type-13 | 0.5949 | 0.15 | 0.5743 | 0.5773 | 0.5716 | 0.5756 | 0.5746 | 0.6041 | 0.149 | 0.5745 | 0.5816 | 0.5674 | 0.5734 | 0.5775 |
| SVM | QSOrder | 0.8855 | 0.6445 | 0.8209 | 0.7905 | 0.8513 | 0.842 | 0.8143 | 0.9034 | 0.6882 | 0.844 | 0.8298 | 0.8582 | 0.854 | 0.8417 |
| SVM | TPC-type-1 | 0.3367 | 0.1438 | 0.55 | 0.35 | 0.7536 | 0.5689 | 0.3582 | 0.3671 | 0.2067 | 0.6028 | 0.5532 | 0.6525 | 0.6142 | 0.5821 |
| SVM | TPC-type-2 | 0.1025 | 0.1636 | 0.5402 | 0.2839 | 0.8 | 0.6991 | 0.279 | 0.0721 | 0.3946 | 0.6348 | 0.2695 | 1 | 1 | 0.4246 |

| | | | | | | | | | | | | | | | |
|---|---|---|---|---|---|---|---|---|---|---|---|---|---|---|---|
| XGB | Bepler | 0.9062 | 0.6705 | 0.8333 | 0.8154 | 0.8512 | 0.847 | 0.8284 | 0.9403 | 0.745 | 0.8723 | 0.8582 | 0.8865 | 0.8832 | 0.8705 |
| XGB | CPCProt | 0.8982 | 0.6825 | 0.8397 | 0.8082 | 0.8711 | 0.8643 | 0.834 | 0.9276 | 0.7114 | 0.8546 | 0.8156 | 0.8936 | 0.8846 | 0.8487 |
| XGB | ESM | 0.9172 | 0.7169 | 0.8575 | 0.8314 | 0.8834 | 0.8786 | 0.8537 | 0.9309 | 0.6744 | 0.8369 | 0.8156 | 0.8582 | 0.8519 | 0.8333 |
| XGB | ESM1b | 0.9072 | 0.6816 | 0.8396 | 0.8101 | 0.8691 | 0.8621 | 0.8343 | 0.9161 | 0.6402 | 0.8191 | 0.7801 | 0.8582 | 0.8462 | 0.8118 |
| XGB | ESM1v | 0.9054 | 0.6507 | 0.8244 | 0.7904 | 0.8585 | 0.8484 | 0.8181 | 0.9293 | 0.7133 | 0.8546 | 0.8014 | 0.9078 | 0.8968 | 0.8464 |
| XGB | ESM2-t6 | 0.8856 | 0.6129 | 0.8056 | 0.7849 | 0.8262 | 0.8194 | 0.8008 | 0.9152 | 0.6965 | 0.8475 | 0.8156 | 0.8794 | 0.8712 | 0.8425 |
| XGB | ESM2-t12 | 0.9184 | 0.6733 | 0.836 | 0.8227 | 0.8495 | 0.8448 | 0.8329 | 0.9365 | 0.668 | 0.8333 | 0.8014 | 0.8652 | 0.8561 | 0.8278 |
| XGB | ESM2-t30 | 0.903 | 0.6693 | 0.8334 | 0.8136 | 0.8531 | 0.848 | 0.8289 | 0.932 | 0.7189 | 0.8582 | 0.8156 | 0.9007 | 0.8915 | 0.8519 |
| XGB | ESM2-t33 | 0.9289 | 0.716 | 0.8567 | 0.8317 | 0.8818 | 0.8762 | 0.8521 | 0.9297 | 0.7317 | 0.8652 | 0.8369 | 0.8936 | 0.8872 | 0.8613 |
| XGB | ESM2-t36 | 0.9329 | 0.7293 | 0.862 | 0.8135 | 0.9103 | 0.9026 | 0.8543 | 0.9486 | 0.7661 | 0.883 | 0.8723 | 0.8936 | 0.8913 | 0.8817 |
| XGB | FastText | 0.9132 | 0.6974 | 0.8468 | 0.8083 | 0.8853 | 0.8763 | 0.8396 | 0.8965 | 0.6456 | 0.8227 | 0.8085 | 0.8369 | 0.8321 | 0.8201 |
| XGB | GloVe | 0.8938 | 0.6394 | 0.8191 | 0.8028 | 0.8352 | 0.8302 | 0.8156 | 0.9104 | 0.6249 | 0.8121 | 0.7872 | 0.8369 | 0.8284 | 0.8073 |
| XGB | PRNN | 0.9193 | 0.7025 | 0.8504 | 0.8386 | 0.8621 | 0.8604 | 0.8483 | 0.9355 | 0.6959 | 0.8475 | 0.8227 | 0.8723 | 0.8657 | 0.8436 |

| | | | | | | | | | | | | | | | |
|---|---|---|---|---|---|---|---|---|---|---|---|---|---|---|---|
| XGB | PTAB | 0.9003 | 0.6525 | 0.8243 | 0.8082 | 0.8406 | 0.8391 | 0.8211 | 0.9223 | 0.7021 | 0.8511 | 0.8511 | 0.8511 | 0.8511 | 0.8511 |
| XGB | PTBB | 0.8777 | 0.5918 | 0.7948 | 0.7743 | 0.8156 | 0.8107 | 0.791 | 0.9277 | 0.6538 | 0.8262 | 0.8582 | 0.7943 | 0.8067 | 0.8316 |
| XGB | PTB | 0.9357 | 0.7295 | 0.863 | 0.8245 | 0.9016 | 0.8939 | 0.8568 | 0.935 | 0.6628 | 0.8298 | 0.7801 | 0.8794 | 0.8661 | 0.8209 |
| XGB | PTU | 0.9565 | 0.7763 | 0.887 | 0.8674 | 0.9068 | 0.905 | 0.8848 | 0.9617 | 0.7664 | 0.883 | 0.8652 | 0.9007 | 0.8971 | 0.8809 |
| XGB | PTXU | 0.9528 | 0.7601 | 0.879 | 0.8674 | 0.8908 | 0.89 | 0.8774 | 0.9568 | 0.7806 | 0.8901 | 0.8723 | 0.9078 | 0.9044 | 0.8881 |
| XGB | PTXLU | 0.8635 | 0.5753 | 0.7867 | 0.7831 | 0.7903 | 0.7905 | 0.7854 | 0.8632 | 0.5535 | 0.7766 | 0.7943 | 0.7589 | 0.7671 | 0.7805 |
| XGB | Seq2Vec | 0.9693 | 0.8318 | 0.9148 | 0.8906 | 0.9391 | 0.9373 | 0.9126 | 0.9827 | 0.9008 | 0.9504 | 0.9433 | 0.9574 | 0.9568 | 0.95 |
| XGB | Word2Vec | 0.8789 | 0.6278 | 0.8127 | 0.7831 | 0.8422 | 0.8333 | 0.8065 | 0.9258 | 0.7056 | 0.8511 | 0.8014 | 0.9007 | 0.8898 | 0.8433 |
| XGB | AAC | 0.8914 | 0.6394 | 0.8181 | 0.785 | 0.8512 | 0.8427 | 0.8116 | 0.9261 | 0.7236 | 0.8617 | 0.8723 | 0.8511 | 0.8542 | 0.8632 |
| XGB | ACC | 0.6888 | 0.2914 | 0.6452 | 0.6507 | 0.6398 | 0.6436 | 0.6463 | 0.6477 | 0.2204 | 0.6099 | 0.5745 | 0.6454 | 0.6183 | 0.5956 |
| XGB | AC | 0.6447 | 0.2531 | 0.6264 | 0.6327 | 0.6201 | 0.6256 | 0.6287 | 0.6182 | 0.1994 | 0.5993 | 0.5532 | 0.6454 | 0.6094 | 0.5799 |
| XGB | APAAC | 0.908 | 0.6562 | 0.8262 | 0.7921 | 0.8602 | 0.8514 | 0.819 | 0.909 | 0.6529 | 0.8262 | 0.8085 | 0.844 | 0.8382 | 0.8231 |
| XGB | ASDC | 0.9641 | 0.8242 | 0.9113 | 0.9211 | 0.9014 | 0.9058 | 0.9125 | 0.966 | 0.8537 | 0.9255 | 0.9645 | 0.8865 | 0.8947 | 0.9283 |

| | | | | | | | | | | | | | | | |
|---|---|---|---|---|---|---|---|---|---|---|---|---|---|---|---|
| XGB | CC | 0.6791 | 0.2861 | 0.6425 | 0.6471 | 0.638 | 0.6422 | 0.6436 | 0.6546 | 0.1915 | 0.5957 | 0.5957 | 0.5957 | 0.5957 | 0.5957 |
| XGB | CKSAAGP-type-1 | 0.9324 | 0.7387 | 0.8683 | 0.8799 | 0.8566 | 0.8623 | 0.8699 | 0.9691 | 0.8369 | 0.9184 | 0.922 | 0.9149 | 0.9155 | 0.9187 |
| XGB | CKSAAGP-type-2 | 0.9499 | 0.7727 | 0.8853 | 0.8782 | 0.8925 | 0.8921 | 0.8839 | 0.9809 | 0.8299 | 0.9149 | 0.9078 | 0.922 | 0.9209 | 0.9143 |
| XGB | CKSAAP-type-1 | 0.962 | 0.8149 | 0.9015 | 0.821 | 0.982 | 0.9792 | 0.8917 | 0.9825 | 0.8544 | 0.922 | 0.844 | 1 | 1 | 0.9154 |
| XGB | CKSAAP-type-2 | 0.9609 | 0.8273 | 0.9078 | 0.8318 | 0.9839 | 0.982 | 0.8986 | 0.9794 | 0.8637 | 0.9291 | 0.8723 | 0.9858 | 0.984 | 0.9248 |
| XGB | CTDC | 0.8804 | 0.6423 | 0.8181 | 0.7544 | 0.8818 | 0.8652 | 0.8052 | 0.8909 | 0.6762 | 0.8369 | 0.7943 | 0.8794 | 0.8682 | 0.8296 |
| XGB | CTDD | 0.9877 | 0.9053 | 0.9516 | 0.9212 | 0.9821 | 0.9811 | 0.95 | 0.9919 | 0.9164 | 0.9574 | 0.9291 | 0.9858 | 0.985 | 0.9562 |
| XGB | CTDT | 0.9369 | 0.8007 | 0.8996 | 0.8905 | 0.9087 | 0.9085 | 0.8987 | 0.9295 | 0.766 | 0.883 | 0.8794 | 0.8865 | 0.8857 | 0.8826 |
| XGB | CTriad | 0.9336 | 0.7839 | 0.8862 | 0.81 | 0.9624 | 0.9571 | 0.8752 | 0.9513 | 0.7873 | 0.8901 | 0.8227 | 0.9574 | 0.9508 | 0.8821 |
| XGB | DDE | 0.9643 | 0.848 | 0.9185 | 0.8442 | 0.9929 | 0.9919 | 0.9106 | 0.9837 | 0.8482 | 0.9184 | 0.8369 | 1 | 1 | 0.9112 |
| XGB | DistancePair | 0.8914 | 0.6394 | 0.8181 | 0.785 | 0.8512 | 0.8427 | 0.8116 | 0.9261 | 0.7236 | 0.8617 | 0.8723 | 0.8511 | 0.8542 | 0.8632 |
| XGB | DPC-type-1 | 0.9635 | 0.8444 | 0.9168 | 0.8425 | 0.9911 | 0.99 | 0.9088 | 0.9861 | 0.8607 | 0.9255 | 0.8511 | 1 | 1 | 0.9195 |
| XGB | DPC-type-2 | 0.9669 | 0.8554 | 0.9221 | 0.8461 | 0.9982 | 0.998 | 0.9145 | 0.9814 | 0.8607 | 0.9255 | 0.8511 | 1 | 1 | 0.9195 |

| | | | | | | | | | | | | | | | |
|---|---|---|---|---|---|---|---|---|---|---|---|---|---|---|---|
| XGB | GDPC-type-1 | 0.9447 | 0.7711 | 0.8844 | 0.8889 | 0.88 | 0.8829 | 0.8846 | 0.9724 | 0.8157 | 0.9078 | 0.9149 | 0.9007 | 0.9021 | 0.9085 |
| XGB | GDPC-type-2 | 0.9591 | 0.8223 | 0.9104 | 0.9122 | 0.9087 | 0.9101 | 0.9104 | 0.9907 | 0.9008 | 0.9504 | 0.9433 | 0.9574 | 0.9568 | 0.95 |
| XGB | Geary | 0.6142 | 0.2034 | 0.6013 | 0.5718 | 0.6306 | 0.6086 | 0.5886 | 0.5416 | 0.0285 | 0.5142 | 0.4681 | 0.5603 | 0.5156 | 0.4907 |
| XGB | GTPC-type-1 | 0.8828 | 0.627 | 0.8128 | 0.8047 | 0.8208 | 0.8187 | 0.8108 | 0.9187 | 0.7323 | 0.8652 | 0.8298 | 0.9007 | 0.8931 | 0.8603 |
| XGB | GTPC-type-2 | 0.8942 | 0.6508 | 0.8244 | 0.8154 | 0.8333 | 0.831 | 0.8218 | 0.9563 | 0.7675 | 0.883 | 0.8511 | 0.9149 | 0.9091 | 0.8791 |
| XGB | KSCTriad | 0.928 | 0.7518 | 0.871 | 0.7957 | 0.9462 | 0.9374 | 0.8595 | 0.9454 | 0.7573 | 0.8759 | 0.8156 | 0.9362 | 0.9274 | 0.8679 |
| XGB | Moran | 0.5771 | 0.1422 | 0.5708 | 0.5592 | 0.5825 | 0.5723 | 0.5644 | 0.5028 | 0.0213 | 0.5106 | 0.4894 | 0.5319 | 0.5111 | 0.5 |
| XGB | NMBroto | 0.6409 | 0.2615 | 0.6299 | 0.6128 | 0.6469 | 0.6365 | 0.6228 | 0.5769 | 0.1066 | 0.5532 | 0.5248 | 0.5816 | 0.5564 | 0.5401 |
| XGB | PAAC | 0.9031 | 0.6453 | 0.8217 | 0.8027 | 0.8405 | 0.8358 | 0.8179 | 0.9231 | 0.6744 | 0.8369 | 0.8156 | 0.8582 | 0.8519 | 0.8333 |
| XGB | PDE | 0.9086 | 0.6587 | 0.828 | 0.8009 | 0.8547 | 0.8485 | 0.8226 | 0.9251 | 0.6813 | 0.8404 | 0.8582 | 0.8227 | 0.8288 | 0.8432 |
| XGB | PseKRAAC-type-4 | 0.6141 | 0.1813 | 0.5895 | 0.5915 | 0.5876 | 0.5946 | 0.59 | 0.5596 | 0.0851 | 0.5426 | 0.5319 | 0.5532 | 0.5435 | 0.5376 |
| XGB | PseKRAAC-type-5 | 0.8329 | 0.5907 | 0.7877 | 0.6847 | 0.8907 | 0.8661 | 0.7617 | 0.8494 | 0.5556 | 0.773 | 0.6809 | 0.8652 | 0.8348 | 0.75 |
| XGB | PseKRAAC-type-6A | 0.6523 | 0.2572 | 0.628 | 0.6092 | 0.6468 | 0.634 | 0.62 | 0.6403 | 0.1988 | 0.5993 | 0.5745 | 0.6241 | 0.6045 | 0.5891 |

| | | | | | | | | | | | | | | | |
|---|---|---|---|---|---|---|---|---|---|---|---|---|---|---|---|
| XGB | PseKRAAC-type-13 | 0.5912 | 0.1588 | 0.5779 | 0.5736 | 0.5826 | 0.5858 | 0.5752 | 0.5796 | 0.1136 | 0.5567 | 0.5816 | 0.5319 | 0.5541 | 0.5675 |
| XGB | QSOrder | 0.8972 | 0.6345 | 0.8154 | 0.7868 | 0.8439 | 0.8368 | 0.809 | 0.922 | 0.6738 | 0.8369 | 0.8298 | 0.844 | 0.8417 | 0.8357 |
| XGB | TPC-type-1 | 0.6909 | 0.3786 | 0.664 | 0.4142 | 0.914 | 0.827 | 0.5489 | 0.8125 | 0.5391 | 0.7411 | 0.5177 | 0.9645 | 0.9359 | 0.6667 |
| XGB | TPC-type-2 | 0.7056 | 0.4007 | 0.6765 | 0.4447 | 0.9085 | 0.8348 | 0.5762 | 0.8184 | 0.5777 | 0.773 | 0.6099 | 0.9362 | 0.9053 | 0.7288 |
